# MIDAS: a deep generative model for mosaic integration and knowledge transfer of single-cell multimodal data

**DOI:** 10.1101/2022.12.13.520262

**Authors:** Zhen He, Yaowen Chen, Shuofeng Hu, Sijing An, Junfeng Shi, Runyan Liu, Jiahao Zhou, Guohua Dong, Jinhui Shi, Jiaxin Zhao, Jing Wang, Yuan Zhu, Le Ou-Yang, Xiaochen Bo, Xiaomin Ying

## Abstract

Rapidly developing single-cell multi-omics sequencing technologies generate increasingly large bodies of multimodal data. Integrating multimodal data from different sequencing technologies, *i.e*. mosaic data, permits larger-scale investigation with more modalities and can help to better reveal cellular heterogeneity. However, mosaic integration involves major challenges, particularly regarding modality alignment and batch effect removal. Here we present a deep probabilistic framework for the mosaic integration and knowledge transfer (MIDAS) of single-cell multimodal data. MIDAS simultaneously achieves dimensionality reduction, imputation, and batch correction of mosaic data by employing self-supervised modality alignment and information-theoretic latent disentanglement. We demonstrate its superiority to other methods and reliability by evaluating its performance in full trimodal integration and various mosaic tasks. We also constructed a single-cell trimodal atlas of human peripheral blood mononuclear cells (PBMCs), and tailored transfer learning and reciprocal reference mapping schemes to enable flexible and accurate knowledge transfer from the atlas to new data. Applications in mosaic integration, pseudotime analysis, and cross-tissue knowledge transfer on bone marrow mosaic datasets demonstrate the versatility and superiority of MIDAS.

## 1 Introduction

Recently emerged single-cell multimodal omics (scMulti-omics) sequencing technologies enable the simultaneous detection of multiple modalities, such as RNA expression, protein abundance, and chromatin accessibility, in the same cell [1, 2]. These technologies, including the trimodal DOGMA-seq [3] and TEA-seq [4] and bimodal CITE-seq [5] and ASAP-seq [3], among many others [6–11], reveal not only cellular heterogeneity at multiple molecular layers, enabling more refined identification of cell characteristics, but also connections across omes, providing a systematic view of ome interactions and regulation at single-cell resolution. The involvement of more measured modalities in analyses of biological samples increases the potential for breakthroughs in the understanding of mechanisms underlying numerous processes, including cell functioning, tissue development, and disease occurrence. The growing size of scMulti-omics datasets necessitates the development of new computational tools to integrate massive high-dimensional data generated from different sources, thereby facilitating more comprehensive and reliable downstream analysis for knowledge mining [1, 2, 12]. Such “integrative analysis” also enables the construction of a large-scale single-cell multimodal atlas, which is urgently needed to make full use of publicly available single-cell multimodal data. Such an atlas can serve as an encyclopedia allowing researchers’ transfer of knowledge to their new data and in-house studies [13–15].

Several methods for single-cell multimodal integration have been presented recently. Most of them have been proposed for the integration of bimodal data [15–23]. Fewer trimodal integration methods have been developed; MOFA+ [24] has been proposed for trimodal integration with complete modalities, and GLUE [25] has been developed for the integration of unpaired trimodal data (*i.e*., datasets involving single specific modalities).

All of these current integration methods have difficulty in handling flexible omics combinations. Due to the diversity of scMulti-omics technologies, datasets from different studies often include heterogeneous omics combinations with one or more missing modalities, resulting in a mosaic-like data. The mosaic-like data is increasing rapidly and is predictably prevalent. Mosaic integration methods are urgently needed to markedly expand the scale and modalities of integration, breaking through the modality scalability and cost limitations of existing scMulti-omics sequencing technologies. Most recently, scVAEIT [26], scMoMaT [27], StabMap [28], and Multigrate [29] have been proposed to tackle this problem. However, these methods are not capable of aligning modalities or correcting batches, which results in limited functions and performances. Therefore, flexible and general multimodal mosaic integration remains challenging [30–32]. One major challenge is the reconciliation of modality heterogeneity and technical variation across batches. Another is the achievement of modality imputation and batch correction for downstream analysis.

To overcome these challenges, we developed a probabilistic framework, MIDAS, for the mosaic integration and knowledge transfer of single-cell multimodal data. By employing self-supervised learning [33] and information-theoretic approaches [34], MIDAS simultaneously achieves modality alignment, imputation, and batch correction for single-cell trimodal mosaic data. We further designed transfer learning and reciprocal reference mapping schemes tailored to MIDAS to enable knowledge transfer. Systematic benchmarks and case studies demonstrate that MIDAS can accurately and robustly integrate mosaic datasets. Through the atlas-level mosaic integration of trimodal human peripheral blood mononuclear cell (PBMC) data, MIDAS achieved flexible and accurate knowledge transfer for various types of unimodal and multimodal query datasets. We also applied MIDAS on mosaic datasets of human bone marrow mononuclear cells (BMMCs) and demonstrated the satisfactory performance of MIDAS for mosaic-data-based pseudotime analysis and cross-tissue knowledge transfer.

## 2 Results

### 2.1 MIDAS enables the mosaic integration and knowledge transfer of single-cell multimodal data

MIDAS is a deep generative model [35, 36] that represents the joint distribution of incomplete single-cell multimodal data with Assay for Transposase-Accessible Chromatin (ATAC), RNA, and Antibody-Derived Tags (ADT) measurements. MIDAS assumes that each cell’s multimodal measurements are generated from two modality-agnostic and disentangled latent variables—the biological state (*i.e*., cellular heterogeneity) and technical noise (*i.e*., unwanted variation induced by single-cell experimentation)—through deep neural networks [37]. Its input consists of a mosaic feature-by-cell count matrix comprising different single-cell samples (batches), and a vector representing the cell batch IDs (Fig. 1a). The batches can derive from different experiments or be generated by the application of different sequencing techniques (*e.g*., scRNA-seq [38], CITE-seq [5], ASAP-seq [3], and TEA-seq [4]), and thus can have different technical noise, modalities, and features. The output of MIDAS comprises biological state and technical noise matrices, which are the two low-dimensional representations of different cells, and an imputed and batch-corrected count matrix in which modalities and features missing from the input data are interpolated and batch effects are removed. These outputs can be used for downstream analyses such as clustering, differential expression analysis, cell typing, and trajectory inference [39].

**Figure 1.**
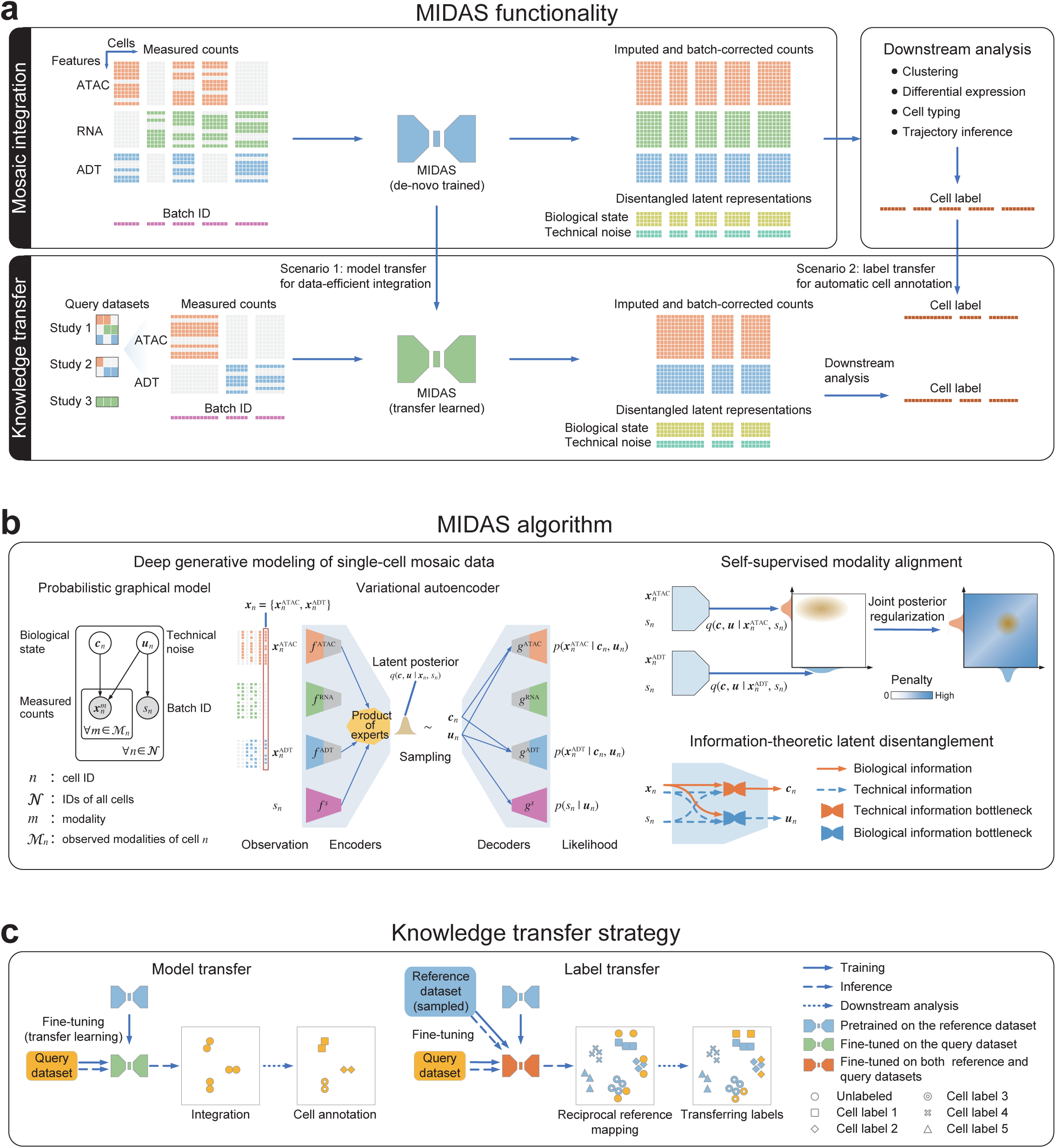
Overview of the MIDAS framework. **a**, Functionality of the MIDAS framework. **b**, MIDAS assumes each cell’s measured counts and batch ID are generated from the biological state and technical noise latent variables, and employs the variational autoencoder to implement model learning and latent variable inference. Self-supervised learning is used to align different modalities on latent space through joint posterior regularization, and information-theoretic approaches help to disentangle the latent variables. **c**, Two strategies are developed for MIDAS to achieve reference-to-query knowledge transfer, where the model transfer utilizes pretrained model for data-efficient integration, and the label transfer reciprocally maps the reference and query datasets onto the latent space for automatic cell annotation.

MIDAS is based on a Variational Autoencoder (VAE) [40] architecture, with a modularized encoder network designed to handle the mosaic input data and infer the latent variables, and a decoder network that uses the latent variables to seed the generative process for the observed data (Fig. 1b). It uses self-supervised learning to align different modalities in latent space, improving cross-modal inference in downstream tasks such as imputation and translation (Fig. 1b, Supplementary Fig. 1). Information-theoretic approaches are applied to disentangle the biological state and technical noise, enabling further batch correction (Fig. 1b). Combining these elements into our optimization objective, scalable learning and inference of MIDAS are simultaneously achieved by the Stochastic Gradient Variational Bayes (SGVB) [41], which also enables large-scale mosaic integration and atlas construction of single-cell multimodal data. For the robust transfer of knowledge from the constructed atlas to query datasets with various modality combinations, transfer learning and reciprocal reference mapping schemes were developed for the transfer of model parameters and cell labels, respectively (Fig. 1c).

### 2.2 MIDAS shows superior performance in trimodal integration with complete modalities

To compare MIDAS with the state-of-the-art methods, we evaluated the performance of MIDAS in trimodal integration with complete modalities, a simplified form of mosaic integration, as few methods are designed specifically for trimodal mosaic integration. We named this task “rectangular integration”. We used two published single-cell trimodal human PBMC datasets (DOGMA-seq [3] and TEA-seq [4], Supplementary Table 1) with simultaneous RNA, ADT, and ATAC measurements for each cell to construct dogma-full and teadog-full datasets. Dogma-full took all four batches (LLL_Ctrl, LLL_Stim, DIG_Ctrl, and DIG_Stim) from the DOGMA-seq dataset, and teadog-full took two batches (W1 and W6) from the TEA-seq dataset and two batches (LLL_Ctrl and DIG_Stim) from the DOGMA-seq dataset (Supplementary Table 2). The integration of each dataset requires the handling of batch effects and missing features and preservation of biological signals, which is challenging, especially for the teadog-full dataset, as the involvement of more datasets amplifies biological and technical variation.

Uniform manifold approximation and projection (UMAP) [42] visualization showed that the biological states of different batches were well aligned and that their grouping was consistent with the cell type labels (Fig. 2a left, Supplementary Fig. 2a left), and that the technical noise was grouped by batch and exhibited little relevance to cell types (Fig. 2b, Supplementary Fig. 2b). Thus, the two inferred latent variables were disentangled well and independently represented biological and technical variation.

**Figure 2.**
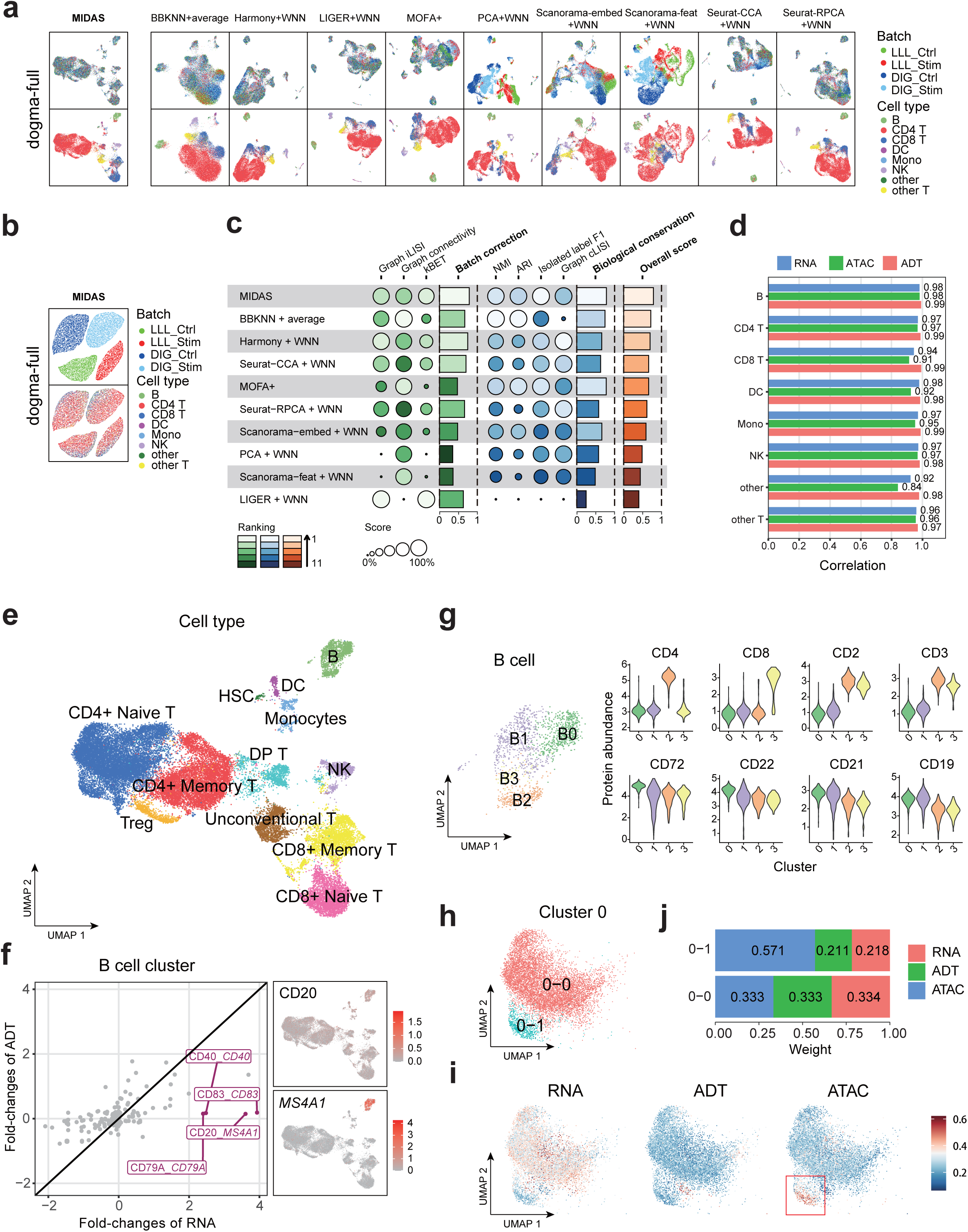
Evaluation and downstream analysis results obtained with MIDAS on rectangular integration tasks. **a**, UMAP visualization of cell embeddings obtained by MIDAS and nine other strategies in the dogma-full dataset. The left two panels show inferred latent biological states and the right panels show dimensionality reduction results obtained with the other strategies. **b**, UMAP visualization of latent technical noise inferred by MIDAS in the dogma-full dataset. **c**, scIB benchmarking of performance on the dogma-full rectangular integration task. **d**, Correlation of fold changes in gene/protein abundance and chromatin accessibility between raw and batch-corrected data. **e**, UMAP visualization of the inferred latent biological states with manually annotated cell types. **f**, Expression inconsistencies between proteins and their corresponding genes in B cells. The left panel shows RNA and ADT fold changes and the right panel shows the UMAP visualization of CD20 and *MS4A1* expression. **g**, UMAP visualization of B-cell subclusters (left) and violin plots of protein abundance across subclusters (right). **h**, UMAP plot of CD4^+^ naive T-cell C0-0 and C0-1 subclusters from the dogma dataset. **i**, Single-cell modality contributions to C0 clustering. The red rectangle highlights the greater contribution of the ATAC modality in cluster C0-1. **j**, Modality contributions to the integrated clustering of C0-0 and C0-1 cells.

Taking the inferred biological states as low-dimensional representations of the integrated data, we compared the performance of MIDAS with that of nine strategies derived from recently published methods (Methods, Supplementary Table 3) in the removal of batch effects and preservation of biological signals. UMAP visualization of the integration results showed that MIDAS ideally removed batch effects and meanwhile preserved cell type information on both dogma-full and teadog-full datasets, whereas the performance of other strategies was not satisfactory. For example, BBKNN+average, MOFA+, PCA+WNN, Scanorama-embed+WNN, and Scanorama-feat+WNN did not mix different batches well, and PCA+WNN and Scanorama-feat+WNN produced cell clusters largely inconsistent with cell types (Fig. 2a, Supplementary Fig. 2a).

In a quantitative evaluation of the low-dimensional representations of different strategies performed with the widely used single-cell integration benchmarking (scIB) [43] tool, MIDAS had the highest batch correction, biological conservation, and overall scores for the dogma-full and teadog-full datasets (Fig. 2c, Supplementary Fig. 2c). In addition, MIDAS preserved cell type-specific patterns in batch-corrected RNA, ADT, and ATAC data (Methods). For each cell type, fold changes in gene/protein abundance and chromatin accessibility in raw and batch-corrected data correlated strongly and positively (all Pearson’s *r >* 0.8; Fig. 2d).

Manual cell clustering and typing based on the integrated low-dimensional representations and batch-corrected data from MIDAS led to the identification of 13 PBMC types, including B cells, T cells, dendritic cells (DCs), natural killer (NK) cells, and monocytes (Fig. 2e). We got a distinct T cell cluster that highly expresses CD4 and CD8 simultaneously. We labeled this cluster as double positive (DP) CD4^+^/CD8^+^ T cells. This phenomenon was also reported in previous studies [44]. Another T cell cluster, containing mucosa-associated invariant T cells and gamma-delta T cells, was distinct from conventional T cells and was labeled as unconventional T cells [45].

As is known, multiple omes regulate biological functions synergistically [1, 2]. MIDAS integrates RNA, ADT and ATAC single-cell data and hence facilitates to discover the intrinsic nature of cell activities in a more comprehensive manner. We found that all omics contributed greatly to the identification of cell types and functions (Supplementary Fig. 3).

Systematic screening for expression inconsistencies between proteins and their corresponding genes, expected to reflect ome irreplaceability, at the RNA and ADT levels demonstrated that several markers in each cell type were expressed strongly in one modality and weakly in the other (Fig. 2f, Supplementary Fig. 4). For instance, *MS4A1*, which encodes a B cell-specific membrane protein, was expressed extremely specifically in B cells, but the CD20 protein encoded by *MS4A1* was rarely detected, confirming the irreplaceability of the RNA modality. We also found that ADT could complement RNA-based clustering. For example, the simultaneous expression of T-cell markers (CD3 and CD4) was unexpectedly observed in two subclusters of B cells (B2 and B3) expressing canonical B-cell makers (CD19, CD21, and CD22; Fig. 2g). As this phenomenon could not be replicated using RNA data alone, this finding confirms the irreplaceability of the ADT modality.

Investigation of the uniqueness of chromatin accessibility in multi-omics integration at the ATAC level showed that ATAC contributed more than did ADT and RNA to the integration of a subcluster of CD4^+^ naive T cells (Methods, Fig. 2h–j). We took the ratio of peak number of a cell to that of all cells as the representation of the cell accessibility level. RNA and ADT expression did not differ between these cells and their CD4^+^ naive T-cell siblings, but surprisingly less accessibility level was observed at the ATAC layer (*<* 0.02, Supplementary Fig. 5). Gene ontology enrichment analysis [46] indicated that the inaccessible regions are related to T-cell activation, cell adhesion, and other immune functions. Therefore, we define this cluster as low chromatin accessible (LCA) naive CD4^+^ T cells. Although this discovery needs to be verified further, it demonstrates the remarkable multi-omics integration capability of MIDAS.

### 2.3 MIDAS enables reliable trimodal mosaic integration

At present, trimodal sequencing techniques are still immature. Most of the existing datasets are unimodal or bimodal with various modality combinations. MIDAS is designed to integrate these diverse multimodal datasets, *i.e*. mosaic datasets. To evaluate the performance of MIDAS on mosaic integration, we further constructed 14 incomplete datasets based on the previously generated rectangular datasets including dogma-full and teadog-full datasets (Methods, Supplementary Table 2). Each mosaic dataset was generated by removing several modality-batch blocks from the full-modality dataset. Then, we took the rectangular integration results as the baseline, and examined whether MIDAS could obtain comparable results on mosaic integration tasks. We assessed MIDAS’s ability of batch correction, modality alignment, and biological conservation. Here we also focused on modality alignment because it guarantees accurate cross-modal inference for processes such as downstream imputation and knowledge transfer. For qualitative evaluation, we use UMAP to visualize the biological states and technical noises inferred from the individual and the joint input modalities (Fig. 3a, b, Supplementary Fig. 6, 7). Taking the dogma-paired-abc dataset for example, for each modality, the biological states were consistently distributed across different batches (Fig. 3a) whereas the technical noises were grouped by batches (Fig. 3b), indicating that the batch effect were well disentangled from the biological states. Similarly, the distributions of biological states and technical noises within batches were very similar across modalities (Fig. 3a, b), suggesting that MIDAS internally aligns different modalities in latent space. Moreover, the biological states of each cell type were grouped together and the cell type silhouettes were consistent across batches and modality combinations (Fig. 3a), reflecting robust conservation of the biological signals after mosaic integration.

**Figure 3.**
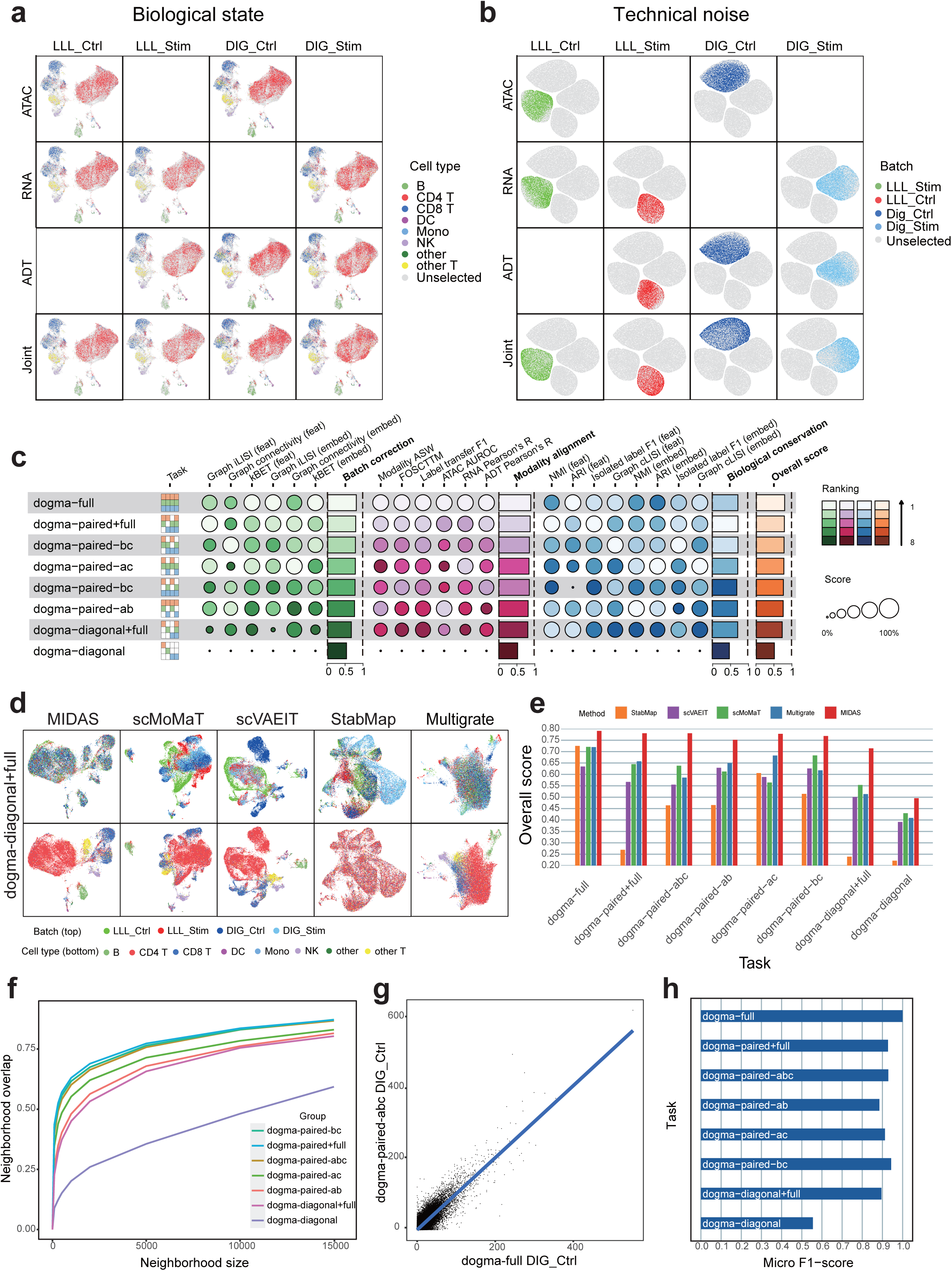
Qualitative and quantitative evaluation of MIDAS performance on mosaic integration tasks. **a, b**, UMAP visualization of the biological states (**a**) and technical noises (**b**) inferred by MIDAS on the dogma-paired-abc dataset. **c**, Benchmarking of MIDAS performance on dogma mosaic integration tasks using our proposed scMIB. **d**, UMAP comparison of embeddings on dogma-diagonal+full mosaic integration tasks. Cells in the top row are colored by batch, and cells in the bottom row are colored by cell type. **e**, Comparison of scIB overall scores on dogma mosaic integration tasks. **f**, Consistency of dimensional reduction results from different tasks with those from the dogma-full task, measured by the overlap of cells’ nearest neighbors. **g**, Consistency of gene regulation links in inferred (dogma-full, DIG_Ctrl batch) and raw (dogma-paired-abc, DIG_Ctrl batch) RNA data. Values represent the regulation importance of gene-transcript factor pairs. **h**, Micro F1-scores reflecting the consistency of downstream-analyzed cell labels between mosaic tasks and the dogma-full task.

To quantitatively evaluate MIDAS on mosaic integration, we proposed single-cell mosaic integration benchmarking (scMIB). scMIB extends scIB with modality alignment metrics, and defines each type of metrics on both embedding (latent) space and feature (observation) space, resulting in 20 metrics in total (Methods, Supplementary Table 4). The obtained batch correction, modality alignment, biological conservation, and overall scores for paired+full, paired-abc, paired-ab, paired-ac, paired-bc, and diagonal+full tasks performed with the dogma and teadog datasets were similar to those obtained with rectangular integration (Fig. 3c, Supplementary Fig. 8a). MIDAS showed moderate performance in the dogma- and teadog-diagonal tasks, likely due to the lack of cell-to-cell correspondence across modalities in these tasks, which can be remedied via knowledge transfer (shown in Result 2.5).

scIB benchmarking showed that MIDAS, when given incomplete datasets (paired+full, paired-abc, paired-ab, paired-ac, and paired-bc for dogma and teadog), outperformed methods that rely on the full-modality datasets (dogma- and teadog-full; Supplementary Fig. 8b, c). Even with the severely incomplete dogma- and teadog-diagonal+full datasets, the performance of MIDAS surpassed that of most the other methods.

We also compared MIDAS against scVAEIT, scMoMaT, Multigrate and StabMap (Methods) that can handle mosaic datasets. UMAP visualization of the low-dimensional cell embeddings showed that MIDAS removed batch effects and preserved biological signals well on various tasks, while the other four methods did not integrate trimodal data well, especially when with missing modalities (dogma in Fig. 3d and Supplementary Fig. 9; teadog in Supplementary Fig. 9). To be specific, MIDAS aligned the cells of different batches well and grouped them consistently with the cell type labels, while the other methods did not mix different batches well and produced cell clusters largely inconsistent with cell types. scIB benchmarking showed that MIDAS had stable performance on different mosaic tasks, and its overall scores were much higher than the other methods (dogma in Fig. 3e; teadog in Supplementary Fig. 10; detailed scores are shown in Supplementary Fig. 11).

The identification of cells’ nearest neighbors based on individual dimensionality reduction results and comparison of neighborhood overlap among tasks showed that this overlap exceeded 0.75 for most tasks, except dogma-diagonal, when the number of neighbors reached 10,000 (Fig. 3f). As imputed omics data has been inferred to deteriorate the accuracy of gene regulatory inference in many cases [47], we evaluated the consistency of downstream analysis results obtained with the performance of different mosaic integration tasks with the dogma datasets. We validated the conservation of gene regulatory networks in the imputed data. In the dogma-paired+full task, for example, the regulatory network predicted from imputed data was consistent with that predicted from the dogma-full data (Fig. 3g). These results indicate that the modality inference performed by MIDAS is reliable.

The MIDAS-based annotation of cell types for the mosaic integration tasks and computation of their confusion matrices and micro-F1 scores showed that the cell type labels generated from the incomplete datasets, except dogma-diagonal, were largely consistent with the dogma-full labels, with all micro F1-scores exceeding 0.885 (Fig. 3h; Supplementary Fig. 12). The separation of monocytes and DCs was difficult in some mosaic experiments, mainly because the latter originate from the former [48] and likely also because the monocyte population in the dogma dataset was small.

To demonstrate the robustness of MIDAS for real-world mosaic integration, we tested MIDAS in more challenging cases including batches with various sequencing depths, batches with inconsistent cell types and perturbations of hyperparameters (Supplementary Note 1; Supplementary Figures 13-15; Supplementary Tables 5, 6). We compared MIDAS with other competing methods on more omics combinations and also benchmarked their computational costs (Supplementary Note 1; Supplementary Figure 16-18). All the results show that MIDAS is a robust, versatile, and efficient tool for single cell multimodal integration.

### 2.4 MIDAS enables the atlas-level mosaic integration of trimodal PBMC data

We used MIDAS for the large-scale mosaic integration of 18 PBMC batches from bimodal sequencing platforms (*e.g*., 10X Multiome, ASAP-seq, and CITE-seq) and the 9 batches from the DOGMA-seq and TEA-seq trimodal datasets (total, 27 batches from 10 platforms comprising 185,518 cells; Methods, Supplementary Table 1, 9). Similar to the results obtained with the dogma-full and teadog-full datasets, MIDAS achieved satisfactory batch removal and biological conservation. UMAP visualization showed that the inferred biological states of different batches maintained a consistent PBMC population structure and conserved batch-specific (due mainly to differences in experimental design) biological information (Fig. 4a; Supplementary Fig. 19, 20a). In addition, the technical noise was clearly grouped by batch (Supplementary Fig. 20b). These results suggest that the biological states and technical noises were disentangled well and the data could be used reliably in downstream analysis.

**Figure 4.**
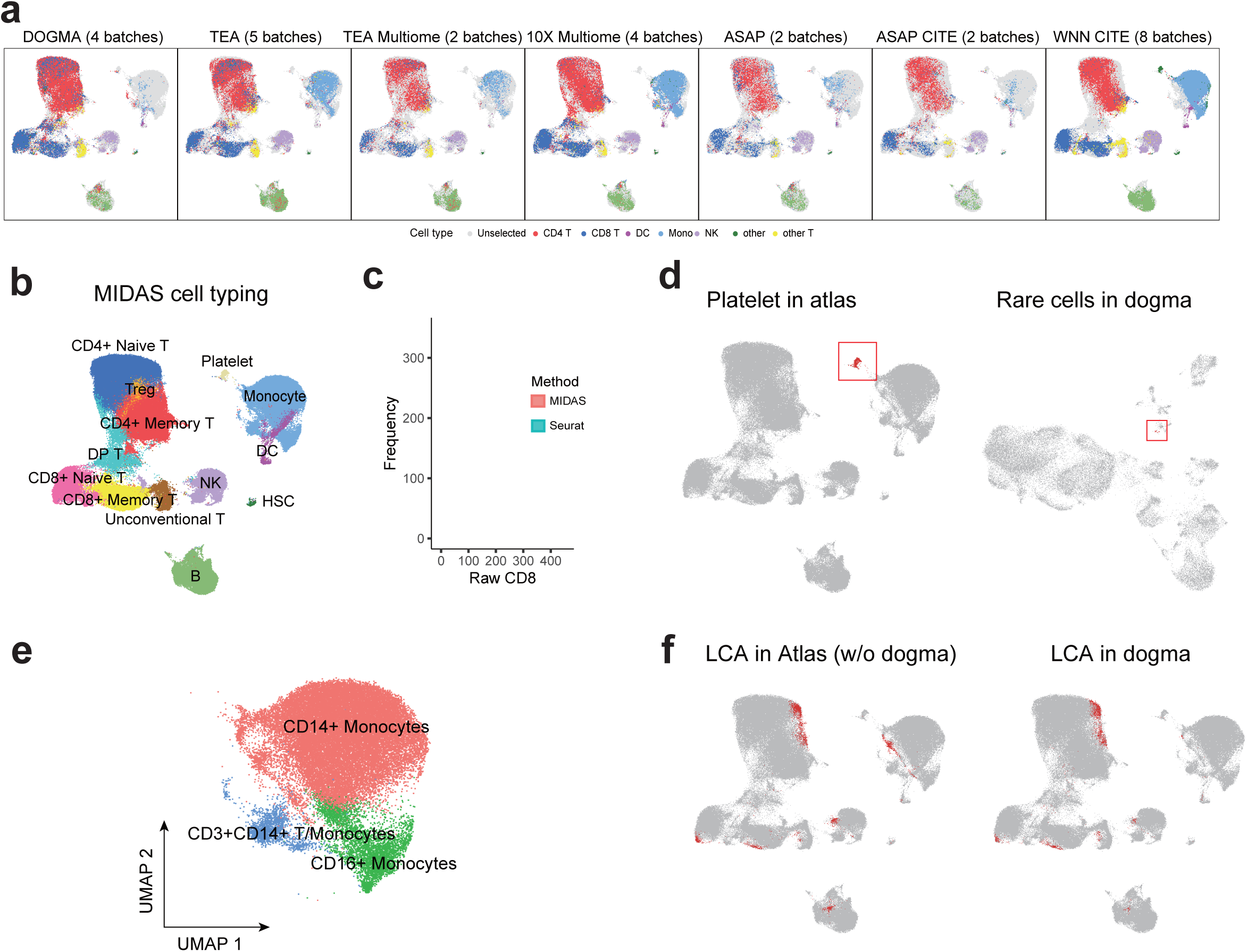
Atlas-level mosaic integration and downstream analysis results obtained with the application of MIDAS to trimodal PBMC data. **a**, UMAP visualization of the biological states inferred by MIDAS with the PBMC mosaic dataset across 7 datasets. Cell type labels are derived from Seurat labeling. **b**, Labels of atlas cells annotated based on clustering of MIDAS embeddings. **c**, The distributions of raw protein levels of CD8 among CD8+ cells labeled by MIDAS and by Seurat, respectively, from the DIG_Ctrl batch of the DOGMA-seq dataset. **d**, Platelets in the atlas (left) and a rare cluster of platelet-like cells in dogma (right) datasets. **e**, High resolution of monocyte types in the atlas. **f**, Cells with low chromatin accessibility (LCA; red) in the atlas. Left, all LCA cells except those from the dogma dataset; right, LCA cells from the dogma dataset.

The manual labeling of cell types according to cluster markers achieved largely consistent separation and annotation with the automatic labeling by Seurat, which indicates the reliability of MIDAS for constructing the atlas [15] (Fig. 4b; Supplementary Fig. 20a). We also found that the MIDAS labeling seems more biologically meaningful when we checked the CD8 protein level of the CD8-labeled cells between the two labeling systems (MIDAS, Seurat; Figure 4c). Consistent with the rectangular integration results (Fig. 2e), we identified all cell types known to be in the atlas, including B cells, conventional T-cell subsets, DP T cells, NK cells, unconventional T cells, and hematopoietic stem cells (HSCs), demonstrating the robustness of MIDAS. Remarkably, the integration of more datasets with MIDAS led to the identification of rare clusters and high-resolution cell typing. For example, a group of cells from the DOGMA-seq dataset aggregated into a much larger cluster with recognizable platelet markers in the PBMC atlas (Fig. 4d). Since platelets have no cell nucleus and are not expected to be present in DOGMA-seq dataset, this rare group of cells could motivate researchers to perform further experiments to validate it. In addition, the atlas contained more monocyte subclusters, including CD14^+^, CD16^+^, and CD3^+^CD14^+^ monocytes, than obtained with rectangular integration (Fig. 4e). Other cell types present in more subclusters in the atlas included CD158e1^+^ NK cells, CD4^+^CD138^+^CD202b^+^ T cells, and RTKN2^+^CD8^+^ T cells (Supplementary Fig. 21a).

Most batches in the atlas contained considerable numbers of LCA cells (Fig. 4f; Supplementary Fig. 21b) with *<* 0.02 accessibility level, as did the DOGMA-seq dataset (Fig. 2i). The chromatin accessibility levels of cells in the atlas showed an obvious bimodal distribution, reflecting the existence of two ATAC patterns (Supplementary Fig. 21b). CD8^+^ T-cell, CD14^+^ monocyte, NK cell, B cell, and other clusters contained LCA cells (Fig. 4b, f) implying that LCA is common in various cell types.

### 2.5 MIDAS enables flexible and accurate knowledge transfer across modalities and datasets

To investigate the knowledge transfer capability of MIDAS, we re-partitioned the atlas dataset into reference (for atlas construction) and query (knowledge transfer target) datasets (Supplementary Table 9). By removing DOGMA-seq from the atlas, we obtained a reference dataset named atlas-no_dogma. To test the flexibility of knowledge transfer, we used DOGMA-seq to construct 14 query datasets: 1 rectangular integration and 7 incomplete mosaic datasets generated previously, and 6 rectangular integration datasets with fewer modalities (Methods, Supplementary Table 10). In consideration of real applications, we defined model and label knowledge transfer scenarios (Methods). In the model transfer scenario, knowledge was transferred implicitly through model parameters via transfer learning. In the label transfer scenario, knowledge was transferred explicitly through cell labels via reference mapping.

We assessed the performance of MIDAS in the model transfer scenario. For the transfer learned models, we used UMAP to visualize the inferred biological states and technical noises and scMIB and scIB for integration benchmarking, and compared the results of different tasks with those generated by *de novo* trained models. Transfer learning greatly improved performance on the dogma-diagonal, dogma-atac, dogma-rna, and dogma-paired-a tasks, with performance levels on the other tasks maintained (Fig. 5a-c; Supplementary Fig. 22, 23). For example, the *de novo* trained model failed to integrate well in the dogma-diagonal task due to lack of cell-to-cell correspondence across modalities (Fig. 5a), whereas the transfer learned model with atlas knowledge successfully aligned the biological states across batches and modalities and formed groups consistent with cell types (Fig. 5b). The results obtained by transfer learned models with all 14 datasets were not only comparable (Supplementary Fig. 23a, b), but even superior to those of many other methods that use the complete dataset. (Fig. 5c; Supplementary Fig. 23b).

**Figure 5.**
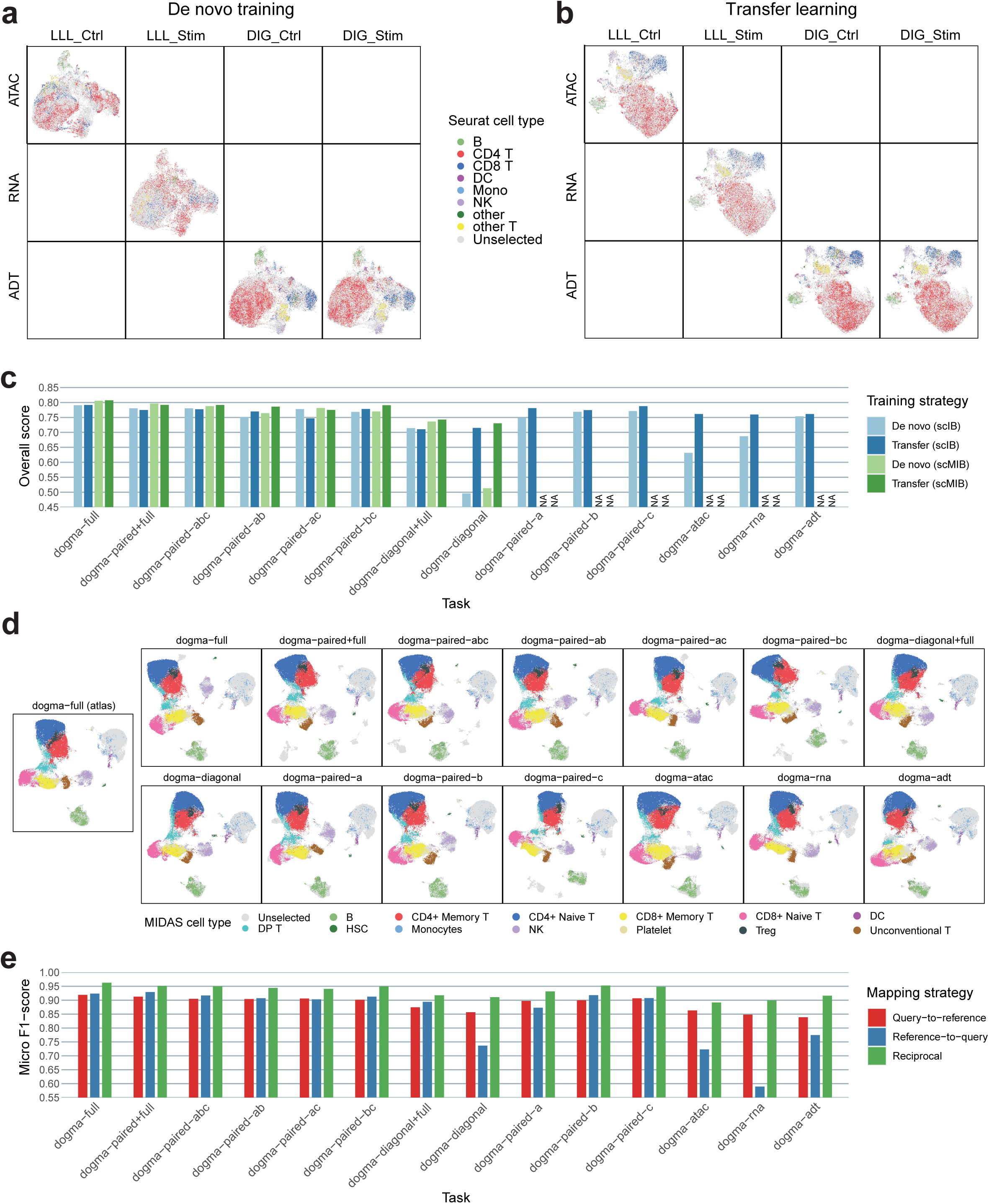
Qualitative and quantitative evaluation of MIDAS on knowledge transfer tasks. **a, b**, UMAP visualization of the biological states inferred by the *de novo* trained (**a**) and transfer-learned (**b**) MIDAS on the dogma-diagonal dataset. **c**, Overall scIB and scMIB scores reflecting transfer-learned and *de novo* trained MIDAS performance on 14 dogma mosaic integration tasks. **d**, UMAP visualization of the biological states obtained by reciprocal reference mapping with 14 dogma mosaic datasets (columns 2–8). Column 1 shows the dogma-full atlas integration. **e**, Label transfer micro F1-scores representing performance on 14 dogma mosaic query datasets.

To assess the performance of MIDAS in the label transfer scenario, we compared the widely used query-to-reference mapping [49, 50], reference-to-query mapping [14, 51], and our proposed reciprocal reference mapping (Methods).

For each strategy, we aligned each query dataset to the reference dataset and transferred cell type labels through the k-Nearest Neighbors (kNN) algorithm, where the ground-truth cell type labels were taken from the trimodal PBMC atlas annotated by MIDAS. Visualization of the mapped biological states showed that reciprocal reference mapping with different query datasets yielded consistent results, with strong agreement with the atlas integration results obtained with the dogma-full dataset (Fig. 5d, Supplementary Fig. 24). Micro F1-scores indicated that reciprocal reference mapping outperformed the query-to-reference and reference-to-query mapping strategies for various forms of query data, achieving robust and accurate label transfer and thereby avoiding the need for *de novo* integration and downstream analysis (Fig. 5e).

Thus, MIDAS can be used to transfer atlas-level knowledge to various forms of users’ datasets without expensive *de novo* training or complex downstream analysis.

### 2.6 Application of MIDAS on BMMC mosaic data

To investigate the application of MIDAS in single-cell datasets with continuous cellular state changes, we constructed a human BMMC mosaic dataset, denoted as “bm”, by combining three distinct batches (ICA, ASAP, CITE) obtained from publicly available datasets of scRNA-seq, ASAP-seq, and CITE-seq, respectively (Methods). The results of *de novo* integration on bm showed that MIDAS accurately aligned different modalities and removed batch effects while preserving cell type information (Supplementary Fig. 25a). Through comparison, we found that MIDAS outperformed the other trimodal mosaic integration methods in both qualitative (Supplementary Fig. 25b) and quantitative (Supplementary Fig. 25c) results.

Next, we performed a pseudotime analysis of the myeloid cells based on the 32-dimensional latent variables generated by MIDAS (Figure. 6a, b). The results showed that hematopoietic stem cells (HSC) (marked by *CD34, SPINK2*) mainly differentiate into two branches. One branch corresponds to the precursor of megakaryocytes and erythrocytes (marked by *GYPA, ASHP*), and the other branch differentiates into granulocyte-macrophage progenitors (GMP) through lymphoid-primed multi-potential progenitors (LMPP), and finally differentiates into monocytes and DC cells (marked by *TYROBP, CTSS*) (Figure. 6c). This differentiation trajectory is consistent with the well-known developmental pathways of myeloid cells in bone marrow [52], demonstrating that MIDAS’s 32-dimensional biological state latent variables can be applied to trajectory inference of cell differentiation. It is worth noting that the original data cannot be directly used for pseudotime analysis since one batch lacks RNA modality.

**Figure 6.**
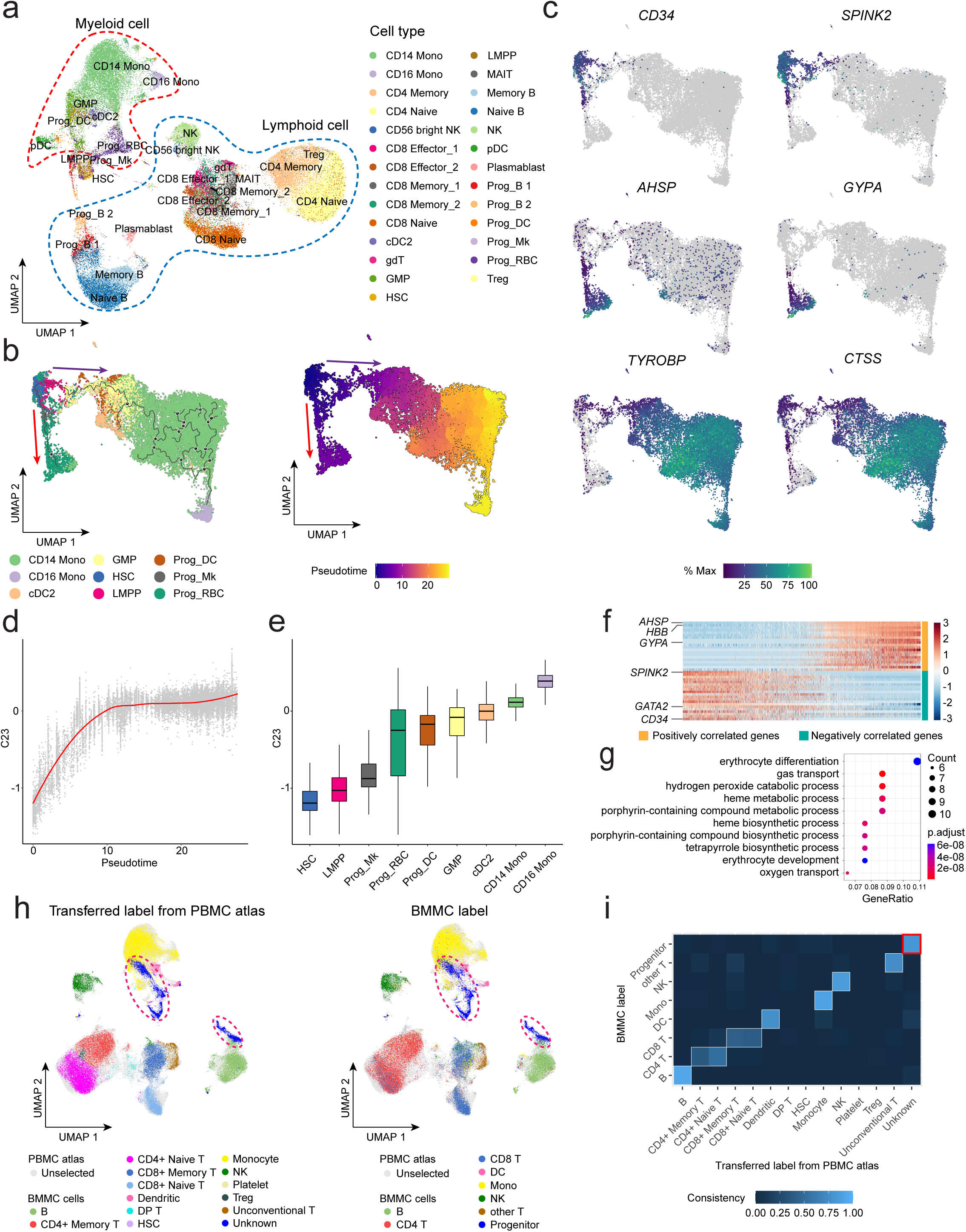
Application of MIDAS on BMMC mosaic dataset. **a**, UMAP visualization of BMMC dataset labeled by Seurat. **b**, UMAP visualization of the inferred trajectory and pseudotime based on 32-dimensional biological state latent representation in myeloid cells. **c**, UMAP visualization colored by gene expression of key cell-type markers. **d**, Loess smoothed curve showing trends of C23 along with the pseudotime. **e**, Boxplots of C23 values in each cell type sorted by the medians. **f**, Heatmap showing scaled expression of the top 20 positively and negatively correlated genes of C23. **g**, Dotplot showing the top 10 significantly enriched GO BP terms of the positively correlated genes of C23 with clusterProfiler. **h**, UMAP visualization of the biological states obtained by cross-tissue reciprocal reference mapping between PBMC atlas reference and BMMC query dataset. The BMMC cells are colored by cell types transferred from PBMC atlas using MIDAS (left panel), and are colored by cell types annotated with Seurat (right panel). **i**, Confusion plot showing the label transfer consistency in a cross-tissue label transfer task on the BMMC mosaic dataset.

We further explored the possible biological meanings of each dimension in the latent space. Notably, latent dimension 23 (C23) corresponded to the pseudotime inferred by monocle3 [53] and captured a gradual transition from HSC to progenitor cells and finally to mature cells (Figure. 6d, e). These results suggested that C23 summarized a gene program contributing to cell development and differentiation. We further calculated correlations between C23 and all genes within the megakaryocyte and erythrocyte developing branch. The negatively correlated genes included the canonical HSC markers, such as *SPINK2, CD34* and *GATA2* (Figure. 6f). The positive correlated genes included many erythrocyte-related genes, such as *AHSP, HBB* and *GYPA* that are demonstated to be involved in erythrocyte differentiation and function by clusterProfiler [54] (Figure. 6g). These results showcase the ability of MIDAS’ biological state latent variables to capture meaningful biological information.

To showcase the capability of MIDAS in cross-tissue knowledge transfer, we employed our constructed PBMC atlas as the reference dataset and bm as the query dataset. Firstly, we performed experiments on model transfer and found that it yielded comparable qualitative and quantitative performance to *de novo* integration (Supplementary Fig. 26), while taking less than half the time (model transfer: 1.28 hours, *de novo* integration: 2.61 hours). Subsequently, we conducted experiments on label transfer, which showed that MIDAS successfully transferred cell type labels from the PBMC atlas reference to the bm query dataset (Fig. 6h). Remarkably, it accurately identified an unknown cell type in the query dataset, which turned out to be the progenitor cell not present in the reference dataset (Fig. 6h, i).

## 3 Discussion

By modeling the single-cell mosaic data generative process, MIDAS can precisely disentangle biological states and technical noises from the input and robustly align modalities to support multi-source and heterogeneous integration analysis. It provides accurate and robust results and outperforms other methods when performing various mosaic integration tasks. It also powerfully and reliably integrates large datasets, as demonstrated with the atlas-scale integration of publicly available PBMC multi-omics datasets. Moreover, MIDAS efficiently and flexibly transfers knowledge from reference to query datasets, enabling convenient handling of new multi-omics data. With superior performance in dimensionality reduction and batch correction, MIDAS supports accurate downstream biological analysis. In addition to enabling clustering, differential expression analysis and cell type identification for mosaic data, MIDAS can also assist in pseudotime analysis for cells with continuous states, which will be especially helpful when no RNA omics is available. When transferring knowledge between different tissues, MIDAS is capable of aligning heterogeneous datasets and identifying cell types and even new types.

To our knowledge, MIDAS is the first model that supports simultaneous dimensionality reduction, modality complementing, and batch correction in single-cell trimodal mosaic integration. MIDAS accurately integrates mosaic data with missing modalities, achieving results comparable to the rectangular integration and superior to those obtained from other methods. These distinct advantages of MIDAS derive from the deep generative modeling, product of experts, information-theoretic disentanglement, and self-supervised modality alignment components of the algorithm, which are specifically designed and inherently competent for the heterogeneous integration of data with missing features and modalities. In addition, MIDAS is the first model that allows knowledge transfer across mosaic data modalities, batches, and even tissues in a highly flexible and reliable manner, enabling researchers to conquer the vast bodies of data produced with continuously emerging multi-omics techniques.

GLUE, designed for the integration of unpaired single-cell trimodal data, aligns the global distributions of different modalities with the use of prior information to improve cell-wise correspondence. It may face challenges when little prior information about the inter-modality (*e.g*., ATAC *vs*. ADT) correspondence of single cells is available. In addition, it is not designed to utilize paired data to enhance modality alignment. With the widespread adoption and rapid development of scMulti-omics sequencing technologies, paired data are rapidly becoming more common and will soon be ubiquitous. By leveraging paired cell data, MIDAS can learn to better align different modalities in a self-supervised manner and can thus play a more important role than other available methods in the blooming scMulti-omics era.

Most recently, scVAEIT, scMoMaT, StabMap and Multigrate were proposed for mosaic integration. However, they do not have the functionalities of modality alignment and batch correction. StabMap also requires to manually select paired datasets as reference sets in advance, leading to biased results. A general mosaic integration method should allow input of diverse mosaic combinations, and support modality alignment and batch correction both in embedding space and feature space [31]. These functions are all essential and urgently needed in real-world scenarios. Compared to scVAEIT, scMoMaT, StabMap and Multigrate, MIDAS is the only one that tackles the problem of general mosaic integration (Supplementary Table 11).

Recently proposed methods for scalable reference-to-query knowledge transfer for single-cell multimodal data have issues with generalization to unseen query data [49, 50] or the retention of information learned on the reference data [14, 51], which make the alignment of reference and query datasets difficult. In addition, they support limited numbers of specific modalities. The MIDAS knowledge transfer scheme stands out from these methods because it supports various types of mosaic query dataset and enables model transfer for sample-efficient mosaic integration and label transfer for automatic cell annotation. Moreover, the generalization and retention problems are mitigated through a novel reciprocal reference mapping scheme.

We envision two major developmental directions for MIDAS. At present, MIDAS integrates only three modalities. By fine tuning the model architecture, we can achieve the integration of four or more modalities, overcoming the limitations of existing scMulti-omics sequencing technologies. In addition, the continuous incorporation of rapidly increasing bodies of newly generated scMulti-omics data is needed to update the model and improve the quality of the atlas. This process requires frequent model retraining, which is computationally expensive and time consuming. Thus, employing incremental learning [55] is an inevitable trend to achieve continuous mosaic integration without model retraining.

## 4 Methods

### 4.1 Deep generative modeling of mosaic single-cell multimodal data

For Cell *n* ∈ *N* = {1, …, *N*} with batch ID *s*_*n*_ ∈ *B* = {1, …, *B*}, let 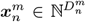 be the count vector of size 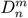 from Modality *m*, and 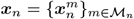 the set of count vectors from the measured modalities *M*_*n*_ ⊆ *M* = {ATAC, RNA, ADT}. We define two modality-agnostic low-dimensional latent variables 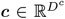 and 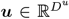 to represent each cell’s biological state and technical noise, respectively. To model the generative process of the observed variables ***x*** and *s* for each cell, we factorize the joint distribution of all variables as below:

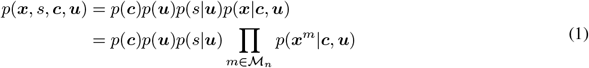

where we assume that *c* and *u* are independent of each other and the batch ID *s* only depends on *u* in order to facilitate the disentanglement of both latent variables, and that the count variables 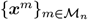 from different modalities are conditional independent given *c* and *u*.

Based on the above factorization, we define a generative model for ***x*** and *s* as follows:

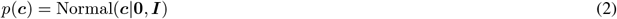

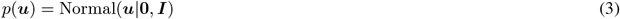

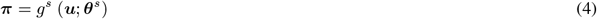

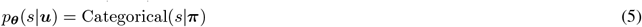

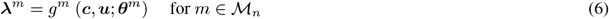

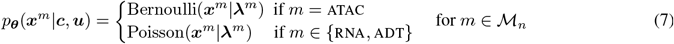

where the priors *p*(***c***) and *p*(***u***) are set as standard Gaussians. The likelihood *p*_***θ***_(*s*|***u***) is set as a categorical distribution with probability vector ***π*** *∈* Δ^*B−*1^ generated through a batch-ID decoder *g*^*s*^ which is a neural network with learnable parameters ***θ***^*s*^. The likelihood *p*_***θ***_(***x***^*m*^|***c, u***) is set as a Bernoulli distribution with mean 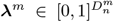 when *m* = ATAC, and as a Poisson distribution with mean 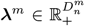 when *m ∈* {RNA, ADT}, where ***λ***^*m*^ is generated through a modality decoder neural network *g*^*m*^ parameterized by ***θ***^*m*^. To mitigate overfitting and improve generalization, we share parameters of the first few layers of different modality decoders {*g*^*m*^}_*m* ∈ *ℳ*_ (the gray parts of the decoders in Fig. 1b middle). The corresponding graphical model is shown in Fig. 1b (left).

Given the observed data {***x***_*n*_, *s*_*n*_}_*n∈ 𝒩*_, we aim to fit the model parameters ***θ*** = {***θ***^*s*^, {***θ***^*m*^}_*m* ∈ *ℳ*_} and meanwhile infer the posteriors of latent variables {***c, u***} for each cell. This can be achieved by using the SGVB [41] which maximizes the expected Evidence Lower Bound (ELBO) for individual datapoints. The ELBO for each individual datapoint {***x***_*n*_, *s*_*n*_} can be written as:

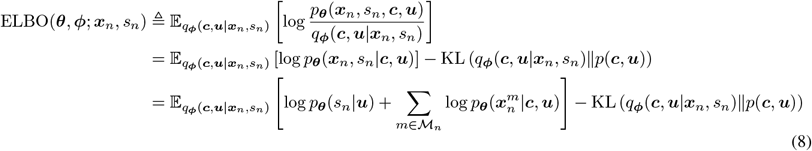

where *q*_***ϕ***_(***c, u***|***x***_*n*_, *s*_*n*_), with learnable parameters ***ϕ***, is the variational approximation of the true posterior *p*(***c, u***|***x***_*n*_, *s*_*n*_) and is typically implemented by neural networks, and KL(·∥·) is the Kullback–Leibler divergence (KLD) between two distributions.

### 4.2 Scalable variational inference via the Product of Experts

Let *M* = | ℳ| be the total modality number, since there are (2^*M*^ *−* 1) possible modality combinations for the count data 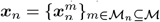, naively implementing *q*_***ϕ***_(***c, u***|***x***_*n*_, *s*_*n*_) in Eq. 8 requires (2^*M*^ *−* 1) different neural networks to handle different cases of input (***x***_*n*_, *s*_*n*_), making inference unscalable. Let ***z*** = {***c, u***}, inspired by [56] which utilizes the Product of Experts (PoEs) to implement variational inference in a combinatorial way, we factorize the posterior *p*(***z***|***x***_*n*_, *s*_*n*_) and define its variational approximation *q*_***ϕ***_(***z***|***x***_*n*_, *s*_*n*_) as follows:

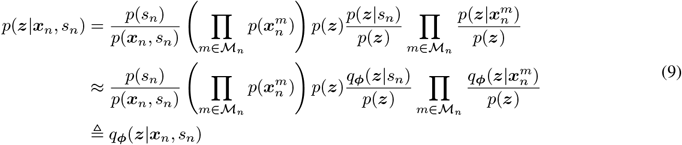

where *q*_***ϕ***_(***z***|*s*_*n*_) and 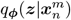 are the variational approximations of the true posteriors *p*(***z***|*s*_*n*_) and 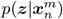, respectively (see Supplementary Note x for detailed derivation). Let 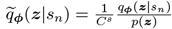 and 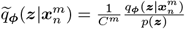 be the normalized quotients of distributions with normalizing constants *C*^*s*^ and *C*^*m*^, respectively. From Eq. 9 we further get:

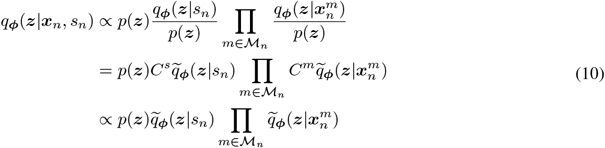

where we set *q*_***ϕ***_(***z***|*s*_*n*_) and 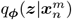 to be diagonal Gaussians, resulting in 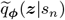 and 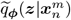 being diagonal Gaussians, which are defined as follows:

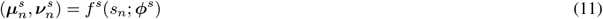

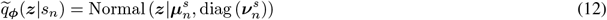

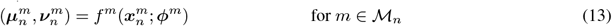

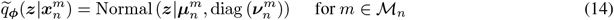

where *f*^*s*^, with parameters ***ϕ***^*s*^, is the batch-ID encoder neural network for generating the mean 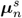 and variance 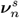 of 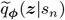, and *f*^*m*^, with parameters ***ϕ***^*m*^, is the modality encoder neural network for generating the mean 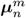 and variance 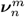 of 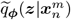. The operator diag(·) converts a vector into a diagonal matrix.

In Eq. 10, since *q*_***ϕ***_(***z***|***x***_*n*_, *s*_*n*_) is proportional to the product of individual Gaussians (or “experts”), itself is a Gaussian, whose mean ***μ***_*n*_ and variance ***v***_*n*_ can be calculated using those of the individual Gaussians:

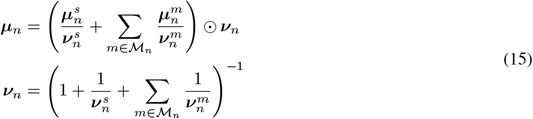

where ⊙ is the Hadamard product.

In doing this, *q*_***ϕ***_(***z***|***x***_*n*_, *s*_*n*_) is modularized into (*M* + 1) neural networks to handle (2^*M*^ *−* 1) different modality combinations, increasing the model’s scalability. Similar to the modality decoders, we also share parameters of the last few layers of different modality encoders {*f*^*m*^}_*m* ∈ *ℳ*_ (the gray parts of the encoders in Fig. 1b middle) to improve generalization.

### 4.3 Handling missing features via padding and masking

For each modality, as different cells can have different feature sets (*e.g*., genes for RNA modality), it is hard to use a fixed-size neural network to handle these cells. To remedy this, we first convert 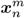 of variable size into a fixed size vector for inference. For Modality *m*, let 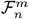 be the features of cell *n*, and 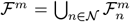 the feature union of all cells. The missing features of cell *n* can then be defined as 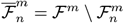. We pad 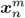 of size 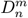 with zeros corresponding to its missing features 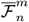 through a zero-padding function *h*:

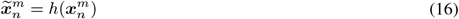

where 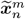 is the zero-padded count vector of constant size *D*^*m*^ = |*ℱ*^*m*^|. The modality encoding process is thus decomposed as:

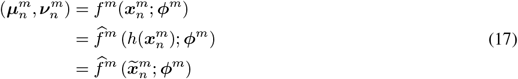

where 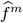 is the latter part of the modality encoder to handle a fixed size input 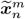. On the other hand, given the sampled latent variables {***c***_*n*_, ***u***_*n*_}, to calculate the likelihood 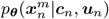 we also need to generate a mean 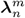 of variable size for 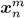. To achieve this, we decompose the modality decoding process as follows:

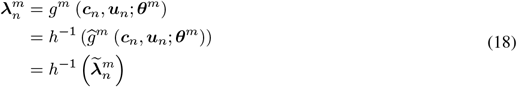

where 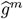 is the front part of the modality decoder to generate the mean 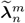 of fixed size *D*^*m*^, and *h*^*−*1^, the inverse function of *h*, is the mask function to remove the padded missing features 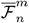 from 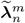 to generate 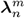. Note that 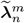 can also be taken as the imputed values for downstream analyses (Methods 4.8 and 4.9).

### 4.4 Self-supervised modality alignment

To achieve cross-modal inference in downstream tasks, we resort to aligning different modalities in the latent space. Leveraging self-supervised learning, we first use each cell’s multimodal observation 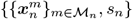 to construct unimodal observations 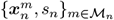, each of which is associated with the latent variables ***z***^*m*^ = {***c***^*m*^, ***u***^*m*^}. Then, we construct a pretext task, which enforces modality alignment by regularizing on the joint space of unimodal variational posteriors with the dispersion of latent variables as a penalty (Fig. 1b upper right), corresponding to a modality alignment loss:

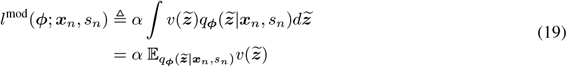

where *α >* 0 is the loss weight, 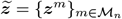 is the set of latent variables, and 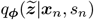 represents the joint distribution of unimodal variational posteriors since:

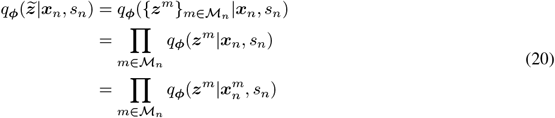

In Eq. 19, 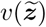 is the Sum of Squared Deviations, which measures the dispersion among different elements in 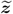 and is used to regula rize 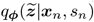. It is defined as:

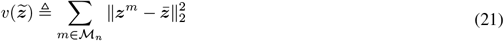

where 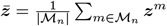 is the mean and ‖ · ‖_2_ the Euclidean distance.

Note that the computation of 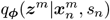 in Eq. 20 is efficient. Since 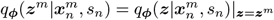, according to Eq. 10 we have:

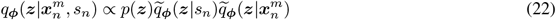

As the mean and covariance of each Gaussian term on the right-hand side of Eq. 22 was already obtained when inferring *q*_***ϕ***_(***z***|***x***_*n*_, *s*_*n*_) (Eq. 10), the mean and covariance of 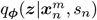 can be directly calculated using Eq. 15, avoiding the need of passing each constructed unimodal observation to the encoders.

### 4.5 Information-theoretic disentanglement of latent variables

To better disentangle the biological state ***c*** and the technical noise ***u***, we adopt an information-theoretic approach, the Information Bottleneck (IB) [34], to control the information flow during inference. We define two types of IB, where the technical IB prevents batch-specific information being encoded into ***c*** by minimizing the Mutual Information (MI) between *s* and ***c***, and the biological IB prevents biological information being encoded into ***u*** by minimizing the MI between ***x*** and ***u*** (Fig. 1b bottom right). Let I(·, ·) denote the MI between two variables, we implement both IBs by minimizing the weighted sum of I(*s*, ***c***) and I(***x, u***):

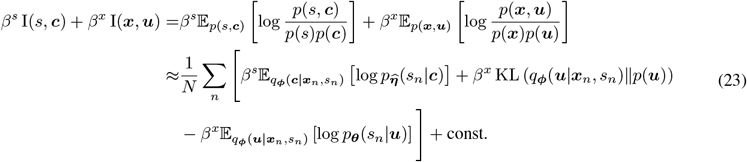

where *β*^*s*^, *β*^*x*^ *>* 0 are the weights, and 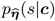 is a learned likelihood with parameters 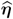 (see Supplementary Note x for detailed derivation). Minimizing (*β*^*s*^*I*(*s*, ***c***) + *β*^*x*^*I*(***x, u***)) thus approximately equals to minimizing the IB loss 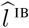 w.r.t ***ϕ*** for all cells:

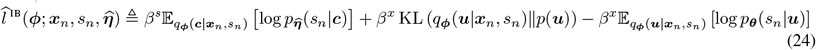

For 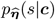, we model it as 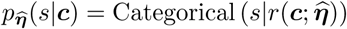, where *r* is a classifier neural network parameterized by 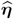.To learn the classifier, we minimize the following expected negative log-likelihood w.r.t 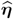:

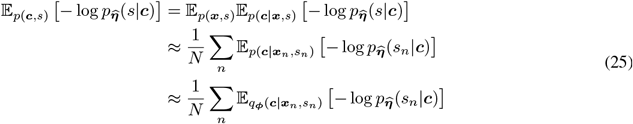

from which we can define a classifier loss:

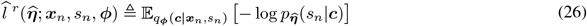

To further enhance latent disentanglement for cross-modal inference, we also apply IBs on the data generated from our self-supervised tasks, *i.e*., minimizing (*β*^*s*^*I*(*s*, ***c***^*m*^) + *β*^*x*^*I*(***x***^*m*^, ***u***^*m*^)) for each modality *m*. Similar to Eqs. 23 and 24, this can be achieved by minimizing 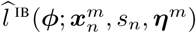 where ***η***^*m*^ is the parameters of the classifier neural network *r*^*m*^ to generate the likelihood 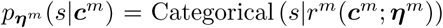. Together with the IB loss of Eq. 24, the total IB loss is defined as:

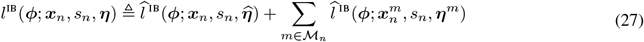

where 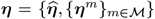. To learn 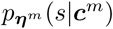, we can also minimize 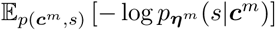, which corresponds to minimizing 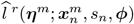 according to Eqs. 25 and 26. With the classifier loss of Eq. 26, we define the total classifier loss as:

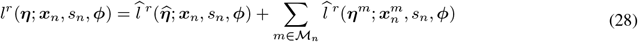

### 4.6 Training MIDAS

To train the encoders and decoders of MIDAS, considering the training objectives defined in Eqs. 8, 19, and 27, we minimize the following objective w.r.t {***θ, ϕ***} for all observations {***x***_*n*_, *s*_*n*_}_*n∈ 𝒩*_ :

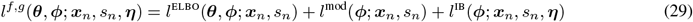

Here the loss *l*^ELBO^ is defined based on the negative of the ELBO of Eq. 8, *i.e*.:

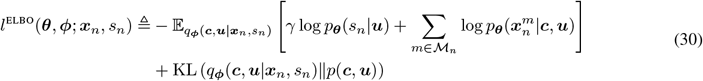

where *γ ≥* 1 is an additional weight that can be set to a higher value to encourage ***u*** to encode more batch-specific information. In Eq. 29, since the classifier parameters ***η*** are unknown and the learning of ***η*** depends on ***ϕ*** as in Eq. 28, we iteratively minimize Eqs. 28 and 29 with Stochastic Gradient Descent (SGD), forming the MIDAS training algorithm (Algorithm 1). In order to better guide the optimization of the IB loss in Eq. 29 for disentangling latent variables, we increase the number of updates (*i.e*., with *K >* 1) of Eq. 28 for the classifier parameters ***η*** in each iteration to ensure that these classifiers stay close to their optimal solutions.

#### Algorithm 1

The MIDAS training algorithm.

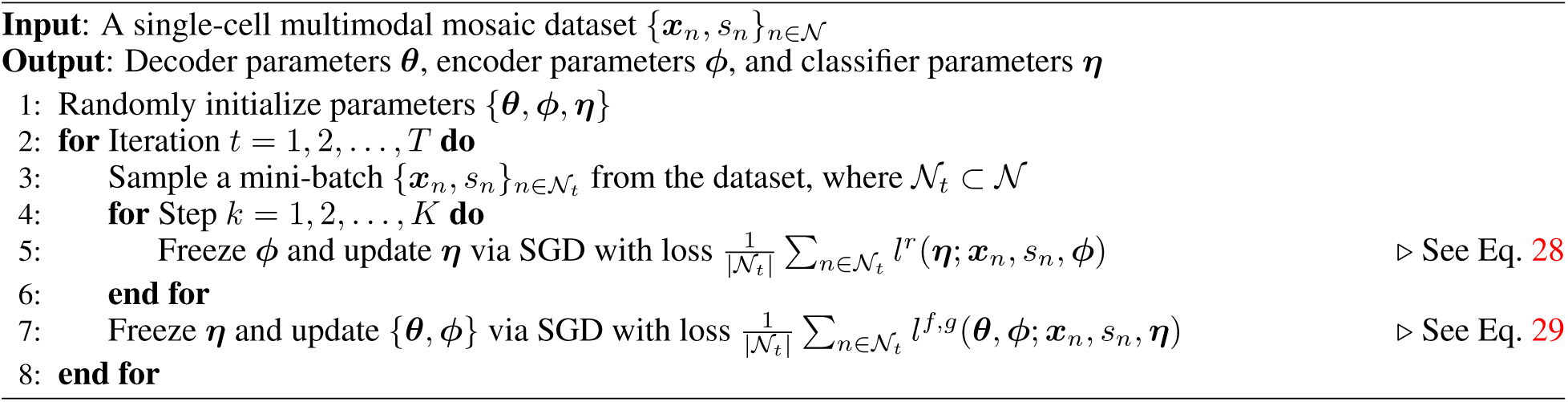

### 4.7 Mosaic integration on latent space

A key goal of single-cell mosaic integration is to extract biological meaningful low-dimensional cell embeddings from the mosaic data for downstream analysis, where the technical variations are removed. To achieve this, for each cell we first use the trained MIDAS to infer the latent posterior *q*_***ϕ***_(***c, u***|***x***_*n*_, *s*_*n*_) through Eq. 10, obtaining the mean 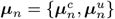 and variance 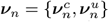. Then, we take the maximum a posteriori (MAP) estimation of {***c, u***} as the integration result on the latent space, which is the mean ***μ***_*n*_ since *q*_***ϕ***_(***c, u***|***x***_*n*_, *s*_*n*_) is Gaussian. Finally, we take 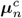, the MAP estimation of ***c***, as the cell embedding.

### 4.8 Imputation for missing modalities and features

Based on the MAP estimation 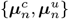 inferred from the single-cell mosaic data (Methods 4.7), it is straightforward to impute missing modalities and features. We first pass 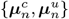 to the decoders to generate padded feature mean 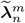 for each modality *m ∈* ℳ via Eq. 18. Then, we sample from a Bernoulli distribution with mean 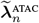 to generate the imputed ATAC counts, and from two Poisson distributions with means 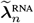 and 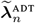 to generate the imputed RNA and ADT counts, respectively.

### 4.9 Batch correction via latent variable manipulation

Besides performing mosaic integration on the latent space (Methods 4.7), we can also perform it on the feature space, *i.e*., imputing missing values and correcting batch effect for the count data. Mosaic integration on feature space is important since it is required by many downstream tasks such as differential expression analysis, cell typing, and trajectory inference.

With the latent variables’ MAP estimation 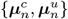, we can perform imputation and batch correction simultaneously by manipulating the technical noise. Concretely, let 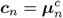 and 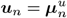, we first calculate the mean of ***u***_*n*_ within each batch *b ∈ ℬ*:

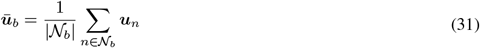

where *𝒩*_*b*_ *⊆ 𝒩* is the set of cell-IDs belonging to batch *b*. Next, we calculate the mean of ***ū***_*b*_ over all batches:

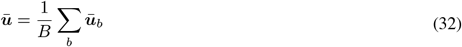

Then, we look for the batch *b*^*^ with the mean ***ū***_*b*_* closest to ***ū***, and treat ***ū***_*b*_* as the “standard” technical noise, where:

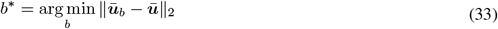

Finally, for each cell we correct the batch effect by substituting ***u***_*n*_ with ***ū***_*b*_*, and pass {***c***_*n*_, ***ū***_*b*_*} to the decoders to generate imputed and batch-corrected data (similar to Methods 4.8 but here we use {***c***_*n*_, ***ū***_*b*_*} instead of {***c***_*n*_, ***u***_*n*_} so as to correct batch effect).

### 4.10 Model transfer via transfer learning

When MIDAS has been pretrained on a reference dataset, we can conduct model transfer to transfer the model’s learned knowledge to a query dataset through transfer learning, *i.e*., on the query dataset we fine-tune the pretrained model instead of train the model from scratch. Since compared to the reference dataset, the query dataset can contain different number of batches collected from different platforms, the batch-ID related modules need to be redefined. Thus, during transfer learning, we reparameterize and reinitialize the batch-ID encoder and decoder {*f*^*s*^, *g*^*s*^} and the batch classifiers {*r*, {*r*^*m*^}_*m* ∈ *ℳ*_}, and only fine-tune the modality encoders and decoders {*f*^*m*^, *g*^*m*^}_*m* ∈ *ℳ*_.

A core advantage of our model transfer scheme is that it can flexibly transfer the knowledge of multimodal data to various types of query datasets, even to those with fewer modalities, improving the *de novo* integration of single-cell data.

### 4.11 Label transfer via reciprocal reference mapping and kNN-based cell annotation

While model transfer implicitly transfers knowledge through model parameters, label transfer explicitly transfers knowledge in the form of data labels. These labels can be different kinds of downstream analysis results such as cell types, cell cycles, or pseudotime. Through accurate label transfer, we can not only avoid the expensive *de novo* integration and downstream analysis, but also improve the label quality.

#### Reciprocal reference mapping

Typically, the first step of label transfer is reference mapping, which aligns the query cells with the reference cells so that labels can be transferred reliably. For MIDAS, we can naively achieve reference mapping in two ways: (1) mapping the query data onto the reference space, *i.e*., applying the model pretrained on the reference data to infer the biological states for the query data [49, 50], and (2) mapping the reference data onto the query space, *i.e*., applying the model transfer-learned on the query data (Methods 4.10) to infer the biological states for the reference data [14, 51]. However, the first way suffers from the “generalization problem” since the pretrained model is hard to generalize to the query data which usually contains unseen technical variations, while the second way suffers from the “forgetting problem” since the transfer-learned model may lose information learned on the reference data, affecting the inferred biological states.

To tackle both problems, we propose a reciprocal reference mapping scheme, where we fine-tune the pretrained model on the query dataset to avoid the generalization problem, and meanwhile feed the model with the historical data sampled from the reference dataset to prevent forgetting. In doing this, the model can find a mapping suitable for both reference and query datasets, and then can align them on the latent space by inferring their biological states.

#### kNN-based cell annotation with novel cell type identification

Based on the aligned latent representations (embeddings), the kNN classifier is employed to transfer the reference labels to the query dataset. When the query and reference datasets belong to the same tissue (*e.g*., PBMCs), we train the kNN classifier using the reference embeddings and labels, and then use it to classify the query cells.

However, if the query and reference datasets are from distinct tissues (*e.g*., BMMCs *vs*. PBMCs), we might encounter new cell types in the query dataset that are not present in the reference dataset. To address this issue, we propose a strategy for novel cell type detection. Specifically, we assign the label “query” to all the query cells, and use the cell embeddings and labels from both the query and reference datasets to train the kNN classifier. Subsequently, we use the classifier to predict the class probabilities for the query cells.

To detect new cell types, we employ a thresholding approach on the predicted probabilities of the “query” class, *i.e*., we leverage a Gaussian Mixture Model (GMM) with two components to group the probabilities into two distinct clusters. This clustering process allows us to establish a suitable threshold for the probabilities. For the cluster with a higher mean, its cells have higher probabilities belonging to the “query” class; we consider these cells as unique to the query dataset and assign them the label “unknown”. Conversely, for the cluster with a lower mean, its cells have lower probabilities belonging to the “query” class; we consider these cells to belong to the types present in the reference dataset, and assign each of these cells the label of the class with the highest predicted probability among all classes except the “query” class.

The above kNN and GMM algorithms are implemented by scikit-learn package’s [57] KNeighborsClassifier function (n_neighbors = 100 and weights = “uniform”) and GaussianMixture function (n_components = 2 and tol = 10^*−*4^), respectively. Similar to model transfer (Methods 4.10), in label transfer knowledge can also be flexibly and accurately transferred to various types of query datasets.

### 4.12 Modality contribution to the integrated clustering

We assess the contribution of different modalities to clustering by measuring the agreement between single-modality clustering and multi-modalities cell clustering. For each cell, the normalized consistency ratio of the nearest neighbors in the single modal clustering and multi-modalities clustering is used to represent contribution of the modal for the final integrated clustering.

### 4.13 Regulatory network inference from scRNA-seq datasets

GRNBoost2 [58] is used to infer the regulatory network from scRNA-seq datasets and is one of the regression-based methods for regulatory network inference. Weighted regulatory links between genes and transcription factors are provided from GRNBoost2. The weights of shared links from different data are compared to indicate the regulatory network retention.

### 4.14 Correlation of expression fold changes between raw and batch-corrected data

For each cell type, expression fold changes of genes and proteins are calculated against all other cells using FoldChange function in Seurat. Pearson correlation coefficient is used to measure linear correlations of fold changes between raw and batch-corrected data.

### 4.15 Generating Seurat cell type labels

To generate cell type labels for both qualitative and quantitative evaluation, we employed the third-party tool, Seurat, to annotate cell types for different datasets through label transfer. We took the CITE-seq PBMC atlas from [15] as the reference set, and utilized the FindTransferAnchors and TransferData functions in Seurat to perform label transfer, where “cca” was used as the reduction method for reference mapping. For cells without raw RNA expression, we first utilized ATAC data to create a gene activity matrix using GeneActivity function in Signac [59]. The gene activity matrix was subsequently used for label transfer.

### 4.16 Evaluation metrics

To evaluate the performance of MIDAS and the state-of-the-art tools on multimodal integration, we utilize metrics from scIB on batch correction and biological conservation, and also propose our own metrics on modality alignment to better evaluate mosaic integration, extending scIB to scMIB (Supplementary Table 4). Since mosaic integration should generate low-dimensional representations and the imputed and batch corrected data, scMIB is performed on both embedding space and feature space. To evaluate the batch correction and biological conservation metrics on the feature space, we convert the imputed and batch corrected feature into a similarity graph via the PCA+WNN strategy (Methods 4.20), and then use this graph for evaluation. Our metrics for batch correction, modality alignment, and biological conservation are defined as follows.

#### 4.16.1 Batch correction metrics

The batch correction metrics comprise graph iLISI (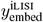 and 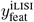), graph connectivity (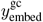 and 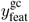), and kBET (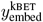 and 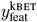), where 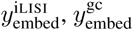, and 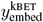 are defined in embedding space and 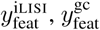, and 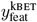 are defined in feature space.

##### Graph iLISI

The graph iLISI (local inverse Simpson’s index [60]) metric is extended from the iLISI, which is used to measure the batch mixing degree. The iLISI scores are computed based on kNN graphs by computing the inverse Simpson’s index for diversity. The scores estimate the effective number of batches present in the neighborhood. iLISI ranges from 1 to *N*, where *N* equals the number of batches. Scores close to the real batch numbers denotes good mixing. However, typical iLISI score is not applicable to graph-based outputs. scIB proposed the Graph iLISI, which utilizes a graph-based distance metric to determine the nearest neighbor list and avoids skews on graph-based integration outputs. The graph iLISI scores are scaled to [0, 1], where 0 indicates strong separation and 1 indicates perfect mixing.

##### Graph connectivity

The graph connectivity is proposed by scIB to inspect whether cells with the same label are connected in the kNN graph of all cells. For each label *c*, we get the largest connected graph component (LCC) of *c*-labeled cells and divide the LCC size by the *c*-labeled cells population size to represent the graph connectivity for cell label *c*. Then, we calculate the connectivity values for all labels and take the average as the total graph connectivity. The score ranges from 0 to 1. The score 1 means that all cells with the same cell identity from different batches are connected in the integrated kNN graph, which also indicates the perfect batch mixing and vice versa.

##### kBET

The kBET (k-nearest neighbor batch-effect test [61]) is used to measure the batch mixing at the local level of the k-nearest neighbors. Certain fractions of random cells are repeatedly selected to test whether the local label distributions are statistically similar to the global label distributions (null hypothesis). The kBET value is the rejection rate over all tested neighborhoods and values close to zero represent that the batches are well mixed. scIB adjusts the kBET with a diffusion-based correction to enable unbiased comparison on graph- and non-graph-based integration results. kBET values are first computed for each label, and then averaged and subtracted from 1 to get a final kBET score.

#### 4.16.2 Modality alignment metrics

The modality alignment metrics comprise modality ASW (*y*^ASW^), FOSCTTM (*y*^FOSCTTM^), label transfer F1 (*y*^ltF1^), ATAC AUROC (*y*^AUROC^), RNA Pearson’s *r* (*y*^RNAr^), and ADT Pearson’s *r* (*y*^ADTr^), where *y*^ASW^, *y*^FOSCTTM^, and *y*^ltF1^ are defined in embedding space and *y*^AUROC^, *y*^RNAr^, and *y*^ADTr^ are defined in feature space.

##### Modality ASW

The modality ASW (averaged silhouette width) is used to measure the alignment of distributions between different modality embeddings. The ASW [62] is originally used to measure the separation of clusters. In scIB, ASW is also modified to measure the performance of batch effect removal, resulting in a batch ASW that ranges from 0 to 1, where 1 denotes perfect batch mixing and 0 denotes strong batch separation. By replacing batch embeddings with modality embeddings, we can define a modality ASW in the same manner as the batch ASW, where 1 denotes perfect modality alignment and 0 denotes strong modality separation. For MIDAS, the modality embeddings are generated by feeding the trained model with each modality individually.

##### FOSCTTM

The FOSCTTM (fraction of samples closer than the true match [63]) is used to measure the alignment of values between different modality embeddings. Let 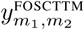 be the FOSCTTM for a modality pair {*m*_1_, *m*_2_}, it is defined as:

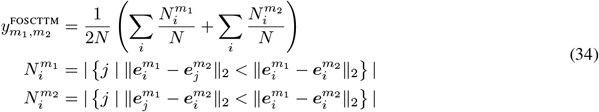

where *N* is the number of cells, *i* and *j* are the cell indices, and 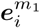 and 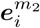 are the embeddings of cell *i* in modalities *m*_1_ and *m*_2_, respectively. 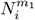 is the number of cells in modality *m*_2_ that are closer to 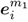 than 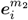 is to 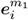, and it is similar for 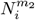. We first get the embeddings of individual modalities, then calculate the FOSCTTM values for each modality pair, and lastly average these values and subtract it from 1 to obtain a final FOSCTTM score. Higher FOSCTTM scores indicate better modality alignment.

##### Label transfer F1

The label transfer F1 is used to measure the alignment of cell types between different modality embeddings. This can be achieve by testing whether cell type labels can be transferred from one modality to another without any bias. For each pair of modalities, we first build a kNN graph between their embeddings, and then transfer labels from one modality to the other based on the nearest neighbors. The transferred labels are compared to the original labels by the micro F1-score, which is defined as the label transfer F1. We take the F1 score averaged from all comparison pairs as the final label transfer F1 score.

##### ATAC AUROC

The ATAC AUROC (area under the receiver operating characteristic) is used to measure the alignment of different modalities in the ATAC feature space. It has been previously used to evaluate the quality of ATAC predictions [64]. For each method to be evaluated, we first use it to convert different modality combinations that do not contain ATAC into ATAC features respectively, then calculate the AUROC of each converted result by taking the true ATAC features as the ground-truth, and finally take the average of these AUROCs as the final score. Taking MIDAS as an example, if ATAC, RNA and ADT are involved, the evaluation is based on the combinations {RNA}, {ADT}, and {RNA, ADT}. For each combination, we feed the data into the trained model to generate the imputed data of all modalities {ATAC, RNA, ADT} (Methods 4.8), where the generated ATAC features are used for AUROC calculation.

##### RNA Pearson’s *r*

The RNA Pearson’s *r* is used to measure the alignment of different modalities in the RNA feature space. For each method to be evaluated, we first use it to convert different modality combinations that do not contain RNA into RNA features respectively, then calculate the Pearson’s *r* between each converted result and the true RNA features, and finally take the average of these Pearson’s *r*s as the final score.

##### ADT Pearson’s *r*

The ADT Pearson’s *r* is used to measure the alignment of different modalities in the ADT feature space. The calculation of the ADT Pearson’s *r* is similar to that of the RNA Pearson’s *r*.

#### 4.16.3 Biological conservation metrics

The biological conservation metrics comprise NMI (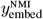 and 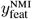), ARI (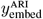 and 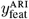), isolated label F1 (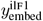 and 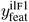), and graph cLISI (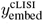 and 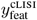), where 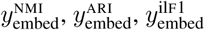, and 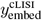 are defined in embedding space and 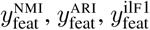, and 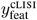 are defined in feature space.

##### NMI

The NMI (Normalized Mutual Information) is used to measure the similarity between two clustering results, namely the predefined cell type labels and the clustering result obtained from the embeddings or the graph. Optimized Louvain clustering is used here according to scIB. The NMI scores are scaled to [0, 1] where 0 and 1 correspond to uncorrelated clustering and a perfect match, respectively.

##### ARI

The ARI (Adjusted Rand Index) also measures the overlap of two clustering results. The RI (Rand Index [65]) considers not only cell pairs that are assigned in the same clusters but also ones in different clusters in the predicted (Louvain clustering) and true (cell type) clusters. ARI corrects RI for randomly correct labels. An ARI of 1 represents perfect match and 0 represents random labeling.

##### Isolated label F1

scIB proposes the isolated label F1 score to evaluate the integration performance, specifically focusing on cells with the label that is share by few batches. Cell labels presented in the least number of batches are identified as isolated labels. The F1 score for measuring the clustering performance on isolated labels is defined as the isolated label F1 score. It reflects how well the isolated labels separate from other cell identities, ranging from 0 to 1, where 1 means all the isolated label cells and no others are grouped into one cluster.

##### Graph cLISI

The Graph cLISI is similar to the Graph iLISI but focuses on cell type labels rather than batch labels. Unlike iLISI that highlights the mixing of groups, cLISI values the separation of groups. The graph-adjusted cLISI is scaled to [0, 1] with value 0 corresponding to low cell-type separation and 1 corresponding to strong cell-type separation.

#### 4.16.4 Overall scores

##### scIB

We compute the scIB overall score using the batch correction and biological conservation metrics defined either on the embedding space (for algorithms generating embeddings or graphs) or the feature space (for algorithms generating batch-corrected features). Following [43], the overall score *y* is the sum of the averaged batch correction metric *y*^batch^ weighted by 0.4 and the averaged biological conservation metric *y*^bio^ weighted by 0.6:

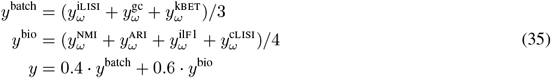

where *ω* = embed for embedding or graph outputs, and *ω* = feat for feature outputs.

##### scMIB

As an extension of scIB, the scMIB overall score *y* is computed from the batch correction, modality alignment, and biological conservation metrics defined on both the embedding and feature space. It is the sum of the averaged batch correction metric *y*^batch^ weighted by 0.3, the averaged modality alignment metric *y*^mod^ weighted by 0.3, and the averaged biological conservation metric *y*^bio^ weighted by 0.4:

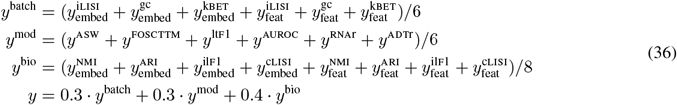

### 4.17 Datasets

All datasets of human PBMCs were publicly available (Supplementary Table 1). Count matrices of gene UMIs, ATAC fragments and antibody-derived tags were downloaded for data analysis.

#### DOGMA dataset

The DOGMA dataset contains four batches profiled by DOGMA-seq, which measures RNA, ATAC and ADT data simultaneously. Trimodal data of this dataset were obtained from Gene Expression Omnibus (GEO) [66] under accession ID GSE166188 [3].

#### TEA dataset

The TEA dataset contains five batches profiled by TEA-seq, which measures RNA, ATAC and ADT data simultaneously. Trimodal data of these batches were obtained from GEO under accession ID GSE158013 [4].

#### TEA Multiome dataset

The TEA Multiome dataset measuring paired RNA and ATAC data was obtained from GEO under accession ID GSE158013 [4]. This dataset contains two batches profiled by 10x Chromium Single Cell Multiome ATAC + Gene Expression.

#### 10X Multiome dataset

The 10X Multiome dataset measuring paired RNA and ATAC data was collected from 10x Genomics (https://www.10xgenomics.com/resources/datasets/) [67–70].

#### ASAP dataset

The ASAP dataset was obtained from GEO under accession ID GSE156473 [3]. Two batches profiled by ASAP-seq are used, which include ATAC and ADT data.

#### ASAP CITE dataset

The ASAP CITE dataset was obtained from GEO under accession ID GSE156473 [3]. Two batches profiled by CITE-seq are used, which include RNA and ADT data.

#### WNN CITE dataset

The WNN CITE dataset measuring paired RNA and ADT data was obtained from https://atlas.fredhutch.org/nygc/multimodal-pbmc [15]. This dataset was profiled by CITE-seq. We selected the eight PBMC batches generated before the administration of HIV vaccine for integration.

#### BMMC mosaic dataset

The BMMC mosaic dataset included three batches. The ICA batch measuring RNA data was obtained from https://www.dropbox.com/s/xe5tithw1xjxrfs/ica_bone_marrow.h5?dl=0 [71]. The ASAP batch measuring ADT and ATAC data was obtained from GEO under accession ID GSE156477 [3]. The CITE batch measuring RNA and ADT data was obtained from GEO under accession ID GSE128639 [13].

### 4.18 Data preprocessing

The count matrices of RNA and ADT were processed via the Seurat package (v4.1.0) [15]. The ATAC fragment files were processed using the Signac package (v1.6.0) [59] and peaks were called via MACS2 [72]. We performed quality control separately for each batch. Briefly, metrics of detected gene number per cell, total UMI number, percentage of mtRNA reads, total protein tag number, total fragment number, TSS score, and nucleosome signal were evaluated. We manually checked the distributions of these metrics and set customized criteria to filter low-quality cells in each batch. The number of cells that passed quality control in each batch is shown in Supplementary Table 1.

For each batch, we adopted common normalization strategies for RNA, ADT and ATAC respectively. Specifically, for RNA data, UMI count matrices are normalized and log-transformed using the NormalizeData function in Seurat; for ADT data, tag count matrices are centered log-ratio (CLR)-normalized using the NormalizeData function in Seurat; for ATAC data, fragment matrices are term frequency inverse document frequency (TF-IDF) normalized using the RunTFIDF function in Signac.

To integrate batches profiled by various technologies, we need to create a union of features for RNA, ADT and ATAC data, respectively. For RNA data, firstly, low-frequency genes are removed based on gene occurrence frequency across all batches; then we selected 4000 high variable genes (HVGs) using the FindVariableFeatures function with default parameters in each batch; the union of these HVGs are ranked using the SelectIntegrationFeatures function and the top 4000 genes are selected. In addition, we also retained genes that encode the proteins targeted by the antibodies. For ADT data, the union of antibodies in all batches are retained for data integration. For ATAC data, we used reduce function in Signac to merge all intersecting peaks across batches and then re-calculated the fragment counts in the merged peaks. The merged peaks are used for data integration.

The input data for MIDAS are UMI counts for RNA data, tag counts for ADT data, and binarized fragment counts for ATAC data. For each modality, the union of features from all batches are used. Counts of missing features are set to 0. Binary feature masks are generated accordingly where 1 and 0 denote presented and missing features, respectively.

### 4.19 Implementation of MIDAS

We implement the MIDAS architecture using PyTorch [73]. The sizes of the shared hidden layers for different modality encoders are set to 1024-128, while the sizes of the shared hidden layers for different modality decoders are set to 128-1024. Additionally, the sizes of the biological state and technical noise latent variables are set to 32 and 2, respectively (refer to Supplementary Fig. 1 and Supplementary Table 12 for details). Each hidden layer is constructed using four PyTorch functions: Linear, LayerNorm, Mish, and Dropout. The input and output layers have varying sizes depending on the datasets used (refer to Supplementary Table 13 for details). In order to effectively reduce the number of model parameters, similar to [64], the input and reconstruction layers for the ATAC modality are both split into 22 independent fully-connected layers based on the genomic regions of different human chromosomes (excluding the sex chromosomes).

To train MIDAS, we set the modality alignment loss weight (*α*) to 50, the technical IB loss weight (*β*^*s*^) to 30, the biological IB loss weight (*β*^*x*^) to 4, and the technical noise likelihood loss weight (*γ*) to 1000. The number of updates (*K*) of the batch classifiers {*r*, {*r*^*m*^}_*m* ∈ *ℳ*_} in each iteration is set to 3. We split the dataset into training and validation sets in a ratio of 95:5. The mini-batch size is set to 256, and we utilize the AdamW [74] optimizer with a learning rate of 10^*−*4^ for implementing SGD. We train the model for a maximum of 2000 epochs and use early stopping to terminate training. The dropout rates for all hidden layers are set to 0.2. All the hyperparameter settings for MIDAS training are listed in Supplementary Table 14.

### 4.20 Implementation of comparing methods

We compared MIDAS with 19 recent methods on different trimodal and bimodal integration tasks (see Supplementary Table 3 for an overview). If a method cannot handle missing features for a certain modality, we used the feature intersection of different batches of that modality for integration. For a fair comparison, we set the size of the low-dimensional representations generated by each method to be 32, the same as that of the biological states inferred by MIDAS. For other settings of each method, if not specified, their default values were used. For trimodal rectangular integration tasks, since few methods are directly applicable to ATAC, RNA, and ADT trimodal data, we decomposed the rectangular integration into two steps, *i.e*., batch correction for each modality independently, and modality fusion for all batch-corrected modalities. We then combined different batch correction and modality fusion methods to achieve rectangular integration, resulting in 9 different strategies in total.

#### 4.20.1 Methods compared in trimodal rectangular integration tasks

##### BBKNN+average

BBKNN [75] is used for batch correction (embedding space) and graph averaging is used for modality fusion. For each batch, we employ functions from the Seurat package to perform dimensionality reduction on the count data. We first use RunTFIDF and RunSVD functions to obtain the low-dimensional representation of ATAC, and then use NormalizeData, ScaleData, and RunPCA functions to obtain the low-dimensional representations of RNA and ADT, respectively. For the obtained low-dimensional representation of each modality, we use the bbknn function of the Scanpy package to remove the batch effect and obtain an similarity graph. Finally, we average the similarity graphs of different modalities to obtain the output.

##### Harmony+WNN

Harmony [60] is used for batch correction (embedding space) and WNN [15] is used for modality fusion. We use the same processing method as BBKNN+average to obtain low-dimensional representations of different batches of ATAC, RNA, and ADT respectively. For the obtained low-dimensional representation of each modality, we employ the RunHarmony function of the Harmony package to remove batch effects. Then, we use Seurat’s FindMultiModalNeighbors function, *i.e*., the WNN algorithm, to fuse the low-dimensional representations of different modalities to obtain the graph output.

##### LIGER+WNN

LIGER [76] is used for batch correction (embedding space) and WNN is used for modality fusion. For each batch, we employ Seurat’s RunTFIDF and ScaleData functions for ATAC normalization and its NormalizeData and ScaleData functions for RNA and ADT normalization. For each modality, we then use RunOptimizeALS and RunQuantileNorm functions of the LIGER package for dimensionality reduction and batch effect removal. Finally, we use the WNN algorithm FindMultiModalNeighbors function to fuse the low-dimensional representations of different modalities to obtain the graph output.

##### MOFA+

MOFA+ [24] is used for simultaneous batch correction (embedding space) and modality fusion. We first use the same processing method as LIGER+WNN to normalize each modality separately, and then use run_mofa and get_factors functions of the MOFA+ package to achieve simultaneous batch effect removal and modality fusion on the normalized data, obtaining low-dimensional representations output.

##### PCA+WNN

SVD (for ATAC) and PCA (for RNA and ADT) are used for dimensionality reduction (no batch correction is applied) and WNN is used for modality fusion. We use the same processing method as BBKNN+average to obtain low-dimensional representations of different batches of ATAC, RNA, and ADT respectively. Then, we use the WNN algorithm FindMultiModalNeighbors function to fuse the low-dimensional representations of different modalities to obtain the graph output.

##### Scanorama-embed+WNN

Scanorama [76] is used for batch correction (embedding space) and WNN is used for modality fusion. For each modality, we employ integrate function from the Scanorama package for dimensionality reduction and batch effect removal. Then, we use the WNN algorithm FindMultiModalNeighbors function to fuse the low-dimensional representations of different modalities to obtain the graph output.

##### Scanorama-feat+WNN

Scanorama is used for batch correction (feature space) and WNN is used for modality fusion. For each modality, we perform batch correction using correct function of the Scanorama package. For the batch-corrected count data, we then use the PCA+WNN strategy to get the graph output.

##### Seurat-CCA+WNN

Seurat’s CCA [13] is used for batch correction (feature space) and WNN is used for modality fusion. For each modality, we use Seurat’s FindIntegrationAnchors function (reduction = “cca”), *i.e*., the CCA algorithm, to anchor different batches, and use its IntegrateData function to correct batch effects. For the batch-corrected count data, we then use the PCA+WNN strategy to get the graph output.

##### Seurat-RPCA+WNN

Seurat’s RPCA [13] is used for batch correction (feature space) and WNN is used for modality fusion. It uses the same strategy as Seurat-CCA+WNN, except that the FindIntegrationAnchors function is applied with reduction = “rpca”.

#### 4.20.2 Methods compared in trimodal mosaic integration tasks

##### Multigrate

Multigrate [29] can be found on GitHub at https://github.com/theislab/multigrate. For data inputs, we took the intersection of genes in scRNA-seq data and proteins in ADT data across different batches. We processed the data using the default method of Multigrate. The values of parameters KL and integ were set to 0.1 and 3000, respectively.

##### scMoMaT

scMoMaT [27] is designed to integrate multimodal mosaic data. The code is downloaded from https://github.com/PeterZZQ/scMoMaT. We take the same preprocessed data as MIDAS. For each modality, since scMoMaT does not handle missing features, we only use the intersected features of different batches of the preprocessed data for integration. We set the mini-batch size to 0.1*×*N for training, where N is the number of cells.

##### scVAEIT

scVAEIT [26] is designed to integrate multimodal mosaic data. The code is downloaded from https://github.com/jaydu1/scVAEIT. After filtering the low-quality cells and features as MIDAS did, we size-normalized and log-normalized the counts of gene and protein separately, while binarized the peaks by changing all nonzero values to 1.

##### StabMap

StabMap [28] is designed to integrate single-cell data with non-overlapping features. The code is downloaded from https://github.com/MarioniLab/StabMap. In order to select suitable highly variable features, we set different parameters for different modalities (mean *>* 0.01 and p.value *≤* 0.05 for RNA; mean *>* 0.25 and p.value *≤* 0.05 for ATAC; mean *>* 0.01 and p.value *≤* 0.1 for ADT). In the case of diagonal integration, since there are no shared features between different modalities, we convert the ATAC regions into the nearest genes using the ClosestFeature function in Signac, and convert the ADT proteins into the corresponding genes. In addition, in order to obtain more shared features between different modalities in diagonal integration, we relaxed the conditions of highly variable features (mean *>* 0.01 and p.value *≤* 0.1 for RNA; mean *>* 0.25 and p.value *≤* 0.05 for ATAC; all features for ADT). In diagonal integration, we choose the RNA batch as the reference set; in other cases, we choose the batch with the largest number of modalities as the reference set.

#### 4.20.3 Methods compared in bimodal (ATAC+RNA) integration tasks

##### Cobolt

Cobolt [21] is designed to integrate bimodal mosaic data from ATAC and RNA. The code is downloaded from https://github.com/epurdom/cobolt. We take the same preprocessed data as MIDAS and retain the intersected features of each modality for different batches. In order for Cobolt to read, we store the preprocessed data of each modality in each batch as a SingleData object. We set the learning rate to 5 *×* 10^*−*4^.

##### MultiVI

MultiVI [22] is designed to integrate bimodal mosaic data from ATAC and RNA. The code is downloaded from https://github.com/scverse/scvi-tools. We take the same preprocessed data as MIDAS. For each modality, we also retain intersected features of different batches. In the model setup, we use batch_key to specify the cell modality, and use categorical_covariate_keys to specify the cell batch.

##### uniPort

uniPort [23] is designed to integrate heterogeneous single-cell bimodal data. The source code can be downloaded from https://github.com/caokai1073/uniPort. Since uniPort supports horizontal, vertical, and diagonal integration, we combine all three integration methods to achieve our tasks.

##### GLUE

GLUE [25] is a tool designed for integrating unpaired multi-modal data (e.g. scRNA-seq, scATAC-seq, snmC-seq, etc.) using graph link unified embeddings. The code can be downloaded from https://github.com/gao-lab/GLUE, while the GTF file we used in the experiments can be obtained from https://ftp.ebi.ac.uk/pub/databases/gencode/Gencode_human/release_43/gencode.v43.annotation.gtf.gz. To remove batch effects, we set use_batch = “batch” in the experiments.

#### 4.20.4 Methods compared in bimodal (RNA+ADT) integration tasks

##### totalVI

totalVI [18] is designed to integrate bimodal mosaic data from RNA and ADT. It can be found on GitHub at https://github.com/scverse/scvi-tools/tree/main. As totalVI does not handle missing genes, we took the intersection of genes in RNA data from different input batches. For the ADT data, the union of proteins from different batches are used.

##### sciPENN

sciPENN [19] features and the union of ADT features https://github.com/jlakkis/sciPENN. Due to sciPENN’s incapability in dealing with missing genes, we took the intersection of RNA features and the union of ADT features for different input batches.

#### Reporting Summary

Further information on research design is available in the Nature Research Reporting Summary linked to this article.

## Data availability

All single-cell datasets of human PBMCs and BMMCs used in this paper are publicly available. See Supplementary Table 1 for detailed information.

