## Supplementary Notes, Figures, and Tables for "MIDAS: a deep generative model for mosaic integration and knowledge transfer of single-cell multimodal data"

### Supplementary Note 1: Evaluation of the robustness, versatility, and efficiency of MIDAS in mosaic integration tasks

Given the several hyperparameters in MIDAS architecture, we systematically evaluated the performance of MIDAS with different settings of each hyperparameter on eight mosaic integration tasks. The results show that the default values of 32 for biological state latent variable dimension ($D^c$) and 3 for hidden layer number ($L$) respectively achieve the highest overall scores and ensure compatibility with complex data while maintaining comparable performance (Supplementary Fig. 13). For the other five hyperparameters, MIDAS exhibited considerable robustness to hyperparameter settings across most tasks (Supplementary Fig. 13). However, disabling these five modules always results in a decrease of MIDAS's performance.

In real-world scenario, MIDAS may suffer from biological or technical differences across batches. Thus we conducted evaluations of MIDAS on integrating constructed mosaic datasets with missing cell types or different sequencing depths (Supplementary Table 5, 6). MIDAS is slightly affected by missing cell types. Loss of major cell types has a greater impact on MIDAS's performance than the loss of minor cell types (Supplementary Fig. 14a). The UMAP plots indicate that MIDAS can effectively prevent batch overcorrection while aligning cell types from different datasets (Supplementary Fig. 15). When given gradually decreasing different depths, the model performance decreased slightly in most datasets, except for the diagonal+full and diagonal datasets (Supplementary Fig. 14b). However, even when the sequencing depth sampling rate was 0.05, the scMIB score of MIDAS were still maintained above 0.7, except for the diagonal+full and diagonal datasets.

Apart from the 13 trimodal integration methods, we also compared MIDAS against other six methods on bimodal mosaic data (Supplementary Table 3). In all ten bimodal integration tasks (Supplementary Table 7, 8), MIDAS outperformed the other methods on both dogma and teadog datasets (Supplementary Fig. 16, 17).

We evaluated the computing time and memory consumed by MIDAS and 19 other approaches on a Linux server with 36 CPU cores (two Intel Xeon Gold 6240 chips), 512 GB of RAM, and NVIDIA Tesla V100S GPUs. MIDAS is the fastest deep learning method with the least memory consumption for tri-omics integration tasks (Supplementary Fig. 18). As for the bi-omics integration tasks, MIDAS is comparable to other methods.

### Supplementary Note 2: Derivation of the variational posterior $q_{\boldsymbol{\phi}}(\boldsymbol{z}|\boldsymbol{x}_n, s_n)$

The detailed derivation of the variational posterior $q_{\boldsymbol{\phi}}(\boldsymbol{z}|\boldsymbol{x}_n, s_n)$ for Eq. 9 is as follow:

$$\begin{aligned}
p(\boldsymbol{z}|\boldsymbol{x}_n, s_n) &= \frac{p(\boldsymbol{z})p(s_n|\boldsymbol{z})p(\boldsymbol{x}_n|\boldsymbol{z})}{p(\boldsymbol{x}_n, s_n)} \\
&= \frac{p(\boldsymbol{z})}{p(\boldsymbol{x}_n, s_n)} p(s_n|\boldsymbol{z}) \prod_{m \in \mathcal{M}_n} p(\boldsymbol{x}_n^m|\boldsymbol{z}) \\
&= \frac{p(\boldsymbol{z})}{p(\boldsymbol{x}_n, s_n)} \frac{p(s_n)p(\boldsymbol{z}|s_n)}{p(\boldsymbol{z})} \prod_{m \in \mathcal{M}_n} \frac{p(\boldsymbol{x}_n^m)p(\boldsymbol{z}|\boldsymbol{x}_n^m)}{p(\boldsymbol{z})} \\
&= \frac{p(s_n)}{p(\boldsymbol{x}_n, s_n)} \left( \prod_{m \in \mathcal{M}_n} p(\boldsymbol{x}_n^m) \right) p(\boldsymbol{z}) \frac{p(\boldsymbol{z}|s_n)}{p(\boldsymbol{z})} \prod_{m \in \mathcal{M}_n} \frac{p(\boldsymbol{z}|\boldsymbol{x}_n^m)}{p(\boldsymbol{z})} \\
&\approx \frac{p(s_n)}{p(\boldsymbol{x}_n, s_n)} \left( \prod_{m \in \mathcal{M}_n} p(\boldsymbol{x}_n^m) \right) p(\boldsymbol{z}) \frac{q_{\boldsymbol{\phi}}(\boldsymbol{z}|s_n)}{p(\boldsymbol{z})} \prod_{m \in \mathcal{M}_n} \frac{q_{\boldsymbol{\phi}}(\boldsymbol{z}|\boldsymbol{x}_n^m)}{p(\boldsymbol{z})} \\
&\triangleq q_{\boldsymbol{\phi}}(\boldsymbol{z}|\boldsymbol{x}_n, s_n)
\end{aligned} \tag{37}$$

where $q_{\boldsymbol{\phi}}(\boldsymbol{z}|s_n)$ and $q_{\boldsymbol{\phi}}(\boldsymbol{z}|\boldsymbol{x}_n^m)$ are the variational approximations of the true posteriors $p(\boldsymbol{z}|s_n)$ and $p(\boldsymbol{z}|\boldsymbol{x}_n^m)$, respectively.

#### Supplementary Note 3: Derivation of the Information Bottleneck training objective

Here we derive the IB objective of Eq. 23. Starting from the definition of $\beta^s \, \mathrm{I}(s, \boldsymbol{c}) + \beta^x \, \mathrm{I}(\boldsymbol{x}, \boldsymbol{u})$ we have:

$$
\begin{aligned}
\beta^s \, \mathrm{I}(s, \boldsymbol{c}) + \beta^x \, \mathrm{I}(\boldsymbol{x}, \boldsymbol{u}) &= \beta^s \mathbb{E}_{p(s,\boldsymbol{c})}\left[\log \frac{p(s,\boldsymbol{c})}{p(s)p(\boldsymbol{c})}\right] + \beta^x \mathbb{E}_{p(\boldsymbol{x},\boldsymbol{u})}\left[\log \frac{p(\boldsymbol{x},\boldsymbol{u})}{p(\boldsymbol{x})p(\boldsymbol{u})}\right] \\
&= \beta^s \mathbb{E}_{p(s,\boldsymbol{c})}\big[\log \underbrace{p(s|\boldsymbol{c})}_{\approx p_{\widehat{\boldsymbol{\eta}}}(s|\boldsymbol{c})}\big] \underbrace{- \beta^s \mathbb{E}_{p(s,\boldsymbol{c})}\left[\log p(s)\right]}_{\text{const.}} + \beta^x \mathbb{E}_{p(\boldsymbol{x},\boldsymbol{u})}\left[\log \frac{p(\boldsymbol{u}|\boldsymbol{x})}{p(\boldsymbol{u})}\right] \\
&\approx \beta^s \mathbb{E}_{p(\boldsymbol{x},s)}\left[\mathbb{E}_{p(\boldsymbol{c}|\boldsymbol{x},s)}\left[\log p_{\widehat{\boldsymbol{\eta}}}(s|\boldsymbol{c})\right]\right] + \text{const.} + \beta^x \mathbb{E}_{p(\boldsymbol{x},s)}\left[\mathbb{E}_{p(\boldsymbol{u}|\boldsymbol{x},s)}\left[\log \frac{p(\boldsymbol{u}|\boldsymbol{x})}{p(\boldsymbol{u})}\right]\right] \\
&= \mathbb{E}_{p(\boldsymbol{x},s)}\left[\beta^s \mathbb{E}_{p(\boldsymbol{c}|\boldsymbol{x},s)}\left[\log p_{\widehat{\boldsymbol{\eta}}}(s|\boldsymbol{c})\right] + \beta^x \mathbb{E}_{p(\boldsymbol{u}|\boldsymbol{x},s)}\left[\log \frac{p(\boldsymbol{u}|\boldsymbol{x})}{p(\boldsymbol{u})}\right]\right] + \text{const.} \\
&\approx \frac{1}{N}\sum_n \left[\beta^s \mathbb{E}_{p(\boldsymbol{c}|\boldsymbol{x}_n,s_n)}\left[\log p_{\widehat{\boldsymbol{\eta}}}(s_n|\boldsymbol{c})\right] + \beta^x \mathbb{E}_{p(\boldsymbol{u}|\boldsymbol{x}_n,s_n)}\left[\log \frac{p(\boldsymbol{u}|\boldsymbol{x}_n)}{p(\boldsymbol{u})}\right]\right] + \text{const.} \\
&\approx \frac{1}{N}\sum_n \left[\beta^s \mathbb{E}_{q_{\boldsymbol{\phi}}(\boldsymbol{c}|\boldsymbol{x}_n,s_n)}\left[\log p_{\widehat{\boldsymbol{\eta}}}(s_n|\boldsymbol{c})\right] + \beta^x \mathbb{E}_{q_{\boldsymbol{\phi}}(\boldsymbol{u}|\boldsymbol{x}_n,s_n)}\left[\log \frac{q_{\boldsymbol{\phi}}(\boldsymbol{u}|\boldsymbol{x}_n)}{p(\boldsymbol{u})}\right]\right] + \text{const.}
\end{aligned} \tag{38}
$$

To calculate $\mathbb{E}_{q_{\boldsymbol{\phi}}(\boldsymbol{u}|\boldsymbol{x}_n,s_n)}\left[\log \frac{q_{\boldsymbol{\phi}}(\boldsymbol{u}|\boldsymbol{x}_n)}{p(\boldsymbol{u})}\right]$, we first define its variational posterior term $q_{\boldsymbol{\phi}}(\boldsymbol{u}|\boldsymbol{x}_n)$ based on the true posterior $p(\boldsymbol{u}|\boldsymbol{x}_n)$:

$$
\begin{aligned}
p(\boldsymbol{u}|\boldsymbol{x}_n) &= p(\boldsymbol{u}|\boldsymbol{x}_n)\underbrace{\frac{p(s_n|\boldsymbol{u},\boldsymbol{x}_n)}{p(s_n|\boldsymbol{u},\boldsymbol{x}_n)}}_{=p(s_n|\boldsymbol{u})}\frac{p(\boldsymbol{x}_n)}{p(\boldsymbol{x}_n)} \\
&= \frac{p(\boldsymbol{u},\boldsymbol{x}_n,s_n)}{p(s_n|\boldsymbol{u})p(\boldsymbol{x}_n)} \\
&= \frac{p(\boldsymbol{u}|\boldsymbol{x}_n,s_n)p(\boldsymbol{x}_n,s_n)}{p(s_n|\boldsymbol{u})p(\boldsymbol{x}_n)} \\
&\approx \frac{q_{\boldsymbol{\phi}}(\boldsymbol{u}|\boldsymbol{x}_n,s_n)p(\boldsymbol{x}_n,s_n)}{p_{\boldsymbol{\theta}}(s_n|\boldsymbol{u})p(\boldsymbol{x}_n)} \\
&\triangleq q_{\boldsymbol{\phi}}(\boldsymbol{u}|\boldsymbol{x}_n)
\end{aligned} \tag{39}
$$

where $p(s_n|\boldsymbol{u},\boldsymbol{x}_n) = p(s_n|\boldsymbol{u})$ since $s$ and $\boldsymbol{x}$ are conditionally independent given $\boldsymbol{u}$. Substituting $q_{\boldsymbol{\phi}}(\boldsymbol{u}|\boldsymbol{x}_n)$ into the expectation $\mathbb{E}_{q_{\boldsymbol{\phi}}(\boldsymbol{u}|\boldsymbol{x}_n,s_n)}\left[\log \frac{q_{\boldsymbol{\phi}}(\boldsymbol{u}|\boldsymbol{x}_n)}{p(\boldsymbol{u})}\right]$ then yields:

$$
\begin{aligned}
\mathbb{E}_{q_{\boldsymbol{\phi}}(\boldsymbol{u}|\boldsymbol{x}_n,s_n)}\left[\log \frac{q_{\boldsymbol{\phi}}(\boldsymbol{u}|\boldsymbol{x}_n)}{p(\boldsymbol{u})}\right] &= \mathbb{E}_{q_{\boldsymbol{\phi}}(\boldsymbol{u}|\boldsymbol{x}_n,s_n)}\left[\log \frac{q(\boldsymbol{u}|\boldsymbol{x}_n,s_n)p(\boldsymbol{x}_n,s_n)}{p(\boldsymbol{u})p_{\boldsymbol{\theta}}(s_n|\boldsymbol{u})p(\boldsymbol{x}_n)}\right] \\
&= \mathbb{E}_{q_{\boldsymbol{\phi}}(\boldsymbol{u}|\boldsymbol{x}_n,s_n)}\left[\log \frac{q(\boldsymbol{u}|\boldsymbol{x}_n,s_n)}{p(\boldsymbol{u})} - \log p_{\boldsymbol{\theta}}(s_n|\boldsymbol{u}) + \log \frac{p(\boldsymbol{x}_n,s_n)}{p(\boldsymbol{x}_n)}\right] \\
&= \mathrm{KL}\left(q_{\boldsymbol{\phi}}(\boldsymbol{u}|\boldsymbol{x}_n,s_n)\|p(\boldsymbol{u})\right) - \mathbb{E}_{q_{\boldsymbol{\phi}}(\boldsymbol{u}|\boldsymbol{x}_n,s_n)}\left[\log p_{\boldsymbol{\theta}}(s_n|\boldsymbol{u})\right] + \text{const.}
\end{aligned} \tag{40}
$$

By further substituting Eq. 40 into Eq. 38, we arrive at Eq. 23:

$$
\begin{aligned}
\beta^s \, \mathrm{I}(s, \boldsymbol{c}) + \beta^x \, \mathrm{I}(\boldsymbol{x}, \boldsymbol{u}) \approx & \frac{1}{N}\sum_n \Big[\beta^s \mathbb{E}_{q_{\boldsymbol{\phi}}(\boldsymbol{c}|\boldsymbol{x}_n,s_n)}\left[\log p_{\widehat{\boldsymbol{\eta}}}(s_n|\boldsymbol{c})\right] + \beta^x \, \mathrm{KL}\left(q_{\boldsymbol{\phi}}(\boldsymbol{u}|\boldsymbol{x}_n,s_n)\|p(\boldsymbol{u})\right) \\
& - \beta^x \mathbb{E}_{q_{\boldsymbol{\phi}}(\boldsymbol{u}|\boldsymbol{x}_n,s_n)}\left[\log p_{\boldsymbol{\theta}}(s_n|\boldsymbol{u})\right]\Big] + \text{const.}
\end{aligned}
$$

where within the summation term on the right-hand side, the first term is approximated by feeding the sample $\boldsymbol{c}_n \sim q_{\boldsymbol{\phi}}(\boldsymbol{c}|\boldsymbol{x}_n, s_n)$ into the classifier $r$, the second term (the Gaussian KLD) is calculated analytically, and the third term is approximated by feeding the sample $\boldsymbol{u}_n \sim q_{\boldsymbol{\phi}}(\boldsymbol{u}|\boldsymbol{x}_n, s_n)$ into the batch-ID decoder $g^s$.

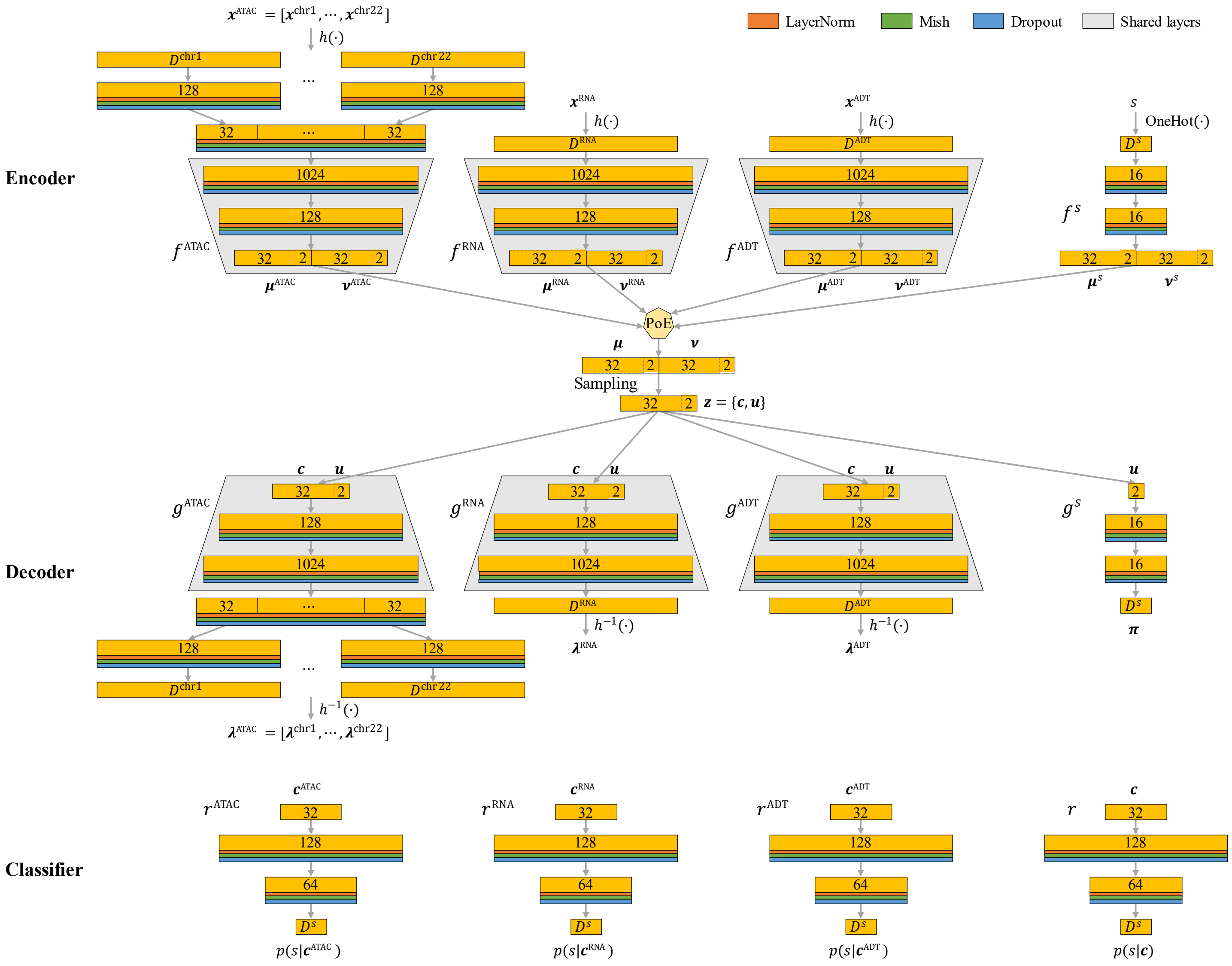

**Supplementary Figure 1. Neural network architecture for MIDAS.** The sizes (number of dimensions) are marked on each layer (yellow), where the sizes of the input and output layers depend on the task. In order to effectively reduce the number of model parameters, the input and reconstruction layers for the ATAC modality are both split into 22 independent fully-connected layers based on the genomic regions of different human chromosomes (excluding the sex chromosomes).

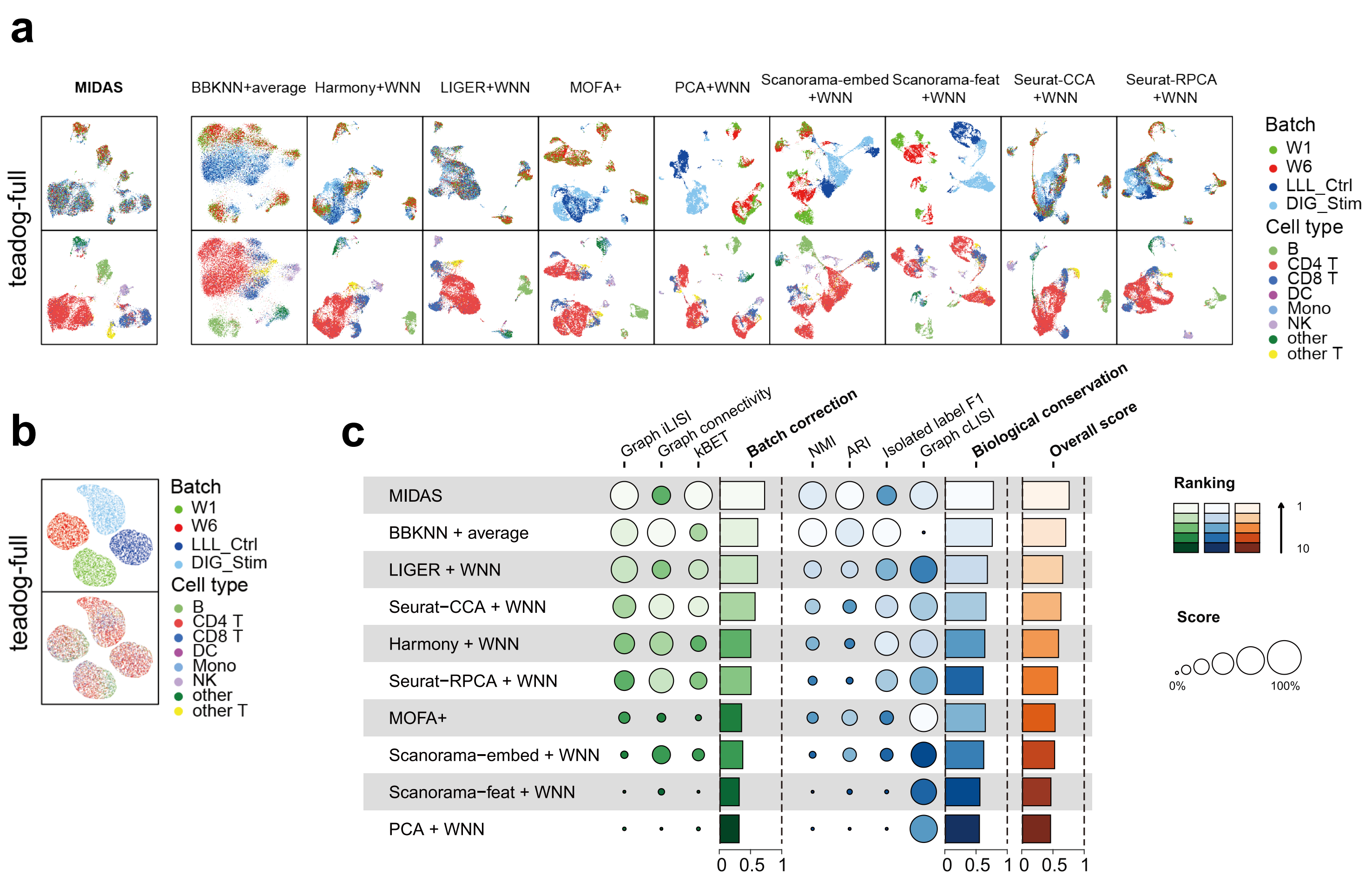

**Supplementary Figure 2. Performance of MIDAS on teadog-full dataset. a**, UMAP visualization of cell embeddings obtained by MIDAS and nine other strategies in the teadog-full datasets. The left two panels show inferred latent biological states and the right panels show dimensionality reduction results obtained with the other strategies. **b**, UMAP visualization of latent technical noise inferred by MIDAS in the teadog-full datasets. **c**, scIB benchmarking of performance on the teadog-full rectangular integration task.

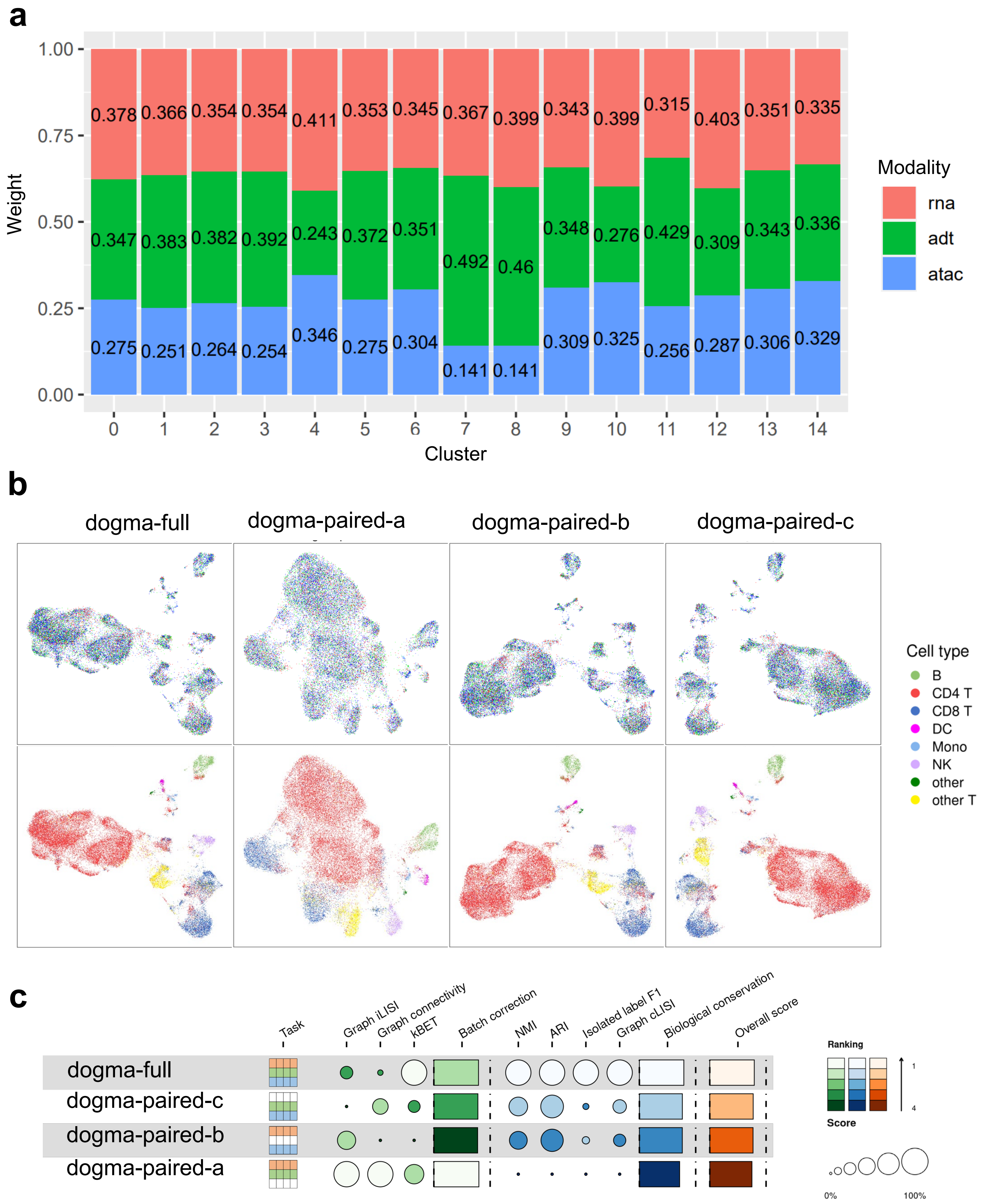

**Supplementary Figure 3. The contribution of different modalities to the joint clustering.** **a**, Contributions of RNA, ADT and ATAC to each cluster of DOGMA-full dataset. **b**, UMAP comparison of MIDAS embeddings on dogma-full, dogma-paired-a (remove ATAC modality), dogma-paired-b (remove RNA modality), and dogma-paired-c (remove ADT modality) integration tasks. In each task, cells in the top row are colored by batch, and cells in the bottom row are colored by cell type. **c**, scIB benchmarking of MIDAS performance on four tasks in (**b**). The removal of a modality significantly affects biological preservation, with the removal of ADT having the greatest impact.

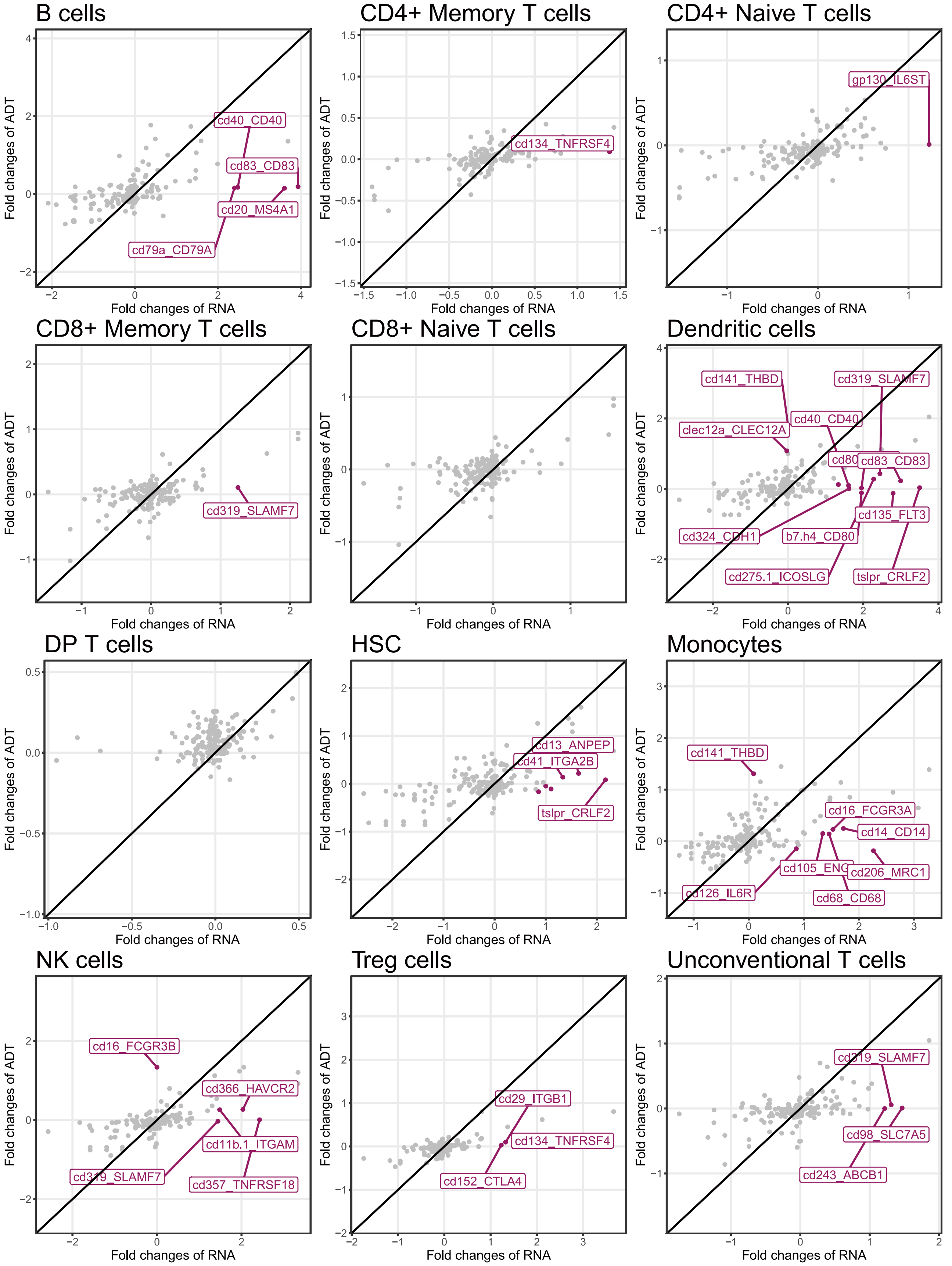

**Supplementary Figure 4. Expression inconsistencies between proteins and their corresponding genes in all cell types.**

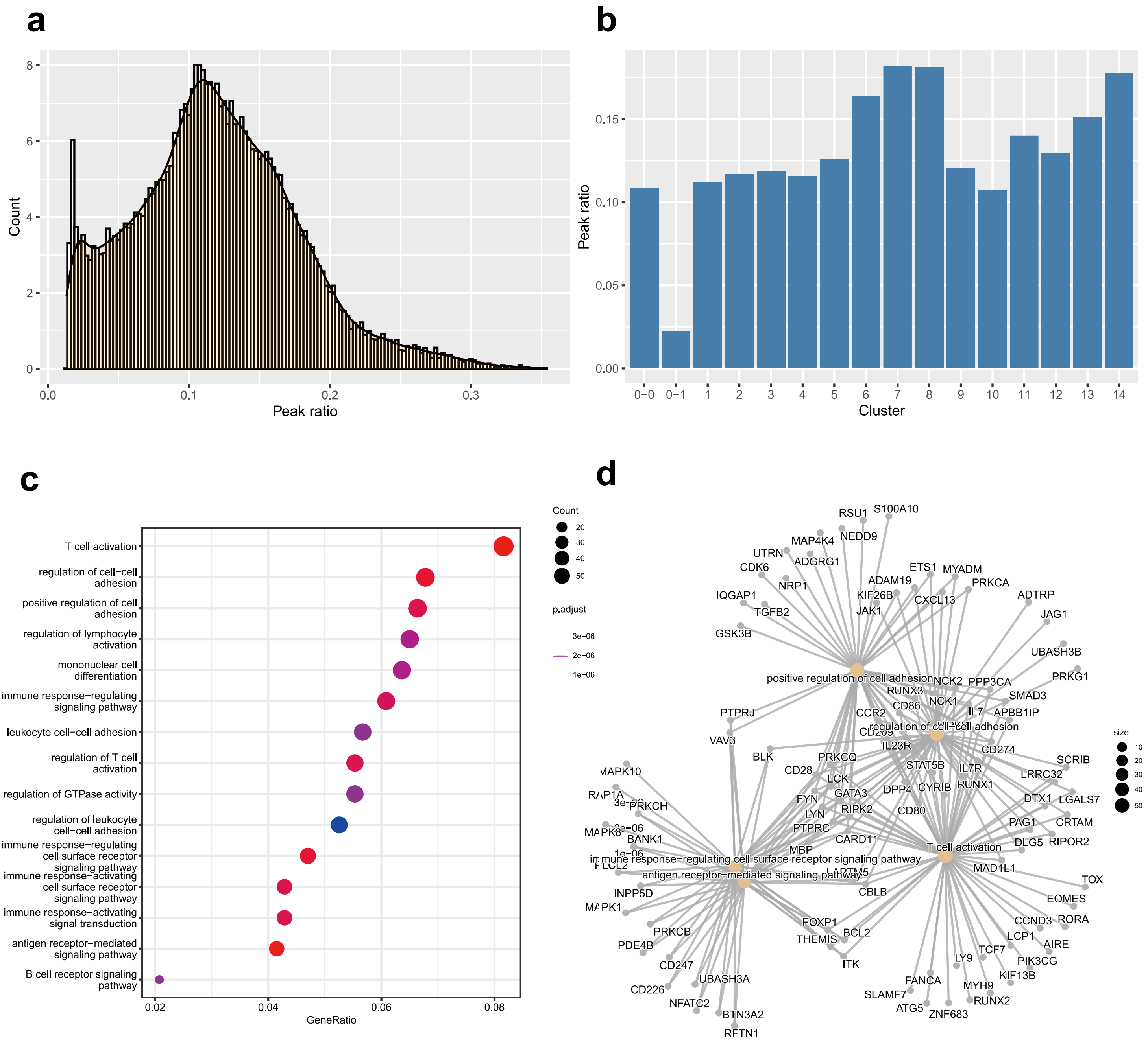

**Supplementary Figure 5. Cells with low chromatin accessibility. a**, ATAC peak ratio of all cells show a bi-modal distribution; Peak ratio of a cell is defined as the ratio of peak numbers of the cell to that of all the cells. **b**, ATAC peak ratio of cells in clusters. **c**, The Gene Ontology analysis for genes with deficient peak coverage. **d**, Linkages of genes with dificient peak coverage and biological concepts.

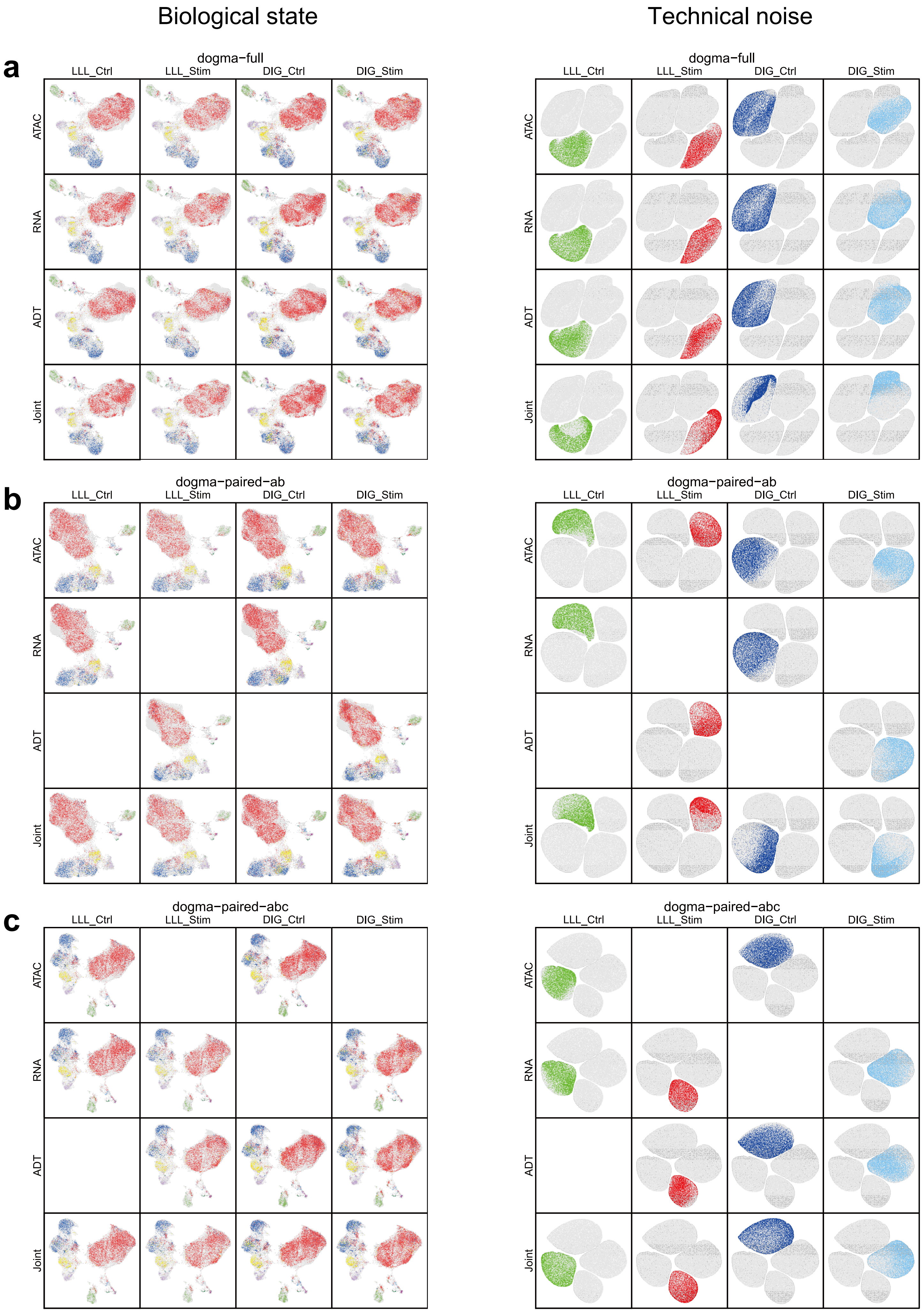
Biological state
Technical noise
a
dogma−full
LLL_Ctrl
LLL_Stim
DIG_Ctrl
DIG_Stim
ATAC
RNA
ADT
Joint
b
dogma−paired−ab
c
dogma−paired−abc

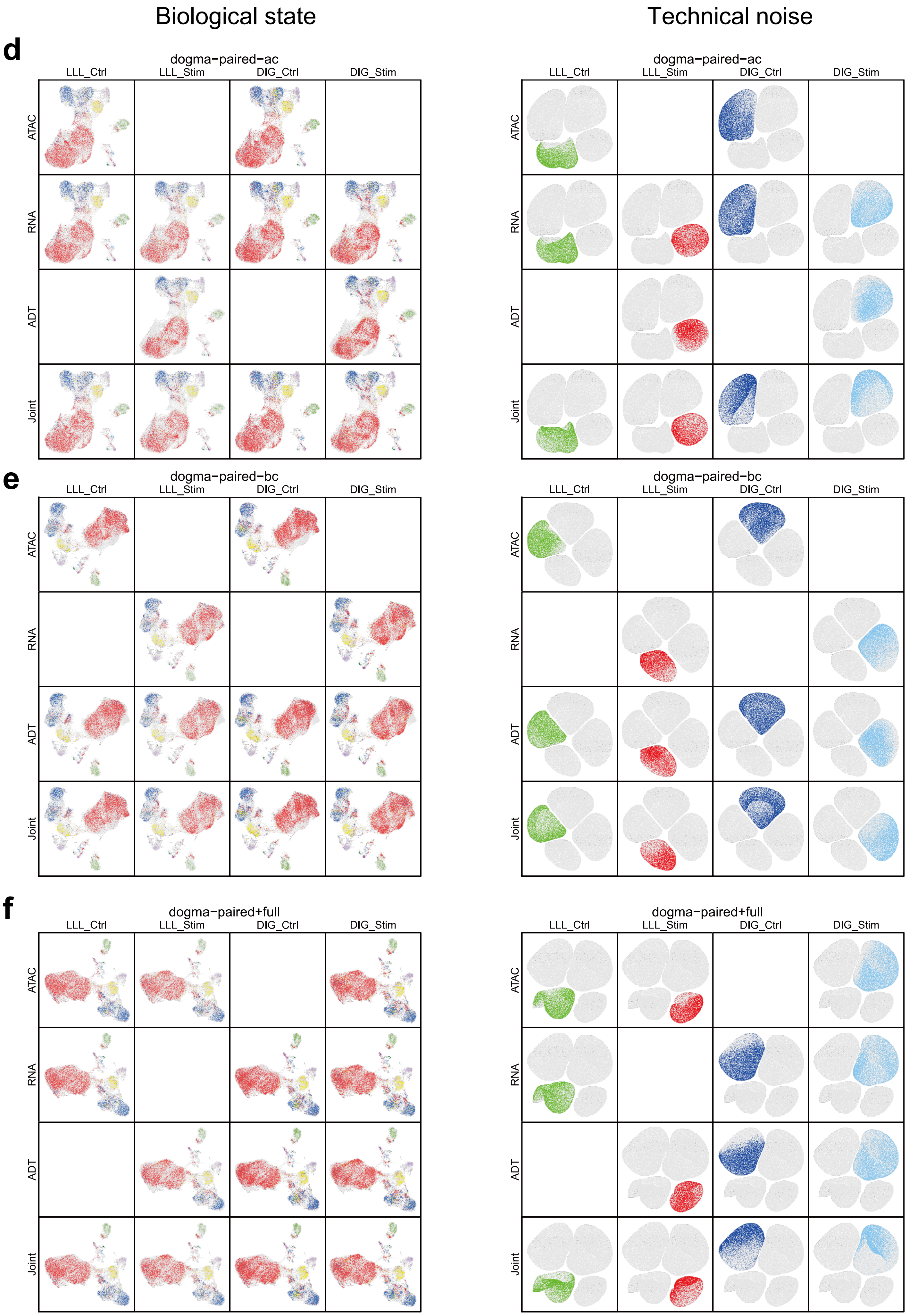
Biological state
Technical noise
d
dogma−paired−ac
e
dogma−paired−bc
f
dogma−paired+full
LLL_Ctrl
LLL_Stim
DIG_Ctrl
DIG_Stim
ATAC
RNA
ADT
Joint

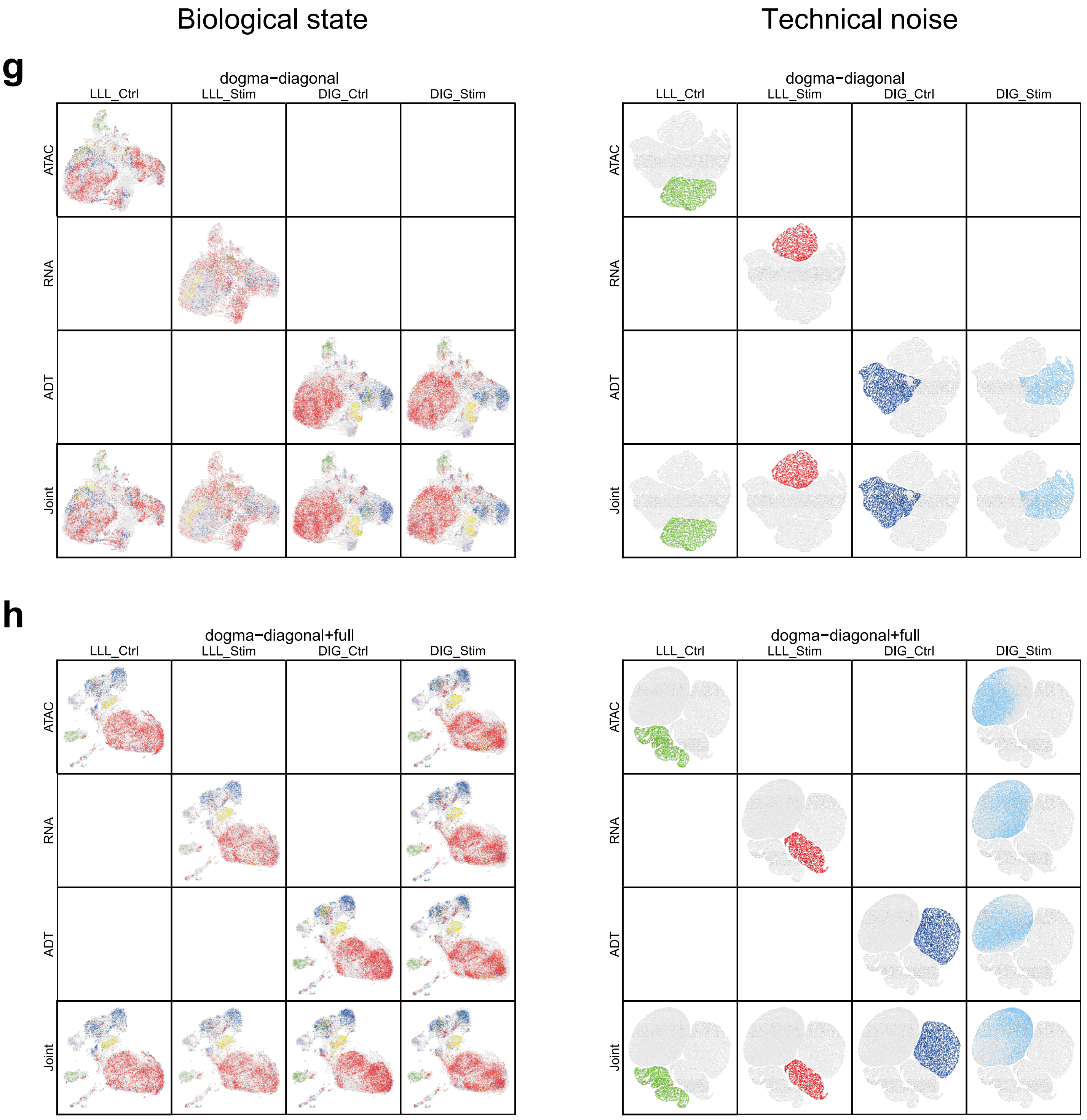

**Supplementary Figure 6. UMAP visualization of the biological states (left, colored by cell type) and technical noise (right, colored by batch) inferred by MIDAS with eight dogma mosaic datasets (a–h).**

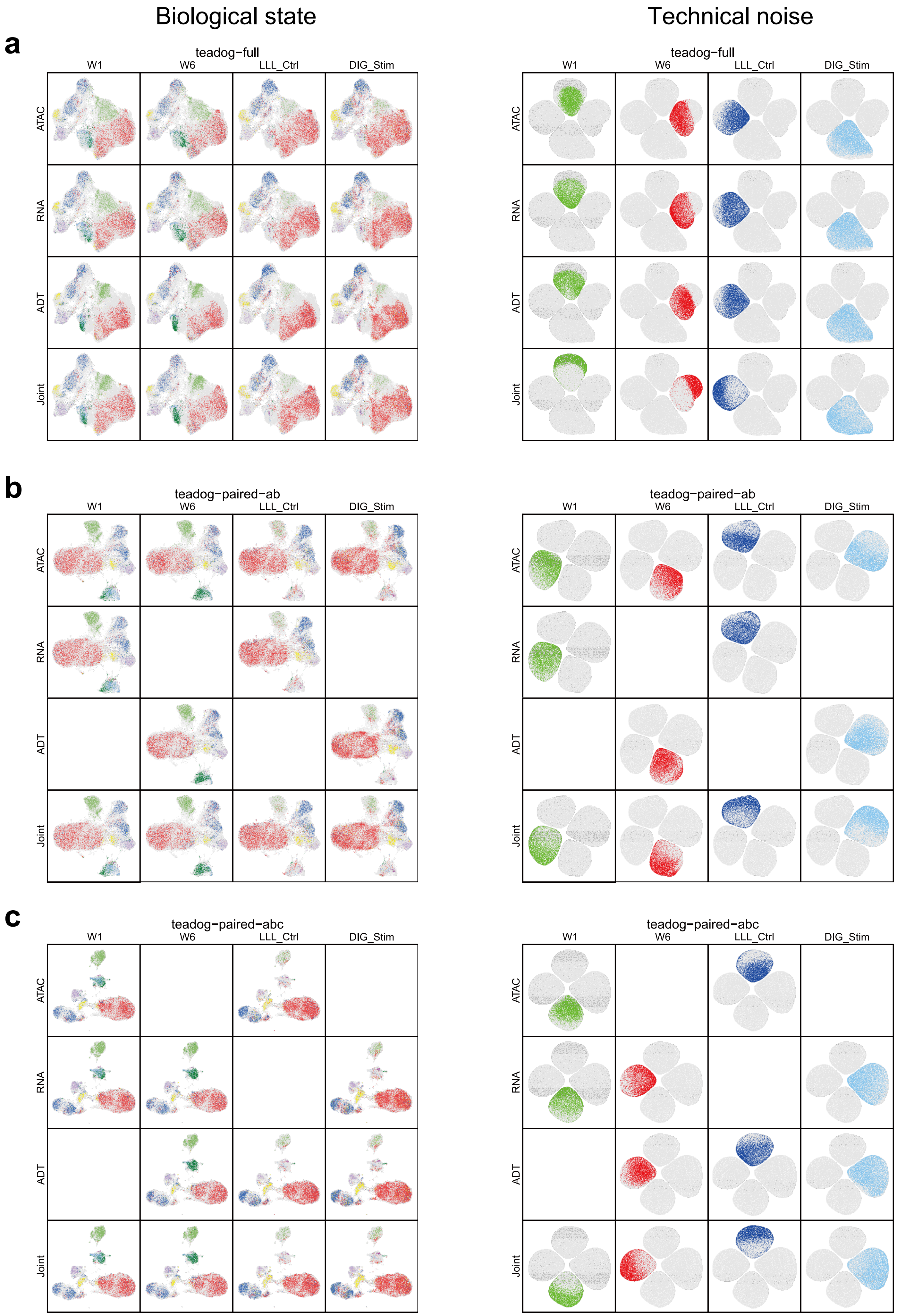

Biological state
Technical noise
a
teadog−full
W1
W6
LLL_Ctrl
DIG_Stim
ATAC
RNA
ADT
Joint
b
teadog−paired−ab
c
teadog−paired−abc

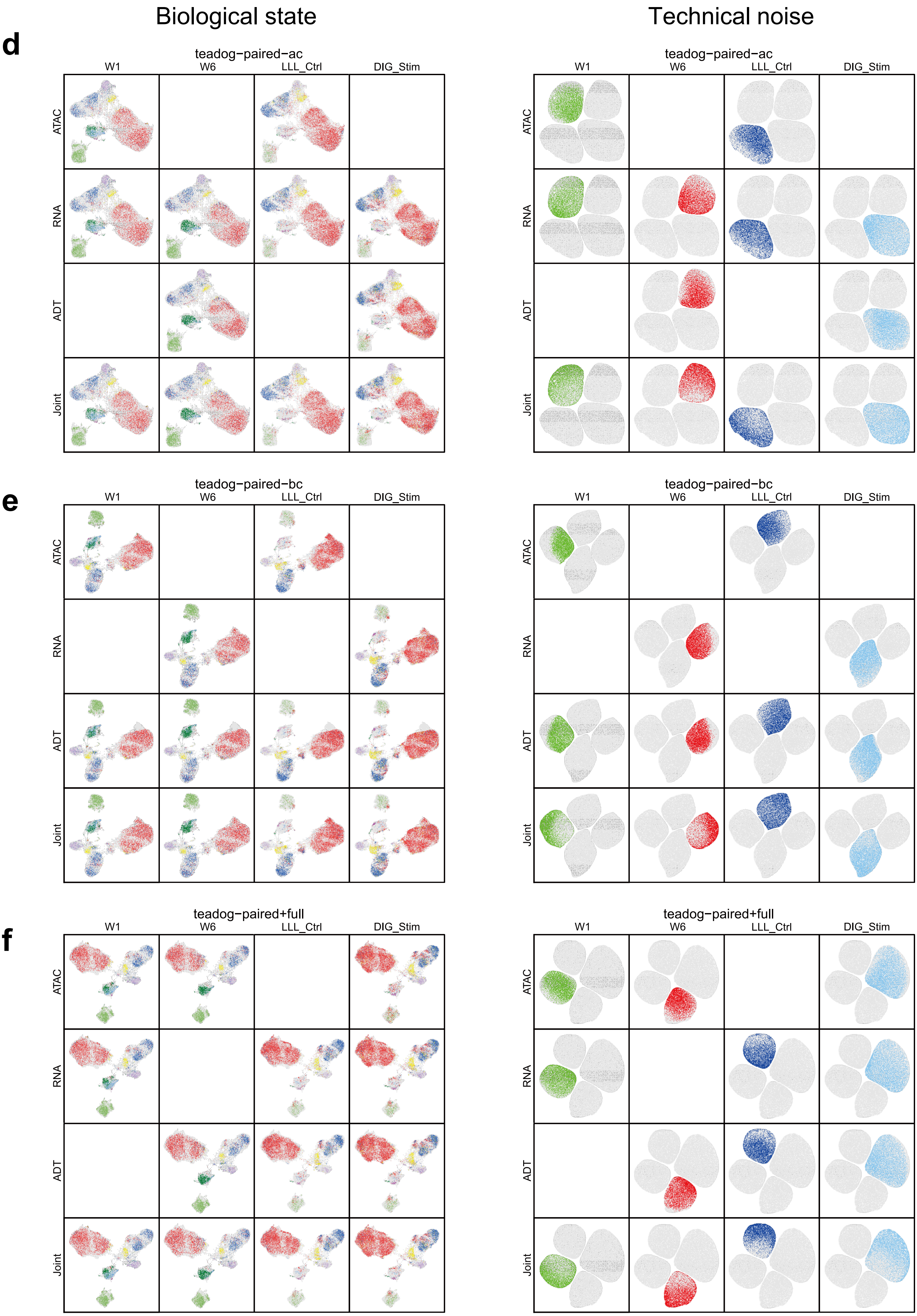
Biological state
Technical noise
d
teadog−paired−ac
W1
W6
LLL_Ctrl
DIG_Stim
ATAC
RNA
ADT
Joint
e
teadog−paired−bc
f
teadog−paired+full

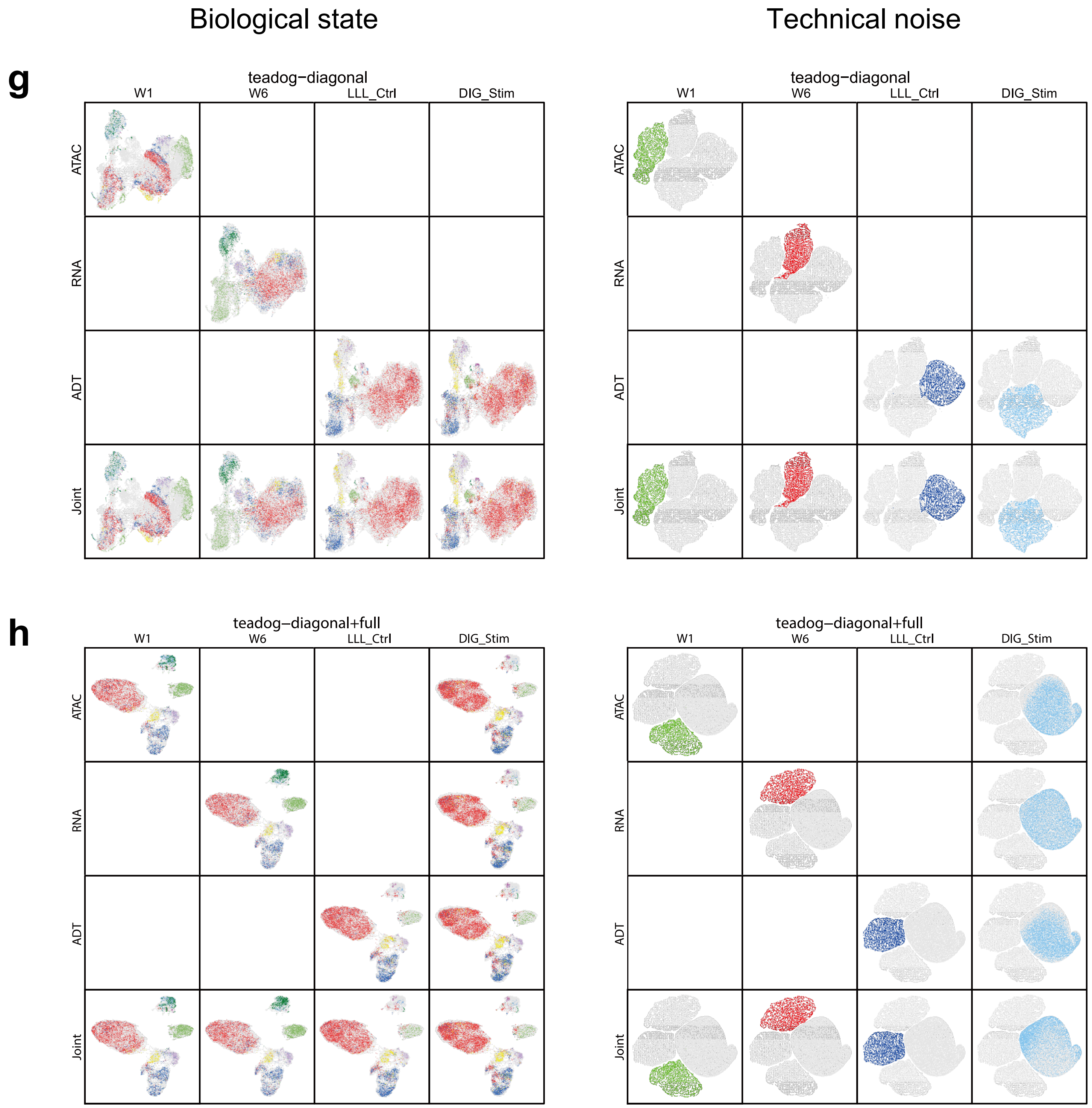

**Supplementary Figure 7. UMAP visualization of the biological states (left, colored by cell type) and technical noise (right, colored by batch) inferred by MIDAS with eight teadog mosaic datasets (a–h).**

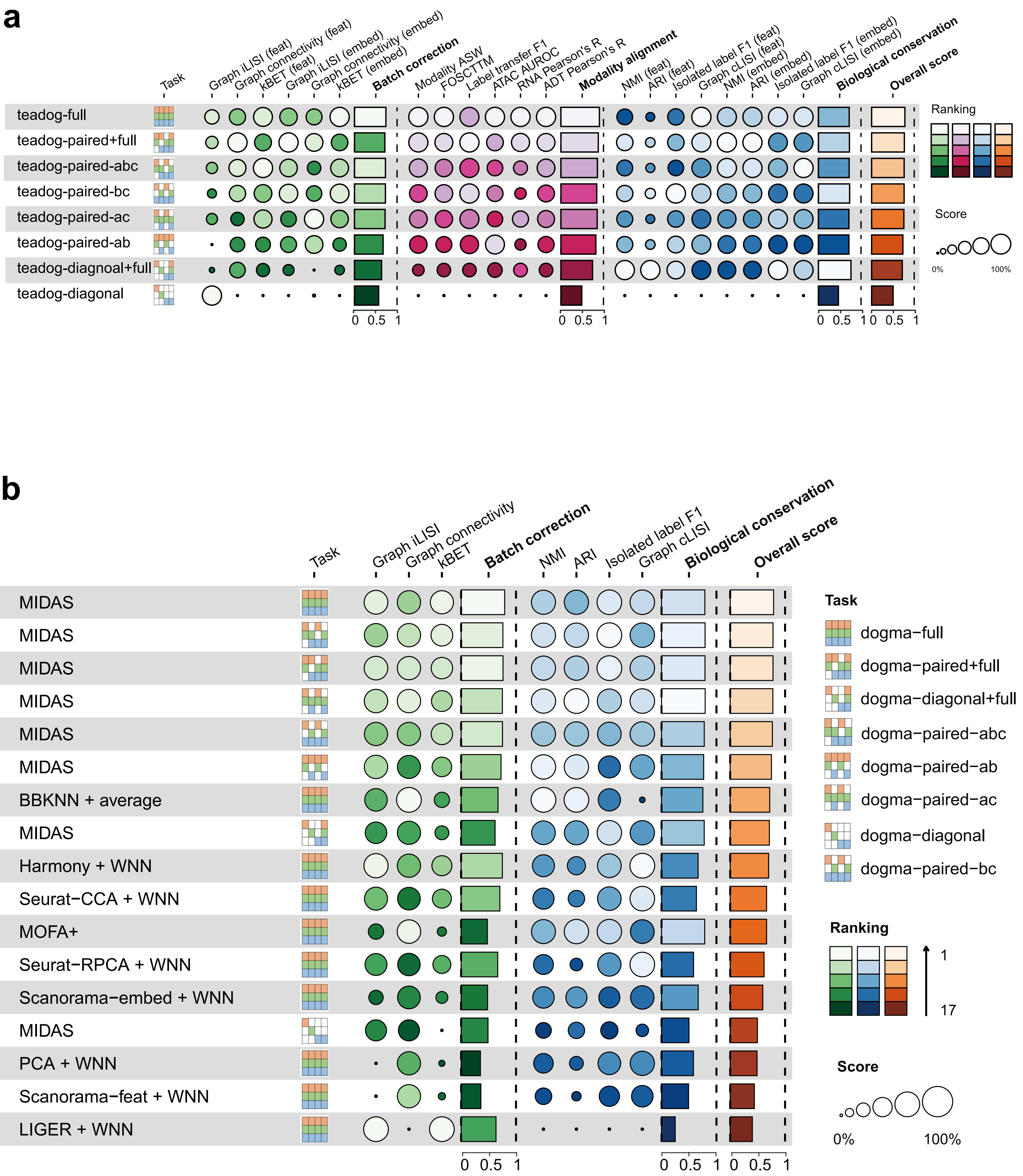

a
Task
Graph iLISI (feat)
Graph connectivity (feat)
kBET (feat)
Graph iLISI (embed)
Graph connectivity (embed)
kBET (embed)
Batch correction
Modality ASW
FOSCTTM
Label transfer F1
ATAC AUROC
RNA Pearson's R
ADT Pearson's R
Modality alignment
NMI (feat)
ARI (feat)
Isolated label F1 (feat)
Graph cLISI (feat)
NMI (embed)
ARI (embed)
Isolated label F1 (embed)
Graph cLISI (embed)
Biological conservation
Overall score
teadog-full
teadog-paired+full
teadog-paired-abc
teadog-paired-bc
teadog-paired-ac
teadog-paired-ab
teadog-diagnoal+full
teadog-diagonal
Ranking
Score
0%
100%
0 0.5 1
0 0.5 1
0 0.5 1
0 0.5 1
b
Task
Graph iLISI
Graph connectivity
kBET
Batch correction
NMI
ARI
Isolated label F1
Graph cLISI
Biological conservation
Overall score
MIDAS
MIDAS
MIDAS
MIDAS
MIDAS
MIDAS
BBKNN + average
MIDAS
Harmony + WNN
Seurat−CCA + WNN
MOFA+
Seurat−RPCA + WNN
Scanorama−embed + WNN
MIDAS
PCA + WNN
Scanorama−feat + WNN
LIGER + WNN
Task
dogma−full
dogma−paired+full
dogma−diagonal+full
dogma−paired−abc
dogma−paired−ab
dogma−paired−ac
dogma−diagonal
dogma−paired−bc
Ranking
1
17
Score
0%
100%
0 0.5 1
0 0.5 1
0 0.5 1

c

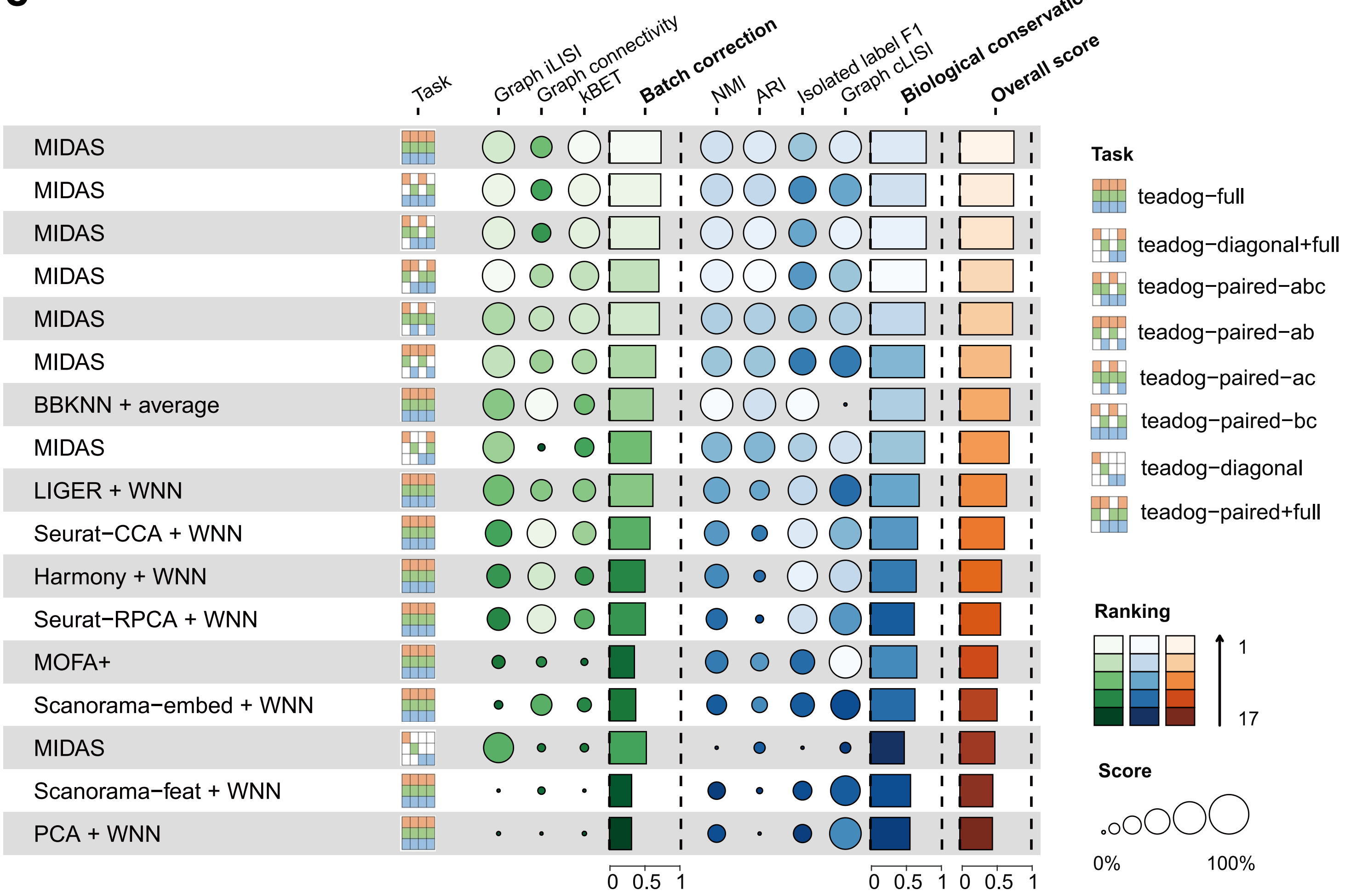

**Supplementary Figure 8. scIB and scMIB benchmarking of the performance of MIDAS and other strategies. a**, Benchmarking of MIDAS performance on teadog mosaic integration tasks using our proposed scMIB. **b**, scIB benchmarking of MIDAS performance on dogma mosaic integration tasks against nine other strategies on the dogma-full rectangular integration task. **c**, scIB benchmarking of MIDAS performance on teadog mosaic integration tasks against nine other strategies on the teadog-full rectangular integration task.

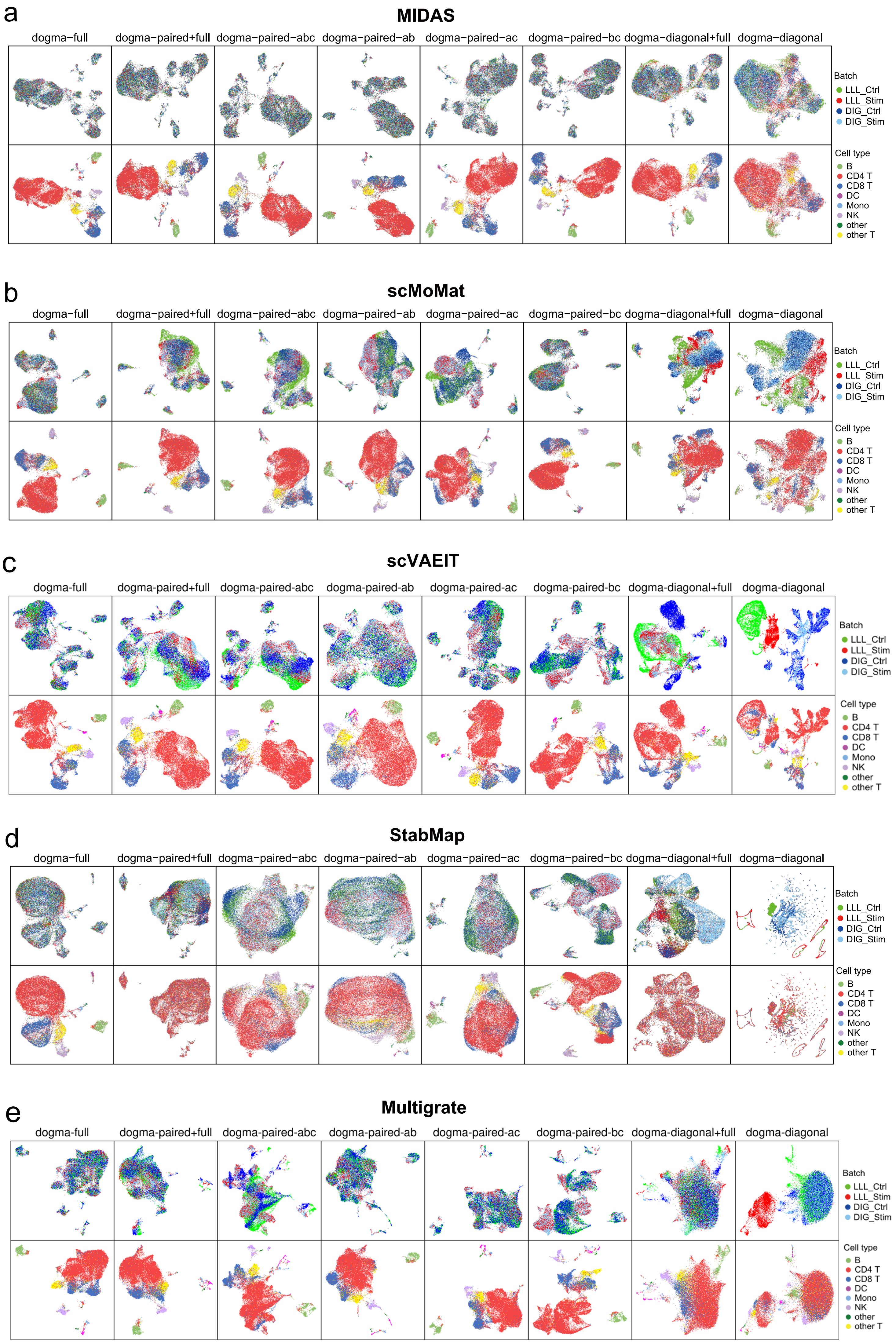
a
MIDAS
dogma−full
dogma−paired+full
dogma−paired−abc
dogma−paired−ab
dogma−paired−ac
dogma−paired−bc
dogma−diagonal+full
dogma−diagonal
Batch
LLL_Ctrl
LLL_Stim
DIG_Ctrl
DIG_Stim
Cell type
B
CD4 T
CD8 T
DC
Mono
NK
other
other T
b
scMoMat
dogma−full
dogma−paired+full
dogma−paired−abc
dogma−paired−ab
dogma−paired−ac
dogma−paired−bc
dogma−diagonal+full
dogma−diagonal
Batch
LLL_Ctrl
LLL_Stim
DIG_Ctrl
DIG_Stim
Cell type
B
CD4 T
CD8 T
DC
Mono
NK
other
other T
c
scVAEIT
dogma-full
dogma-paired+full
dogma-paired-abc
dogma-paired-ab
dogma-paired-ac
dogma-paired-bc
dogma-diagonal+full
dogma-diagonal
Batch
LLL_Ctrl
LLL_Stim
DIG_Ctrl
DIG_Stim
Cell type
B
CD4 T
CD8 T
DC
Mono
NK
other
other T
d
StabMap
dogma−full
dogma−paired+full
dogma−paired−abc
dogma−paired−ab
dogma−paired−ac
dogma−paired−bc
dogma−diagonal+full
dogma−diagonal
Batch
LLL_Ctrl
LLL_Stim
DIG_Ctrl
DIG_Stim
Cell type
B
CD4 T
CD8 T
DC
Mono
NK
other
other T
e
Multigrate
dogma-full
dogma-paired+full
dogma-paired-abc
dogma-paired-ab
dogma-paired-ac
dogma-paired-bc
dogma-diagonal+full
dogma-diagonal
Batch
LLL_Ctrl
LLL_Stim
DIG_Ctrl
DIG_Stim
Cell type
B
CD4 T
CD8 T
DC
Mono
NK
other
other T

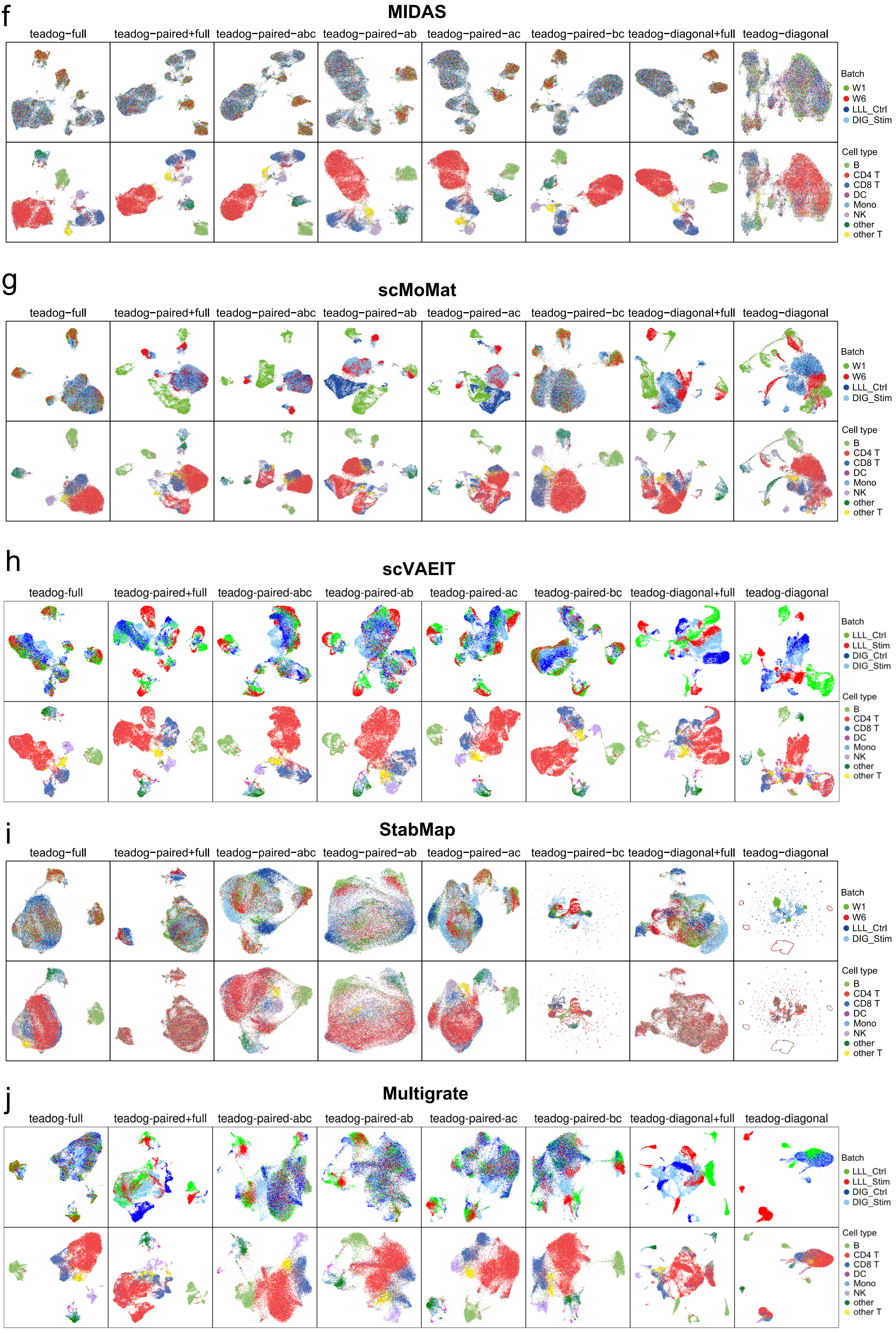
f
MIDAS
teadog-full
teadog-paired+full
teadog-paired-abc
teadog-paired-ab
teadog-paired-ac
teadog-paired-bc
teadog-diagonal+full
teadog-diagonal
Batch
W1
W6
LLL_Ctrl
DIG_Stim
Cell type
B
CD4 T
CD8 T
DC
Mono
NK
other
other T
g
scMoMat
teadog-full
teadog-paired+full
teadog-paired-abc
teadog-paired-ab
teadog-paired-ac
teadog-paired-bc
teadog-diagonal+full
teadog-diagonal
Batch
W1
W6
LLL_Ctrl
DIG_Stim
Cell type
B
CD4 T
CD8 T
DC
Mono
NK
other
other T
h
scVAEIT
teadog-full
teadog-paired+full
teadog-paired-abc
teadog-paired-ab
teadog-paired-ac
teadog-paired-bc
teadog-diagonal+full
teadog-diagonal
Batch
LLL_Ctrl
LLL_Stim
DIG_Ctrl
DIG_Stim
Cell type
B
CD4 T
CD8 T
DC
Mono
NK
other
other T
i
StabMap
teadog-full
teadog-paired+full
teadog-paired-abc
teadog-paired-ab
teadog-paired-ac
teadog-paired-bc
teadog-diagonal+full
teadog-diagonal
Batch
W1
W6
LLL_Ctrl
DIG_Stim
Cell type
B
CD4 T
CD8 T
DC
Mono
NK
other
other T
j
Multigrate
teadog-full
teadog-paired+full
teadog-paired-abc
teadog-paired-ab
teadog-paired-ac
teadog-paired-bc
teadog-diagonal+full
teadog-diagonal
Batch
LLL_Ctrl
LLL_Stim
DIG_Ctrl
DIG_Stim
Cell type
B
CD4 T
CD8 T
DC
Mono
NK
other
other T

**Supplementary Figure 9. UMAP comparison of embeddings on dogma (a–e) and teadog (f–j) mosaic integration tasks.** In each task, cells in the top row are colored by batch, and cells in the bottom row are colored by cell type.

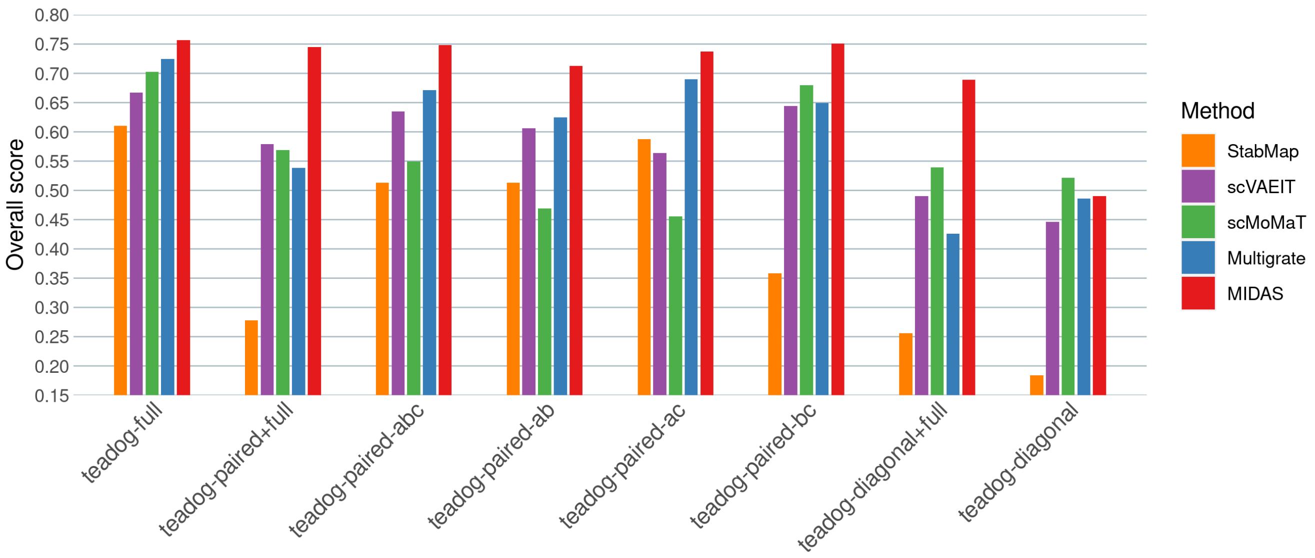

**Supplementary Figure 10. Comparison of scIB overall scores on teadog mosaic integration tasks.**

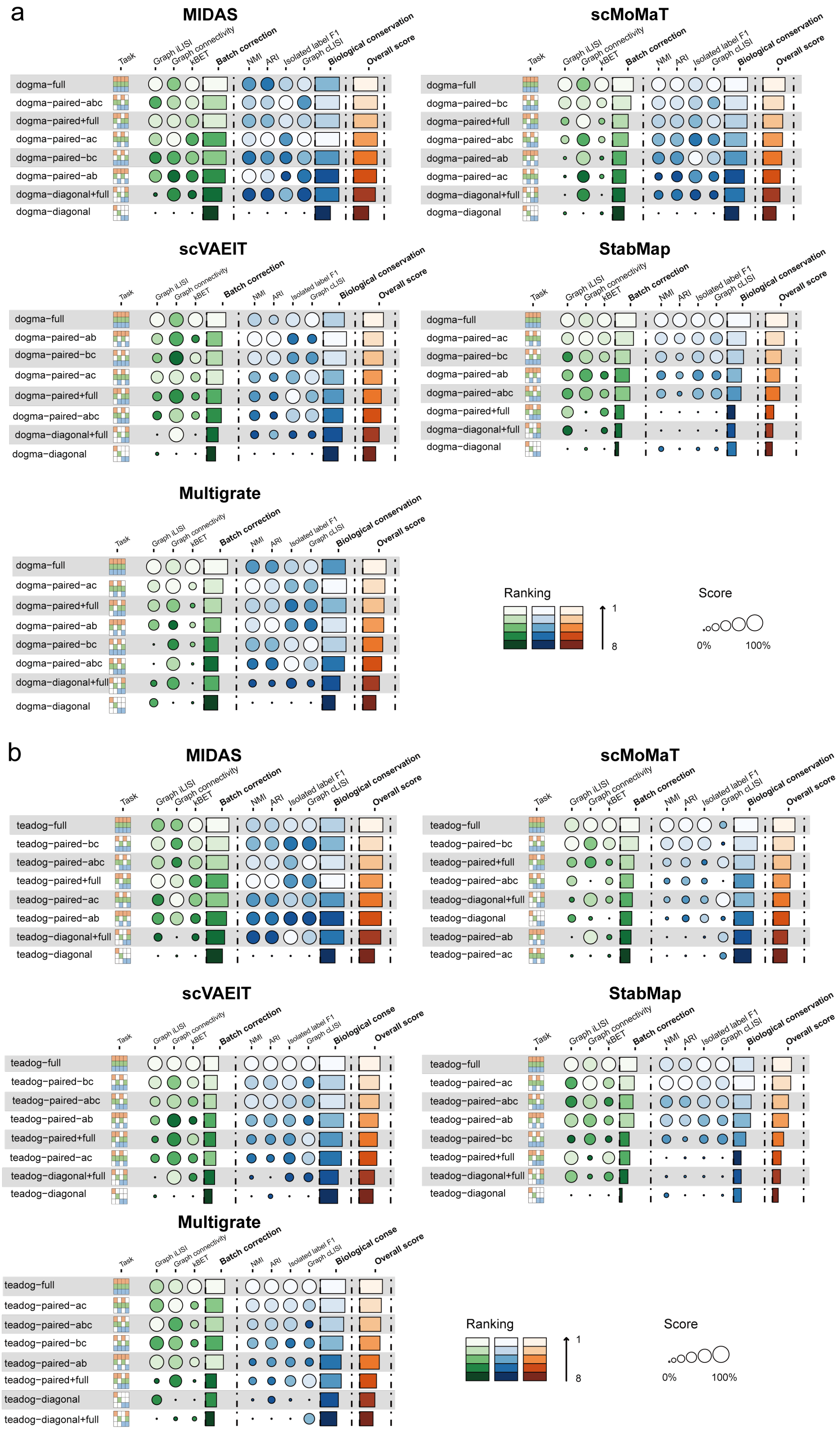

**Supplementary Figure 11. Detailed comparison of scIB scores on dogma (a) and teadog (b) mosaic integration tasks.**

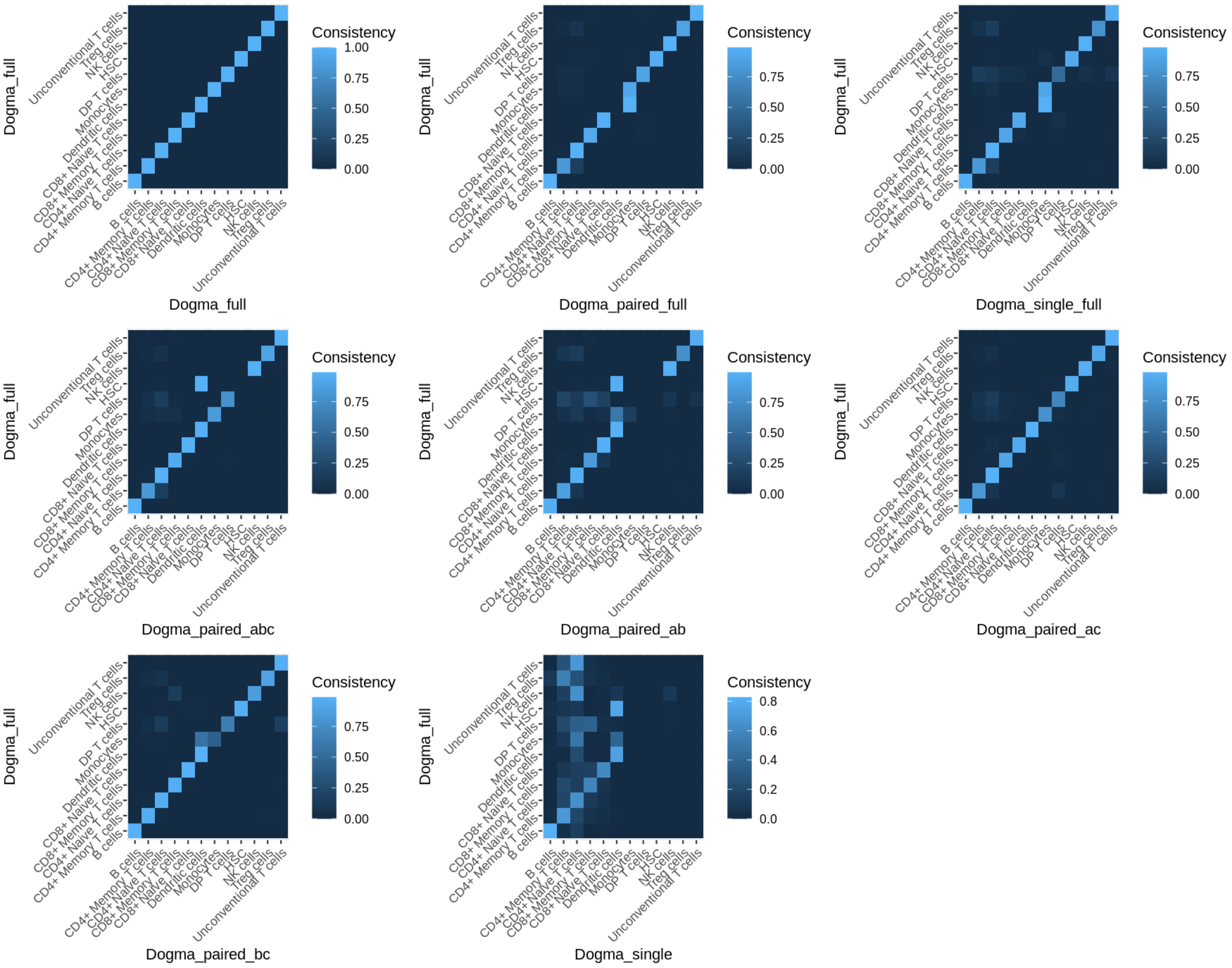

**Supplementary Figure 12. Consistency of downstream labelling results got from different mosaic integration tasks.** Each task result is compared to the dogma-full task result.

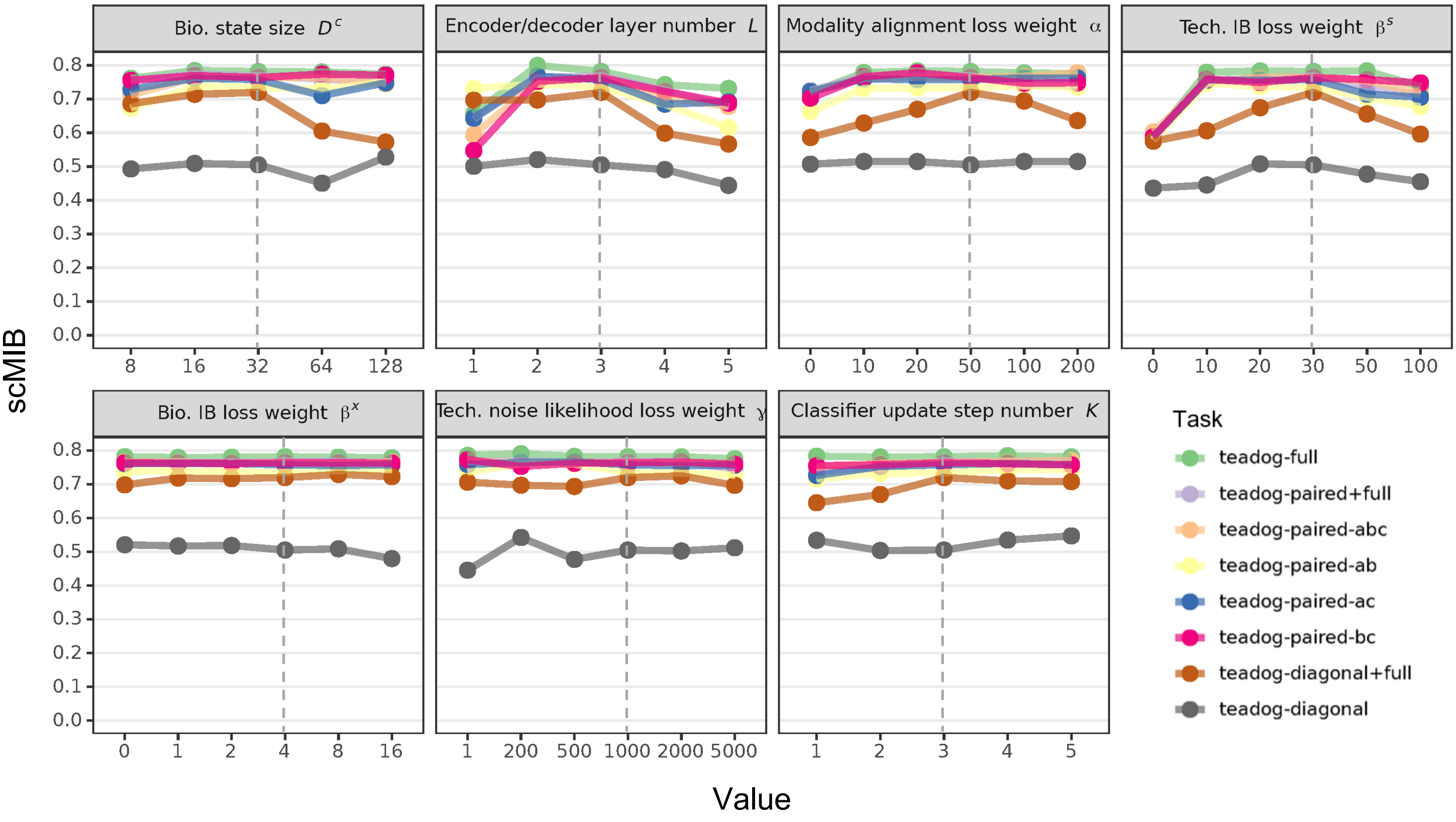

**Supplementary Figure 13. Integration performance of MIDAS under different settings of hyperparameters.** In each plot, the dashed line corresponds to the default value. To control computational costs, we only change one hyperparameter at a time while setting the others to their default values. The value "0" on the x-axis corresponds to the disability of the modality alignment module, technical information bottleneck module, and biological information bottleneck module, respectively. In the "Technical noise likelihood coefficient $\gamma$" panel, the value "1" on the x-axis represents the disability of the technical noise module. The teadog mosaic datasets were chosen as evaluation datasets, which are more challenging and compelling than the dogma mosaic datasets for highlighting the impact of changes in hyperparameters.

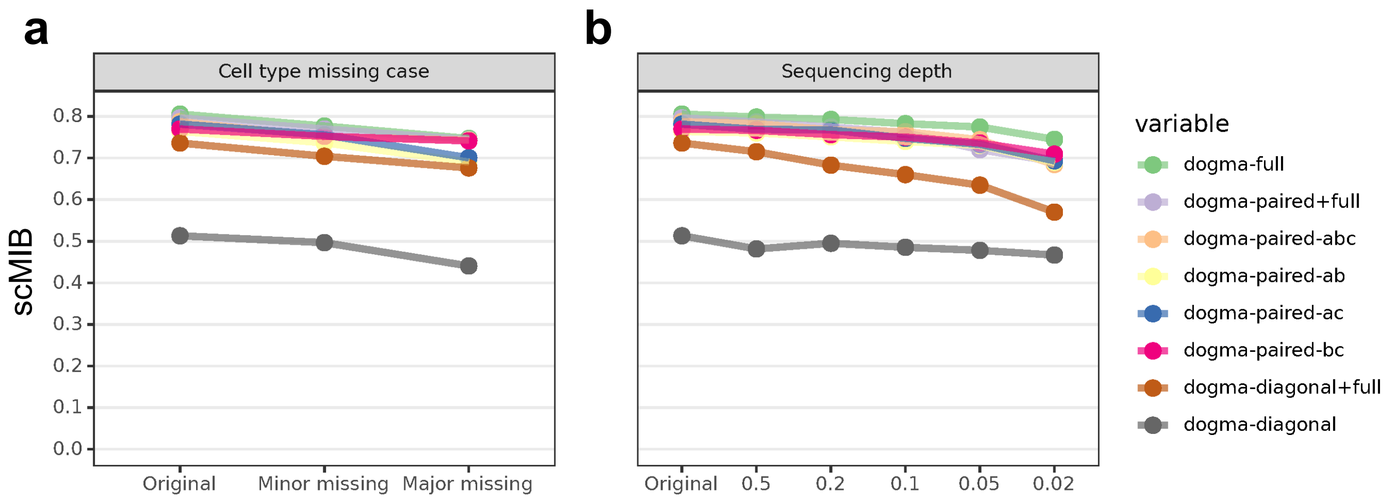

**Supplementary Figure 14. Integration performance of MIDAS under different dataset variations including cell type missing (a) and sequencing depth changing (b).** The dogma mosaic datasets were chosen as evaluation datasets, which exhibit fewer biological and technical variations across batches compared to the teadog mosaic datasets. This selection will be helpful in controlling the variation factors in the dataset, allowing for a clearer understanding of the impact of these changes.

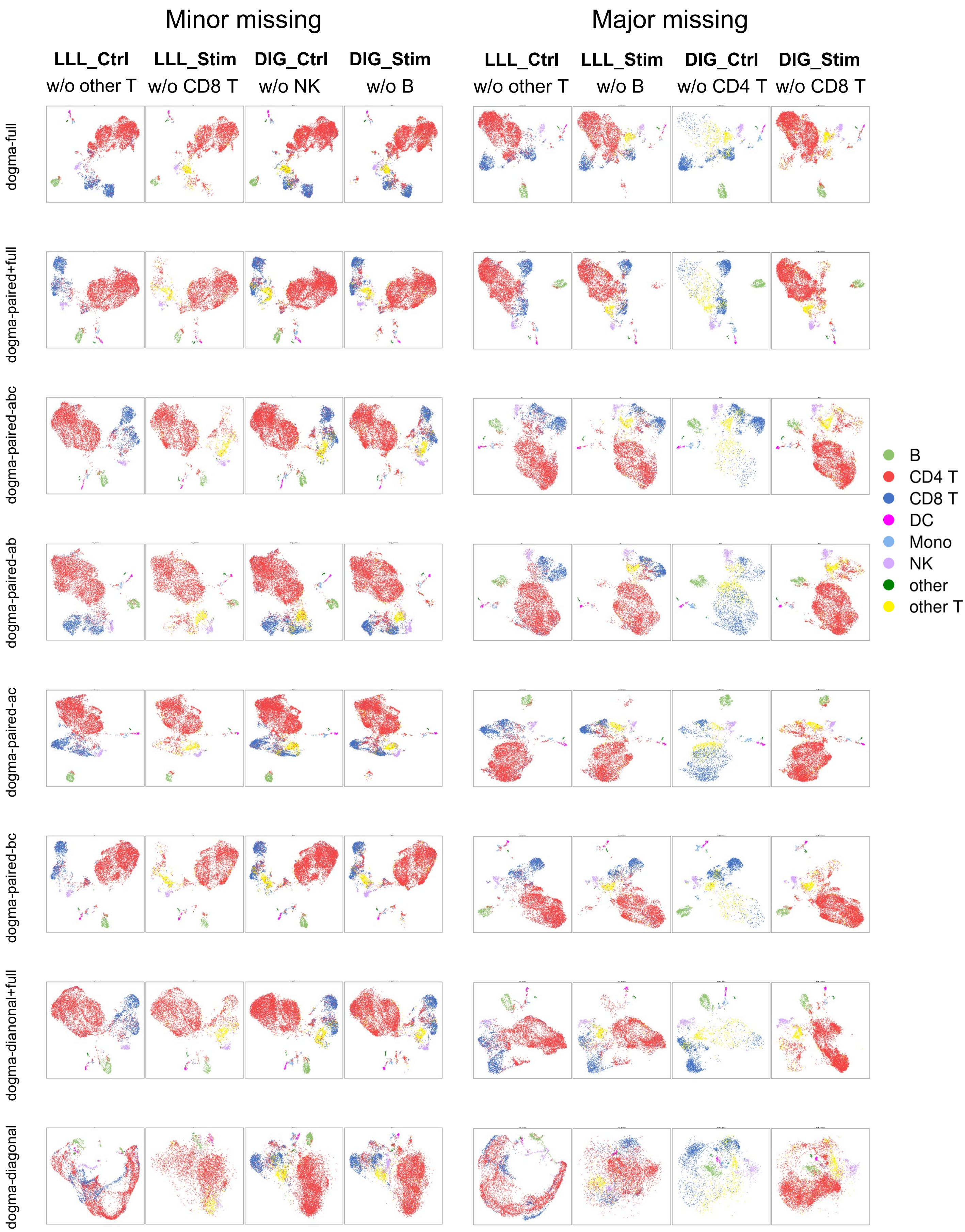

**Supplementary Figure 15. UMAP comparison of MIDAS embeddings on dogma mosaic integration tasks with minor (left) and major (right) cell type missing.**

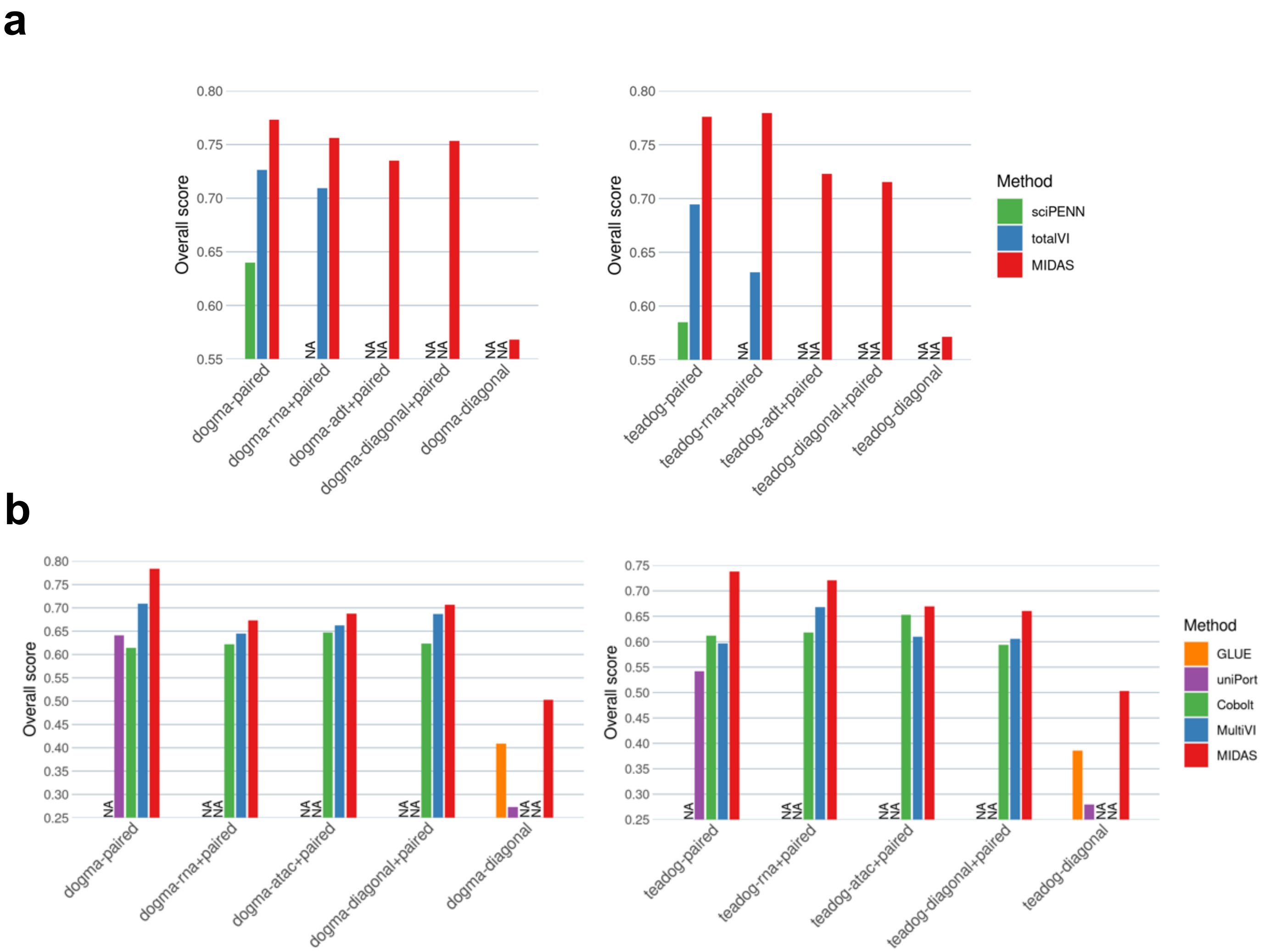

**Supplementary Figure 16. Comparison of scIB overall scores on bimodal mosaic integration tasks. a,** scIB overall scores of MIDAS, sciPENN and totalVI on RNA+ADT integration tasks based on dogma (left panel) and teadog (right panel) datasets. **b,** scIB overall scores of MIDAS, Cobolt, GLUE, multiVI and uniPort on ATAC+RNA integration tasks based on dogma (left panel) and teadog (right panel) datasets.

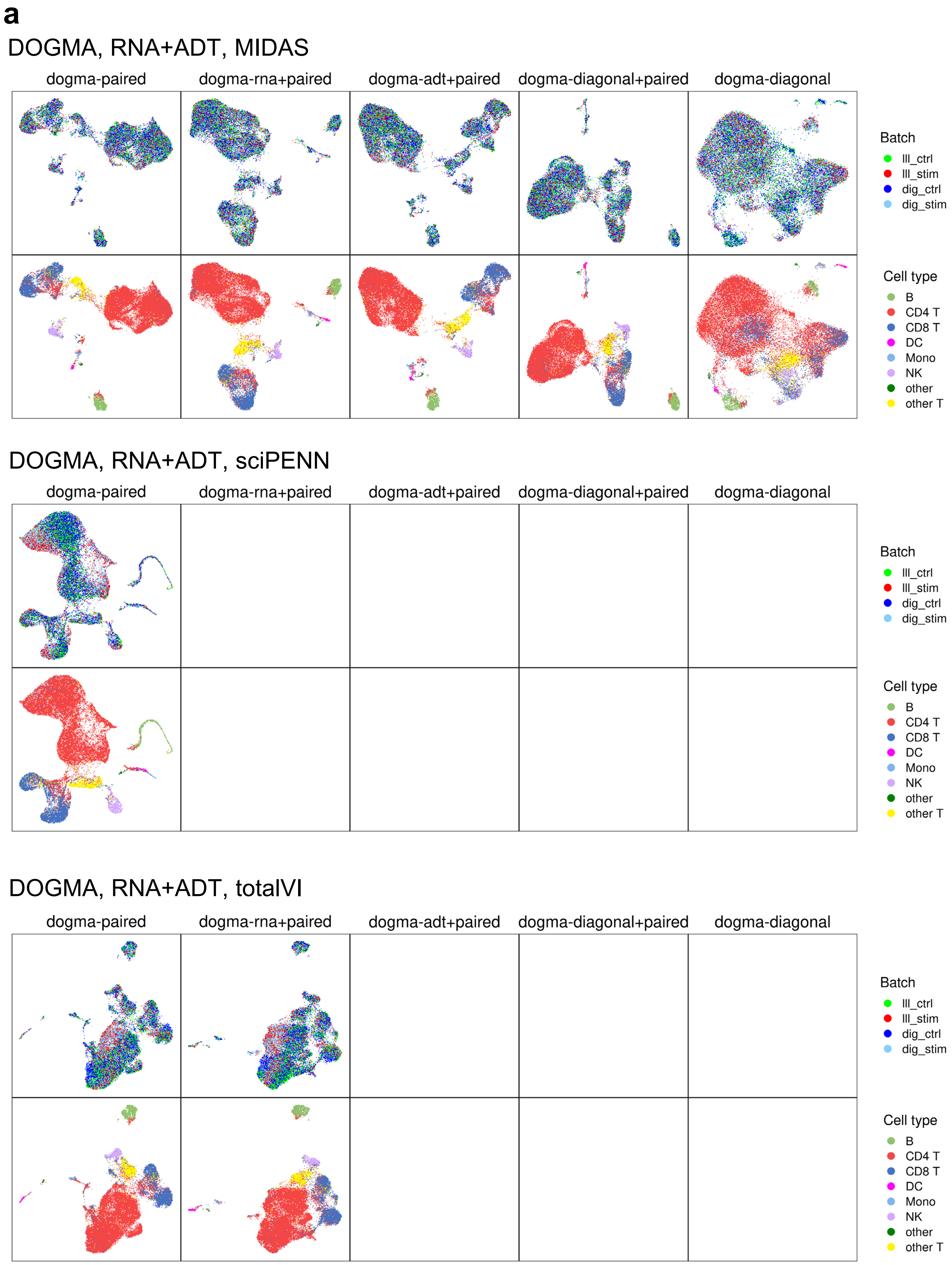
a
DOGMA, RNA+ADT, MIDAS
dogma-paired
dogma-rna+paired
dogma-adt+paired
dogma-diagonal+paired
dogma-diagonal
Batch
lll_ctrl
lll_stim
dig_ctrl
dig_stim
Cell type
B
CD4 T
CD8 T
DC
Mono
NK
other
other T
DOGMA, RNA+ADT, sciPENN
dogma-paired
dogma-rna+paired
dogma-adt+paired
dogma-diagonal+paired
dogma-diagonal
Batch
lll_ctrl
lll_stim
dig_ctrl
dig_stim
Cell type
B
CD4 T
CD8 T
DC
Mono
NK
other
other T
DOGMA, RNA+ADT, totalVI
dogma-paired
dogma-rna+paired
dogma-adt+paired
dogma-diagonal+paired
dogma-diagonal
Batch
lll_ctrl
lll_stim
dig_ctrl
dig_stim
Cell type
B
CD4 T
CD8 T
DC
Mono
NK
other
other T

**b**

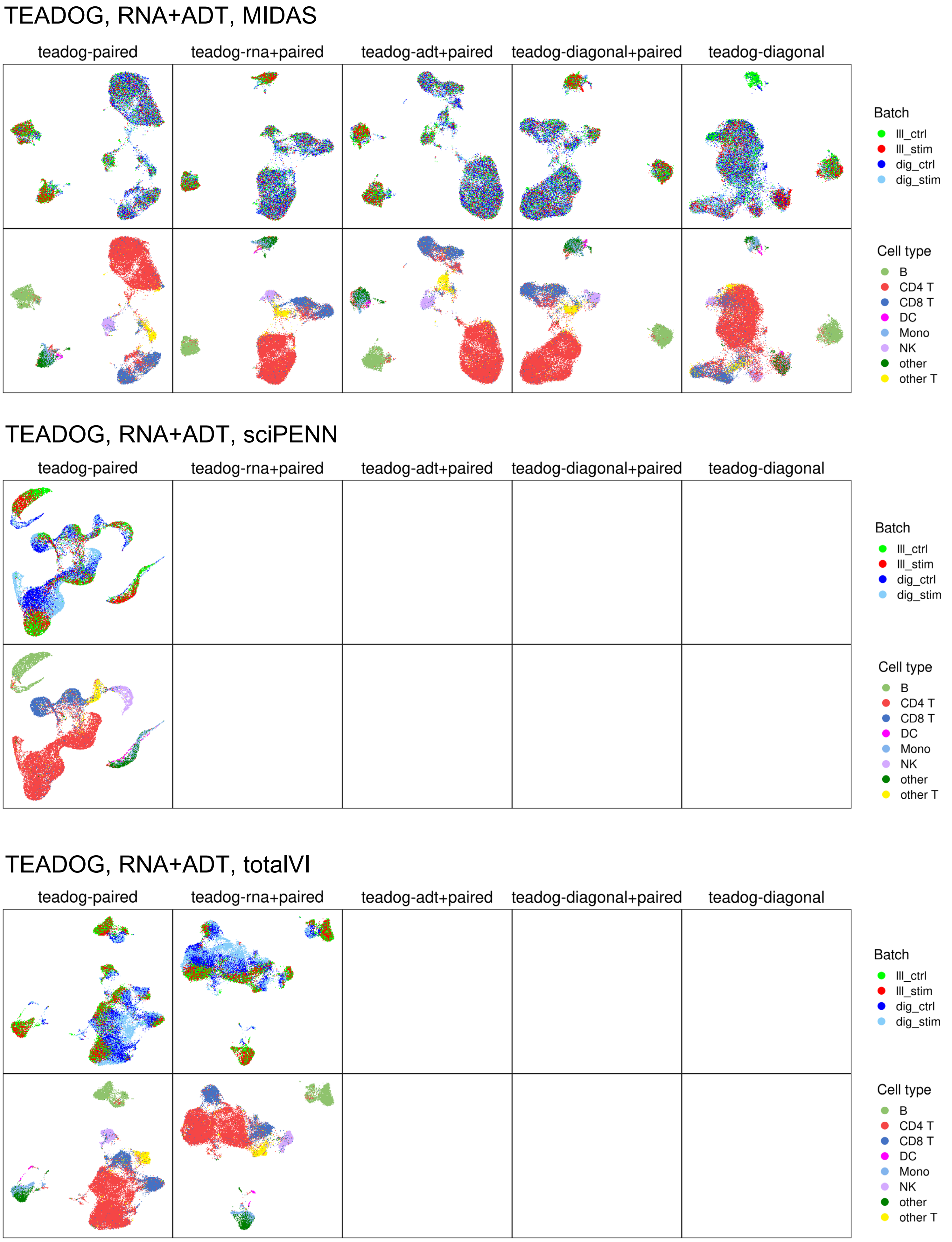
TEADOG, RNA+ADT, MIDAS
teadog-paired
teadog-rna+paired
teadog-adt+paired
teadog-diagonal+paired
teadog-diagonal
Batch
lll_ctrl
lll_stim
dig_ctrl
dig_stim
Cell type
B
CD4 T
CD8 T
DC
Mono
NK
other
other T
TEADOG, RNA+ADT, sciPENN
teadog-paired
teadog-rna+paired
teadog-adt+paired
teadog-diagonal+paired
teadog-diagonal
Batch
lll_ctrl
lll_stim
dig_ctrl
dig_stim
Cell type
B
CD4 T
CD8 T
DC
Mono
NK
other
other T
TEADOG, RNA+ADT, totalVI
teadog-paired
teadog-rna+paired
teadog-adt+paired
teadog-diagonal+paired
teadog-diagonal
Batch
lll_ctrl
lll_stim
dig_ctrl
dig_stim
Cell type
B
CD4 T
CD8 T
DC
Mono
NK
other
other T

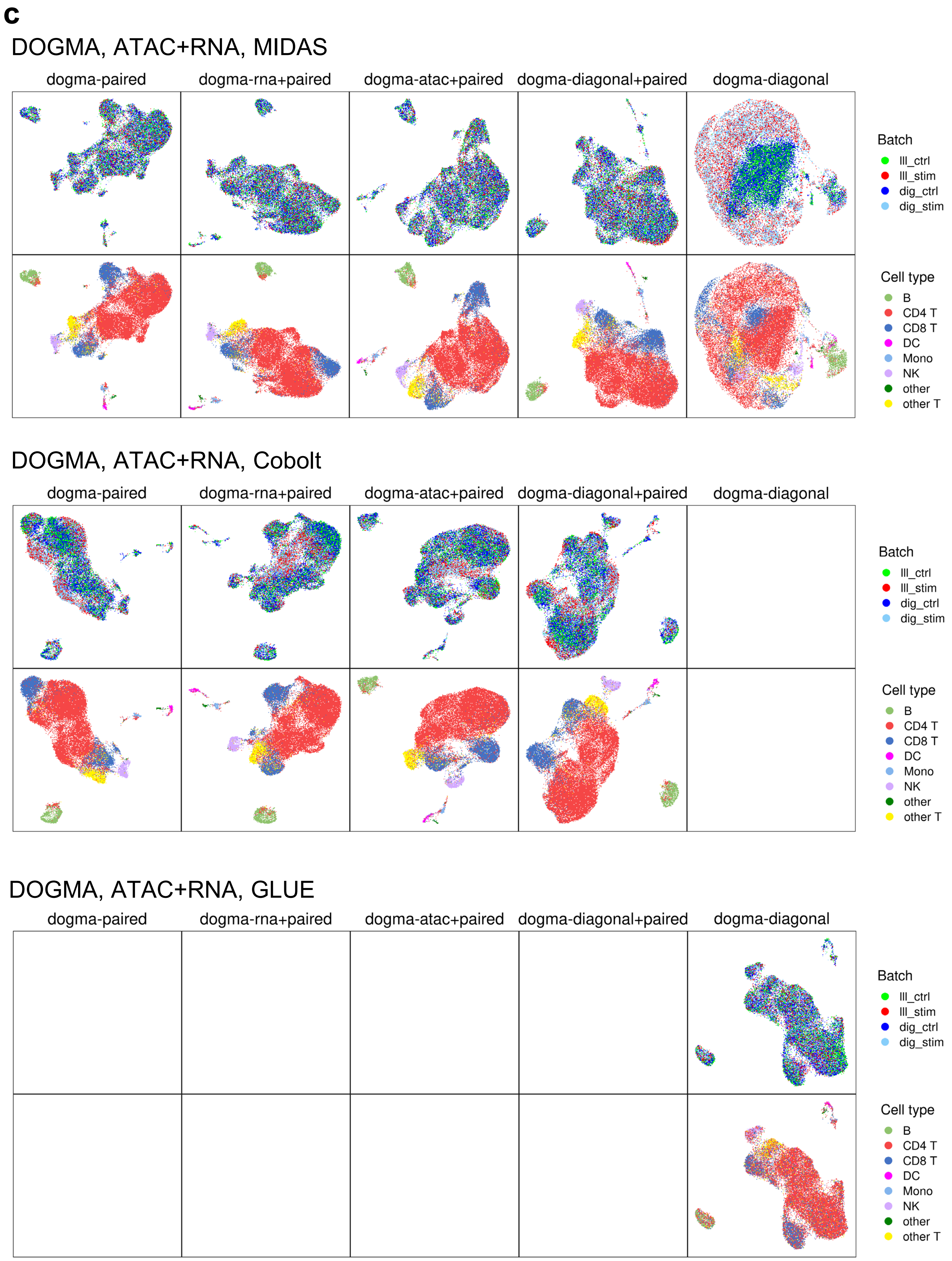
c
DOGMA, ATAC+RNA, MIDAS
dogma-paired
dogma-rna+paired
dogma-atac+paired
dogma-diagonal+paired
dogma-diagonal
Batch
lll_ctrl
lll_stim
dig_ctrl
dig_stim
Cell type
B
CD4 T
CD8 T
DC
Mono
NK
other
other T
DOGMA, ATAC+RNA, Cobolt
dogma-paired
dogma-rna+paired
dogma-atac+paired
dogma-diagonal+paired
dogma-diagonal
Batch
lll_ctrl
lll_stim
dig_ctrl
dig_stim
Cell type
B
CD4 T
CD8 T
DC
Mono
NK
other
other T
DOGMA, ATAC+RNA, GLUE
dogma-paired
dogma-rna+paired
dogma-atac+paired
dogma-diagonal+paired
dogma-diagonal
Batch
lll_ctrl
lll_stim
dig_ctrl
dig_stim
Cell type
B
CD4 T
CD8 T
DC
Mono
NK
other
other T

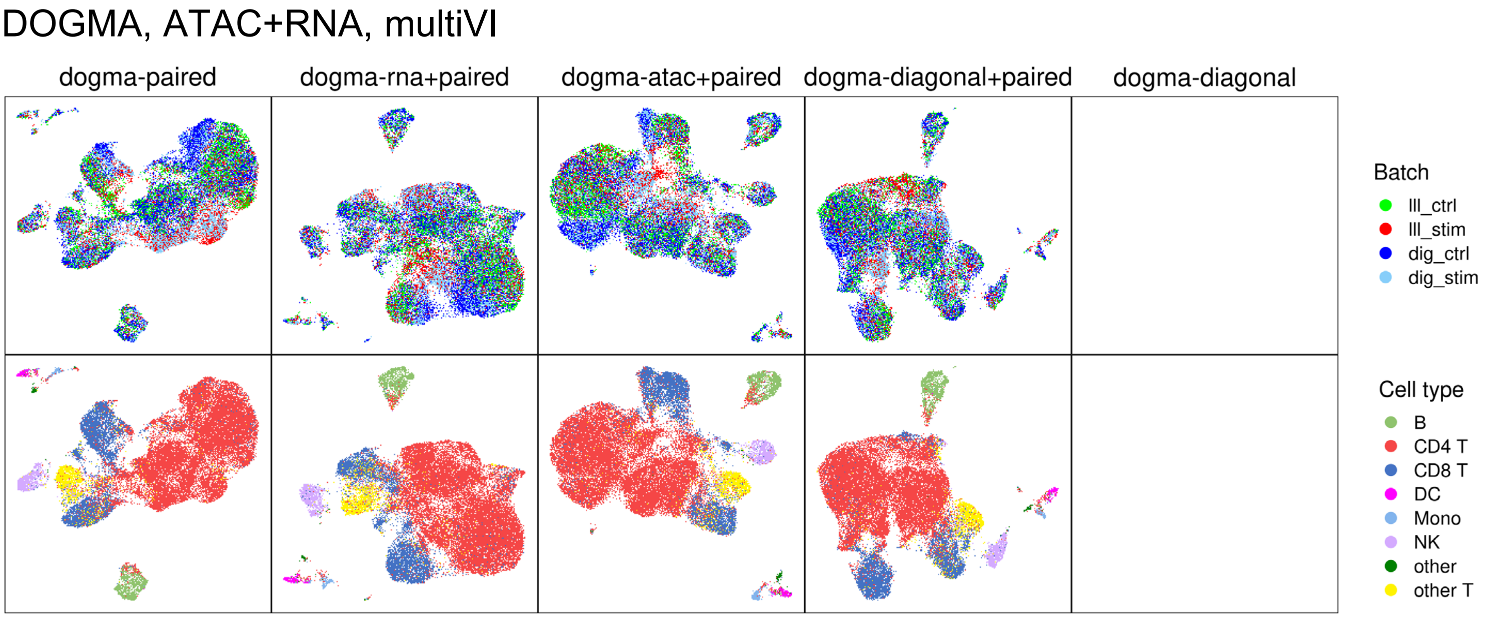
DOGMA, ATAC+RNA, multiVI
dogma-paired
dogma-rna+paired
dogma-atac+paired
dogma-diagonal+paired
dogma-diagonal
Batch
lll_ctrl
lll_stim
dig_ctrl
dig_stim
Cell type
B
CD4 T
CD8 T
DC
Mono
NK
other
other T

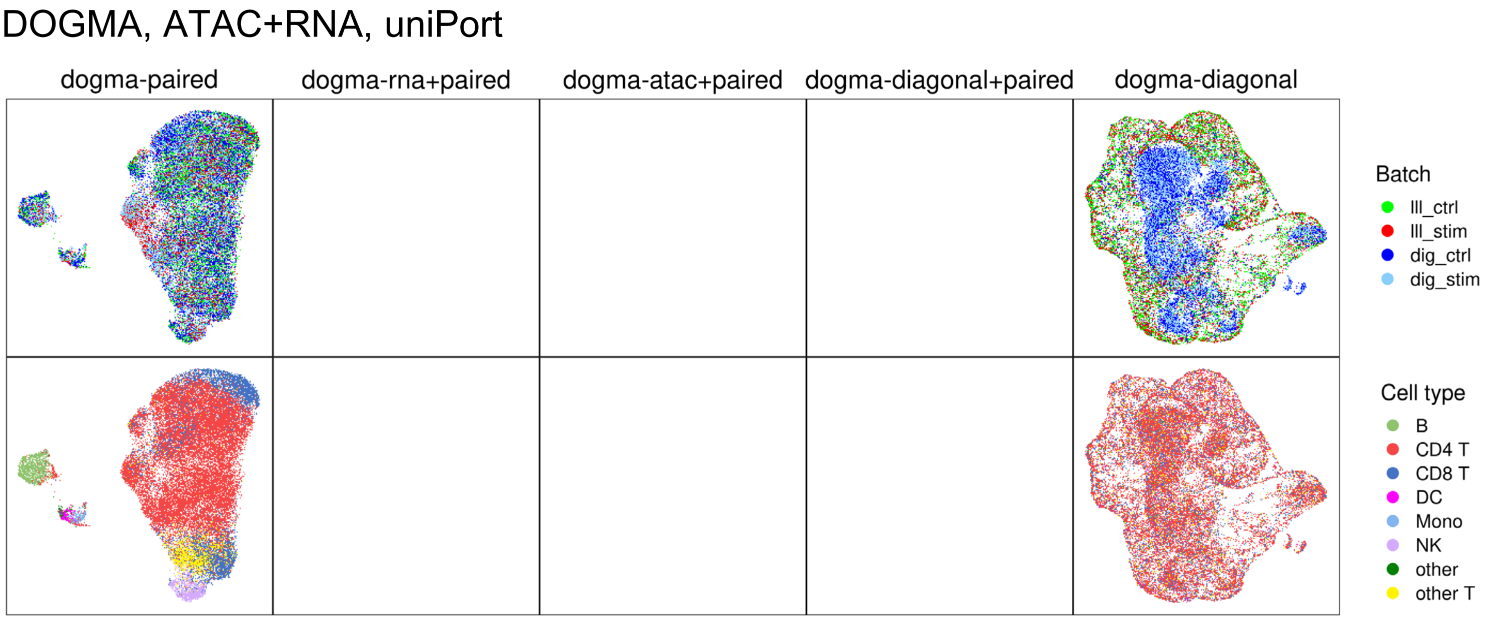
DOGMA, ATAC+RNA, uniPort
dogma-paired
dogma-rna+paired
dogma-atac+paired
dogma-diagonal+paired
dogma-diagonal
Batch
lll_ctrl
lll_stim
dig_ctrl
dig_stim
Cell type
B
CD4 T
CD8 T
DC
Mono
NK
other
other T

**d**

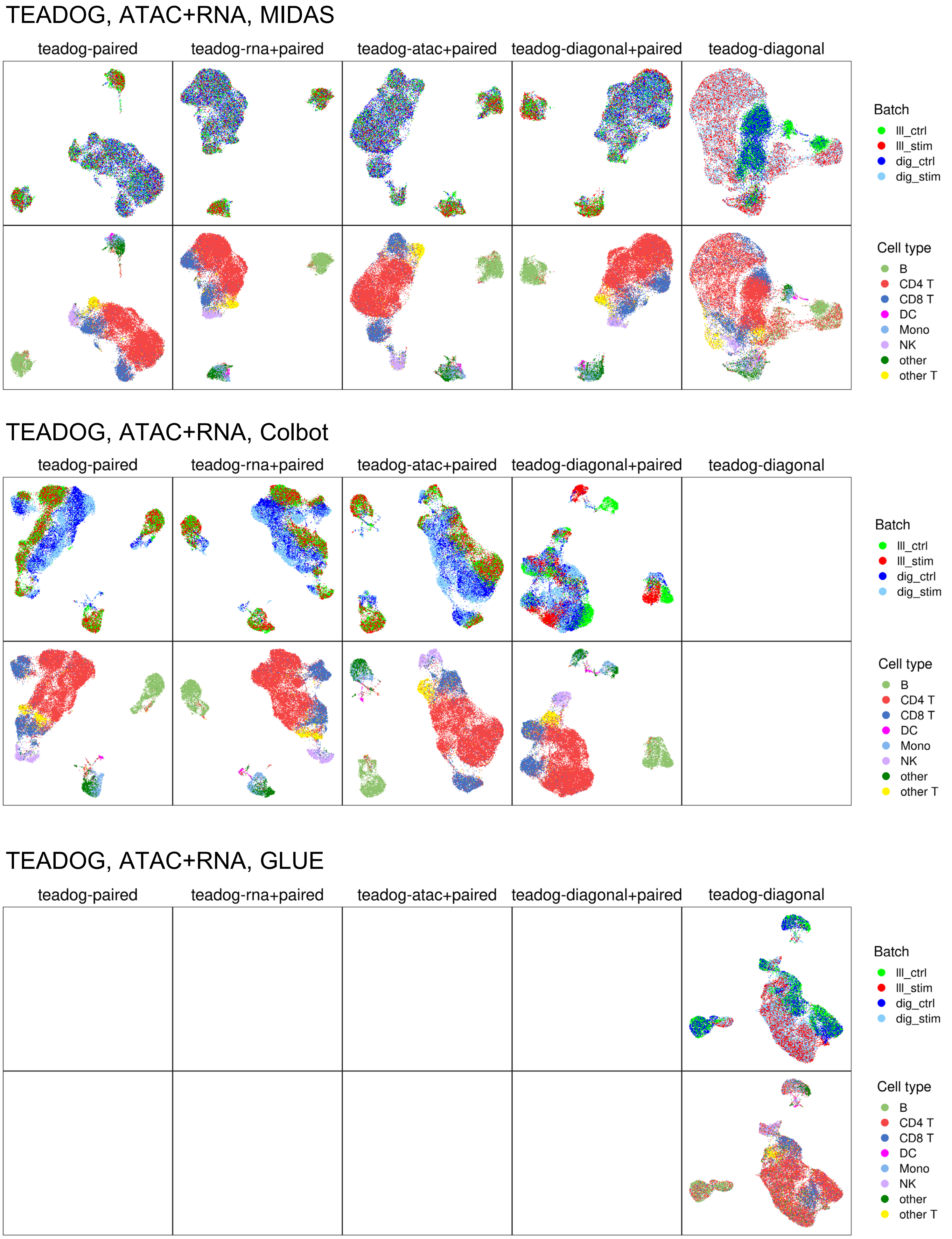

TEADOG, ATAC+RNA, MIDAS
teadog-paired
teadog-rna+paired
teadog-atac+paired
teadog-diagonal+paired
teadog-diagonal
Batch
lll_ctrl
lll_stim
dig_ctrl
dig_stim
Cell type
B
CD4 T
CD8 T
DC
Mono
NK
other
other T
TEADOG, ATAC+RNA, Colbot
teadog-paired
teadog-rna+paired
teadog-atac+paired
teadog-diagonal+paired
teadog-diagonal
Batch
lll_ctrl
lll_stim
dig_ctrl
dig_stim
Cell type
B
CD4 T
CD8 T
DC
Mono
NK
other
other T
TEADOG, ATAC+RNA, GLUE
teadog-paired
teadog-rna+paired
teadog-atac+paired
teadog-diagonal+paired
teadog-diagonal
Batch
lll_ctrl
lll_stim
dig_ctrl
dig_stim
Cell type
B
CD4 T
CD8 T
DC
Mono
NK
other
other T

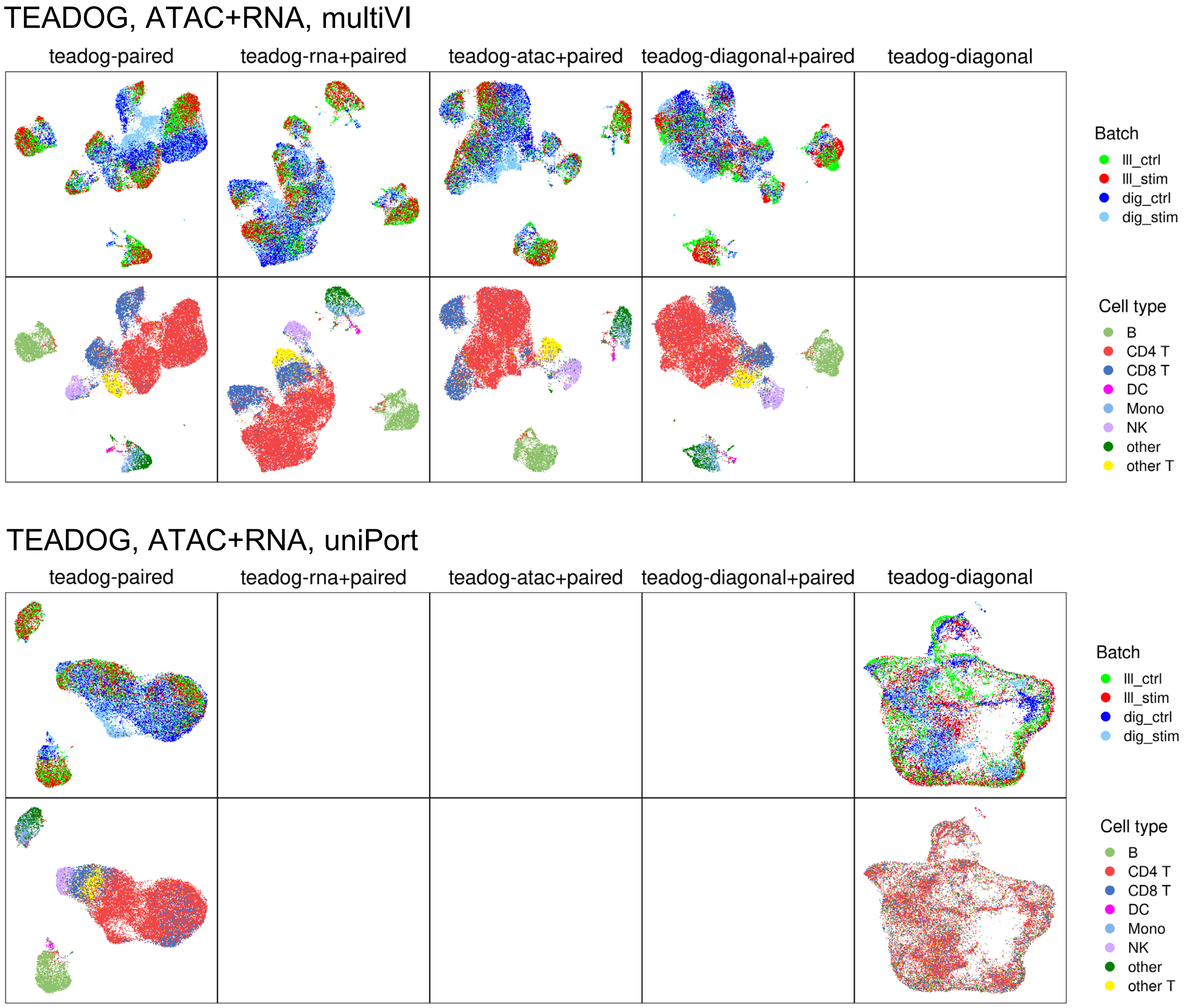

**Supplementary Figure 17. UMAP comparison of performance of MIDAS and other methods on bimodal integration tasks. a**, **b**, UMAP comparison of MIDAS, scPENN and totalVI on RNA+ADT integration tasks based on dogma (a) and teadog (b) datasets. **c**, **d**, UMAP comparison of MIDAS, Cobolt, GLUE, multiVI and uniPort on ATAC+RNA integration tasks based on dogma (c) and teadog (d) datasets.

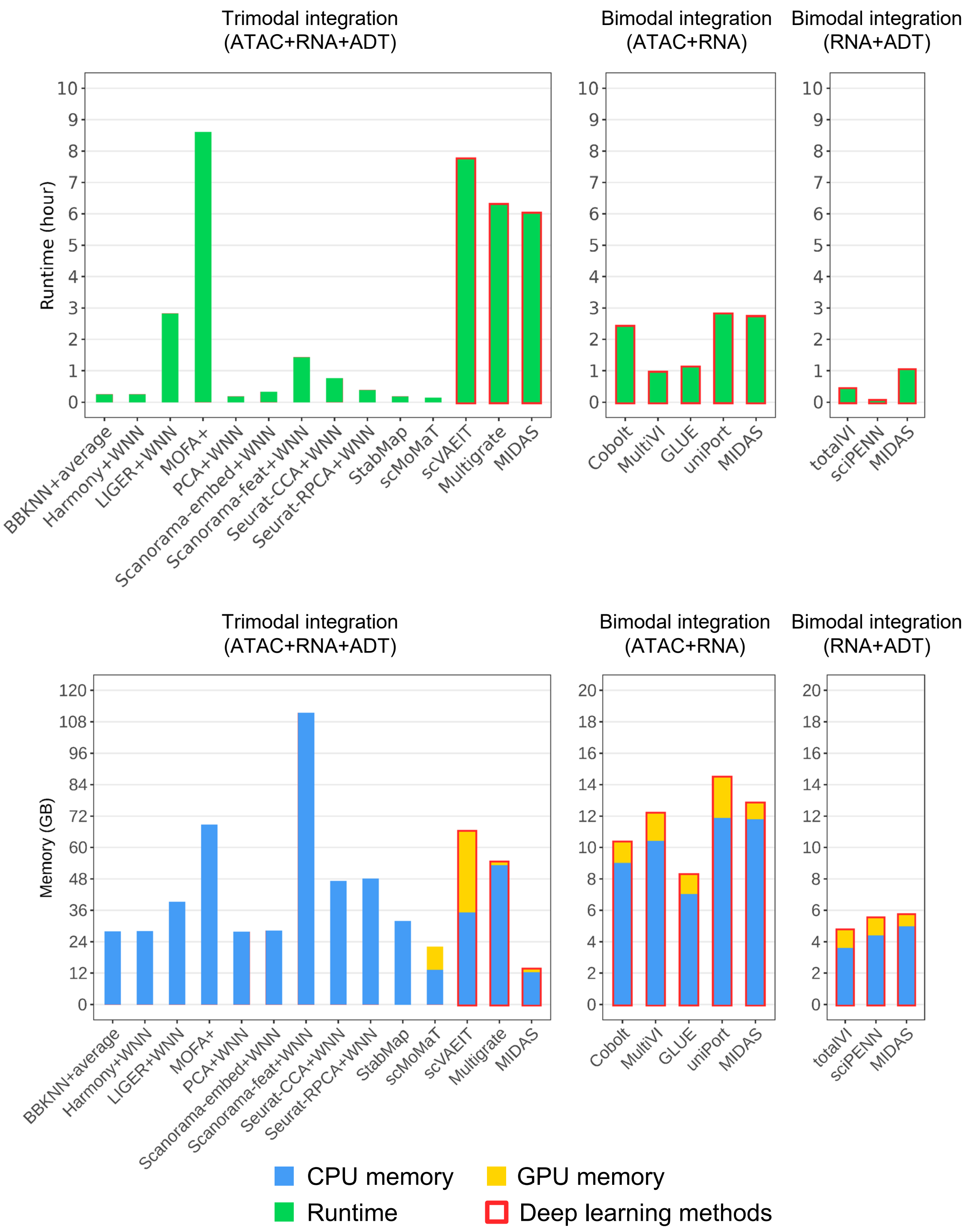

**Supplementary Figure 18. Comparison of the runtime (top) and memory (bottom) costs of different methods.**

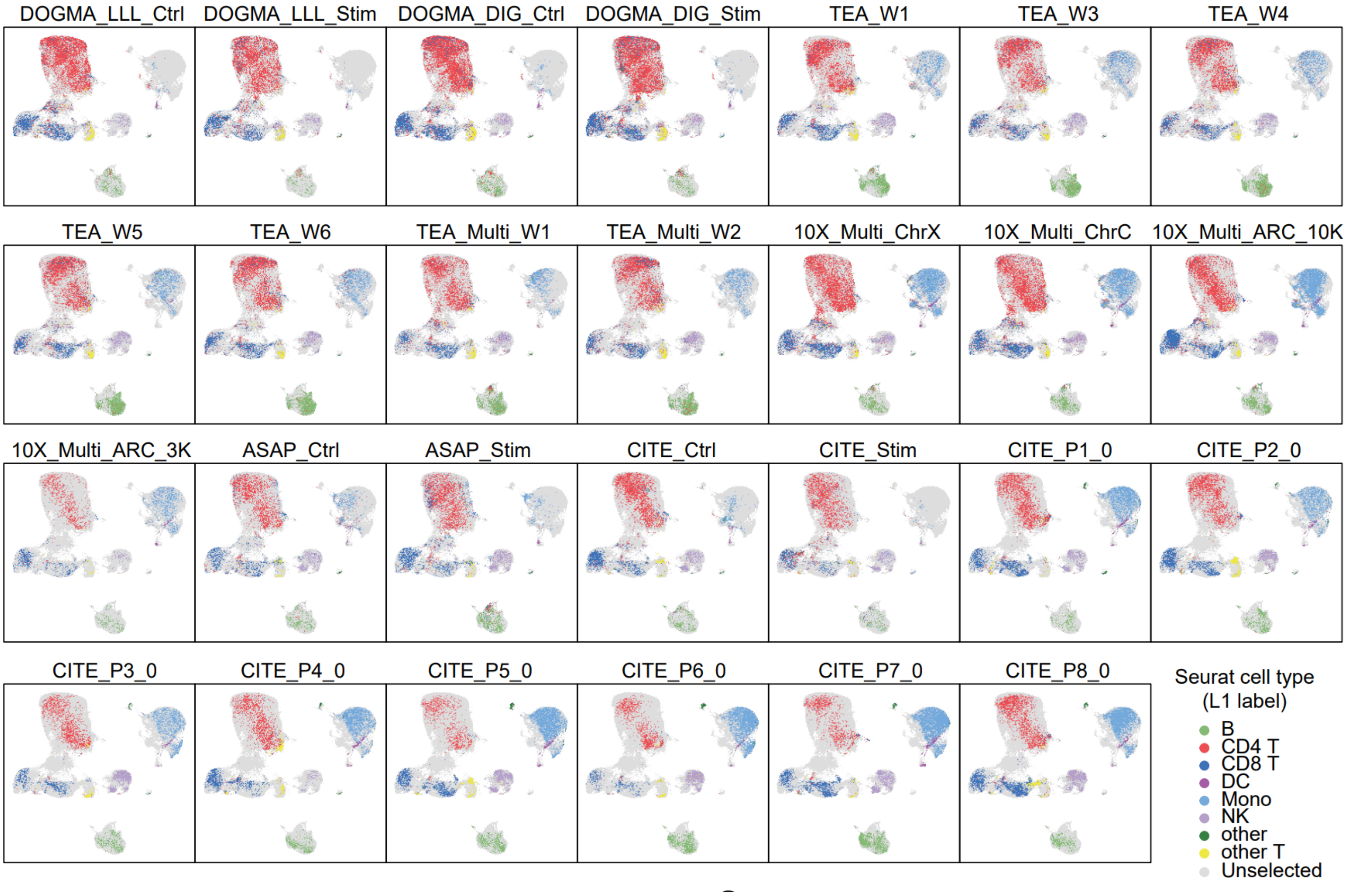

**Supplementary Figure 19. UMAP visualization of the biological states inferred by MIDAS with the PBMC mosaic dataset containing 27 batches.** Cell type labels are derived from Seurat labeling.

**a**

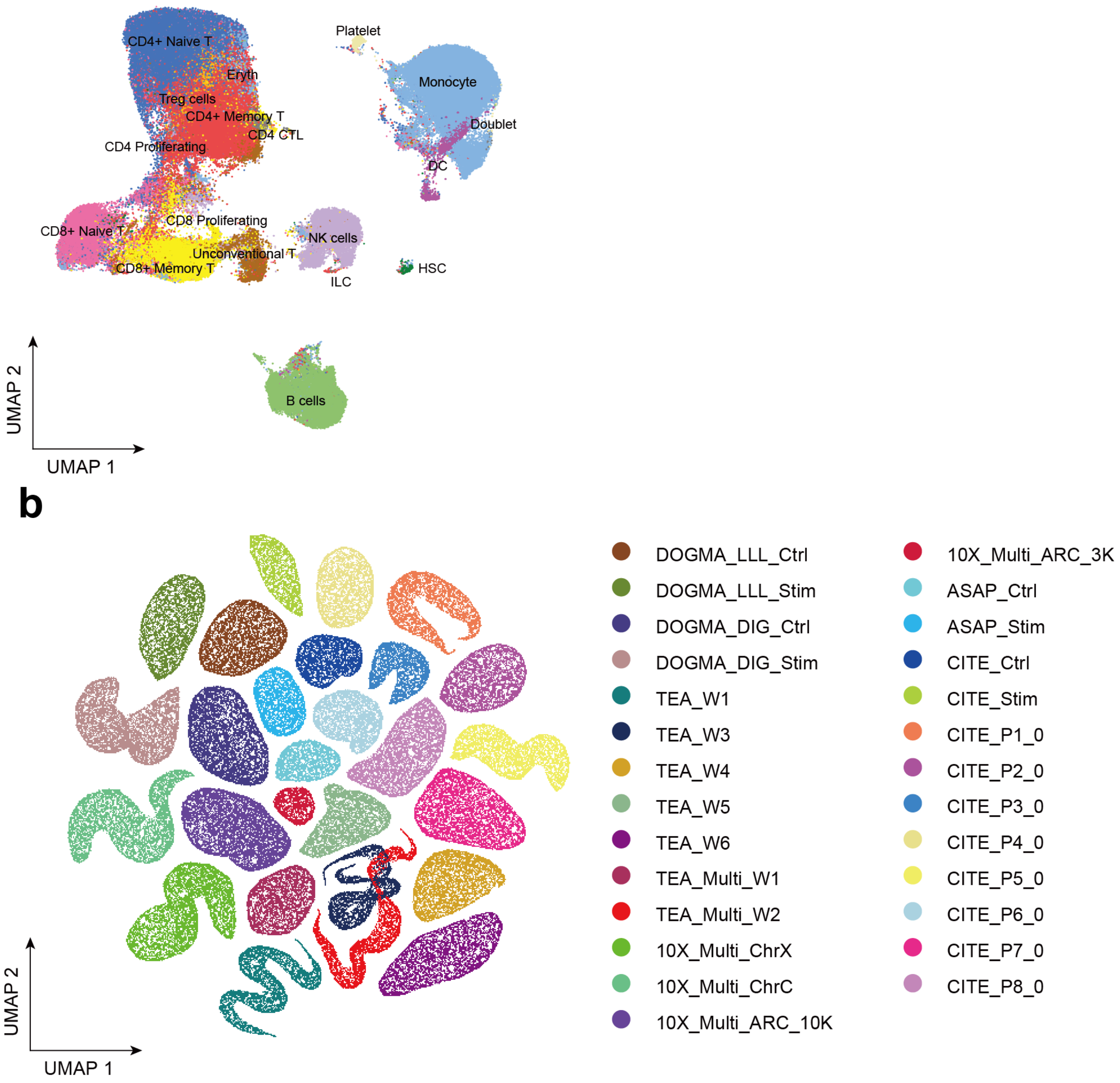

**Supplementary Figure 20. UMAP visualization (merged view) of the biological states (a) and technical noises (b) inferred by MIDAS with the PBMC mosaic dataset containing 27 batches.** Cell type colors are derived from Seurat labeling.

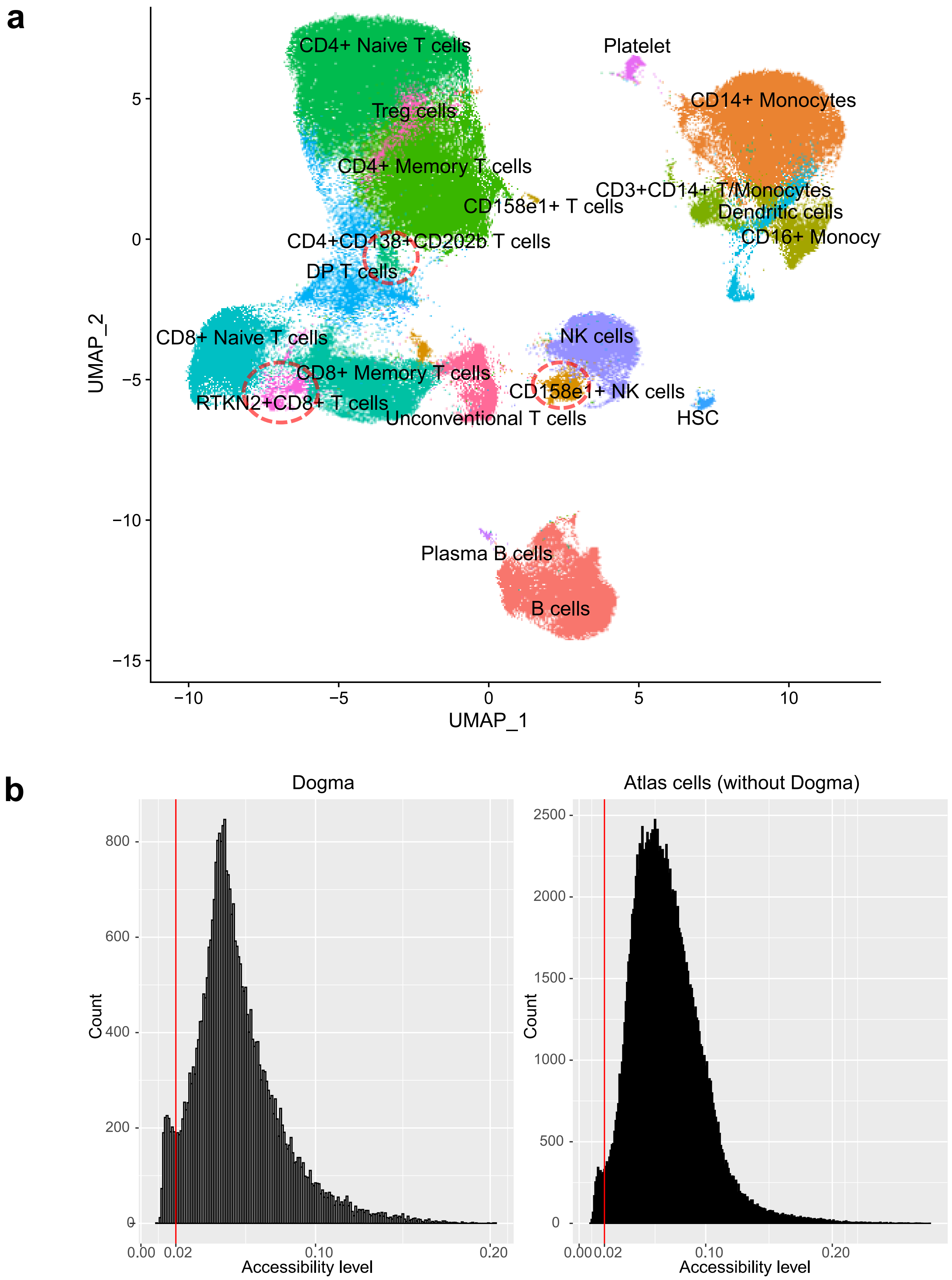

**Supplementary Figure 21. The Atlas leads to a high-resolution cell typing and LCA cells validation. a**, A more refined cell typing result of MIDAS on the atlas. **b**, The bimodal distribution of ATAC peak ration in the dogma (left) cells and atlas (right) cells without dogma. Peak ratio of a cell is defined as the ratio of peak numbers of the cell to that of all the cells

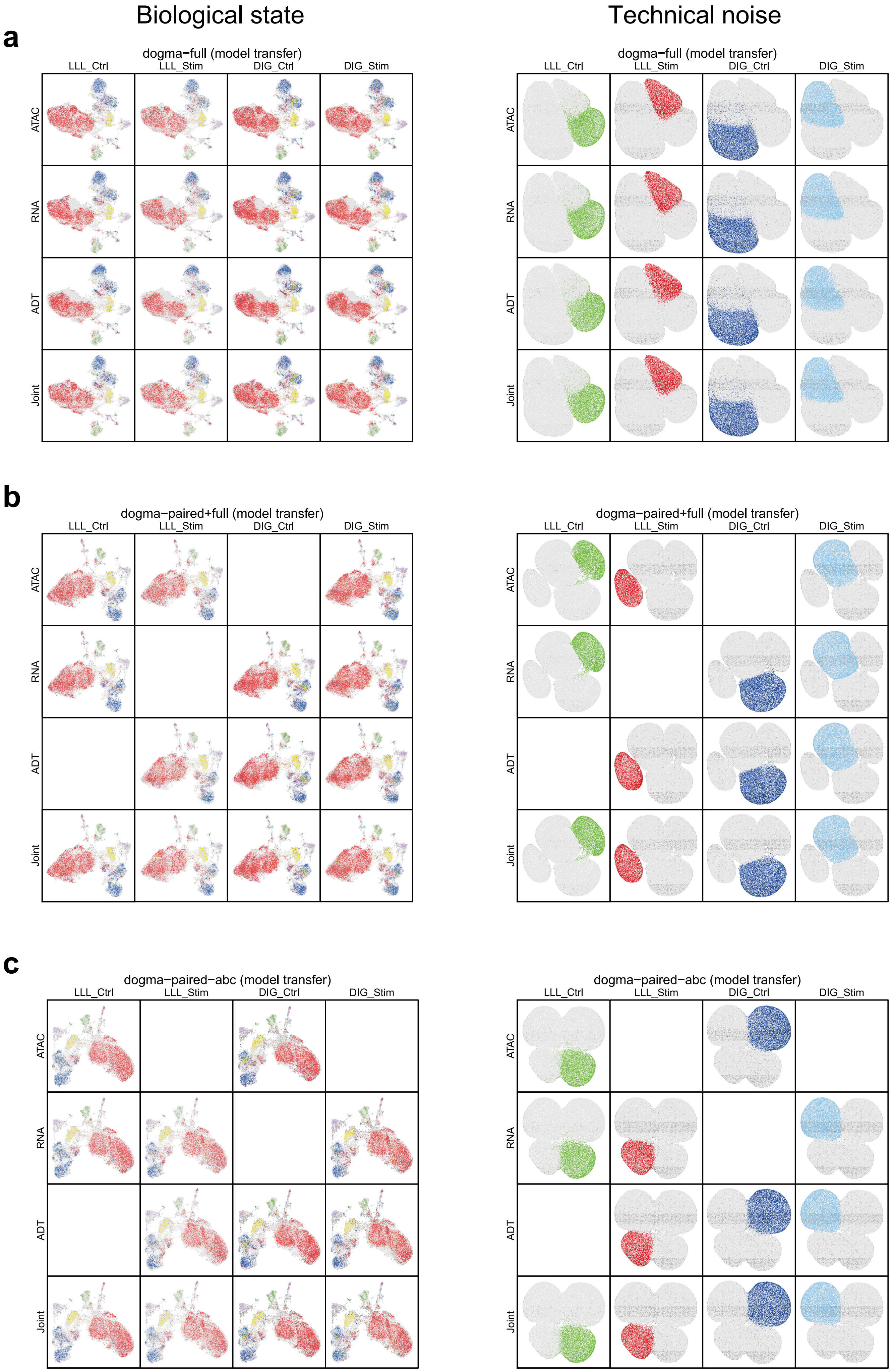
Biological state
Technical noise
a
dogma−full (model transfer)
LLL_Ctrl
LLL_Stim
DIG_Ctrl
DIG_Stim
ATAC
RNA
ADT
Joint
b
dogma−paired+full (model transfer)
c
dogma−paired−abc (model transfer)

d

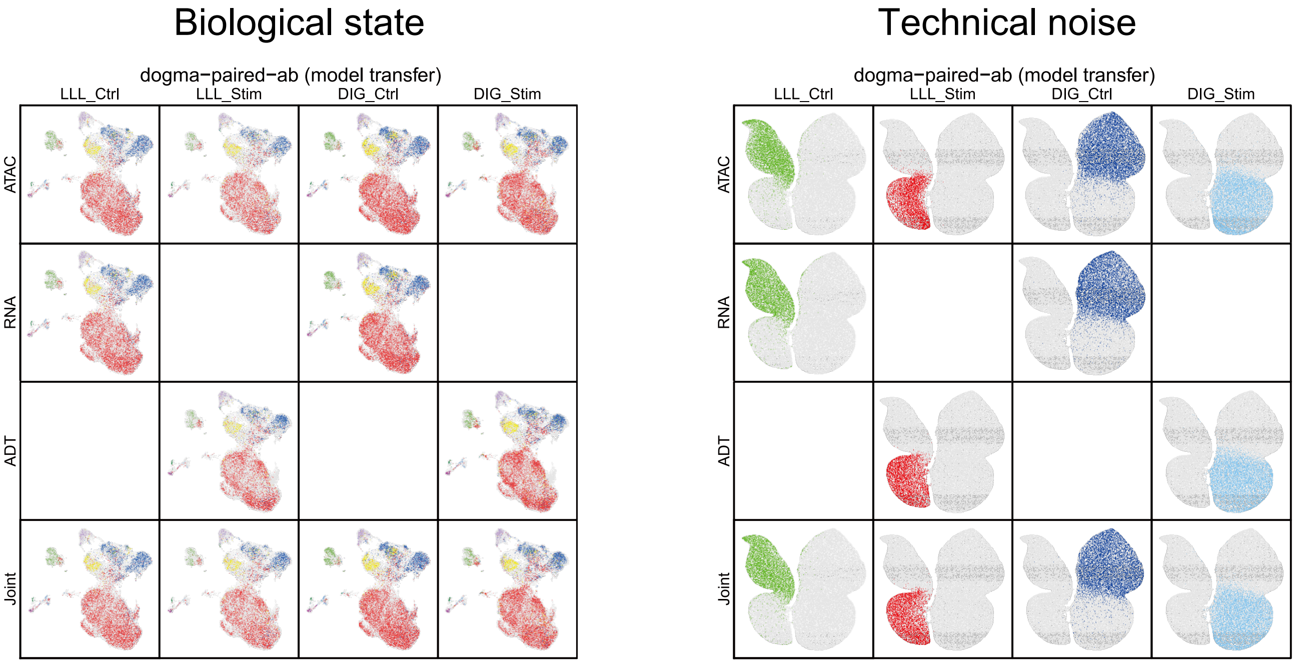
Biological state
Technical noise
dogma−paired−ab (model transfer)
LLL_Ctrl
LLL_Stim
DIG_Ctrl
DIG_Stim
ATAC
RNA
ADT
Joint

e

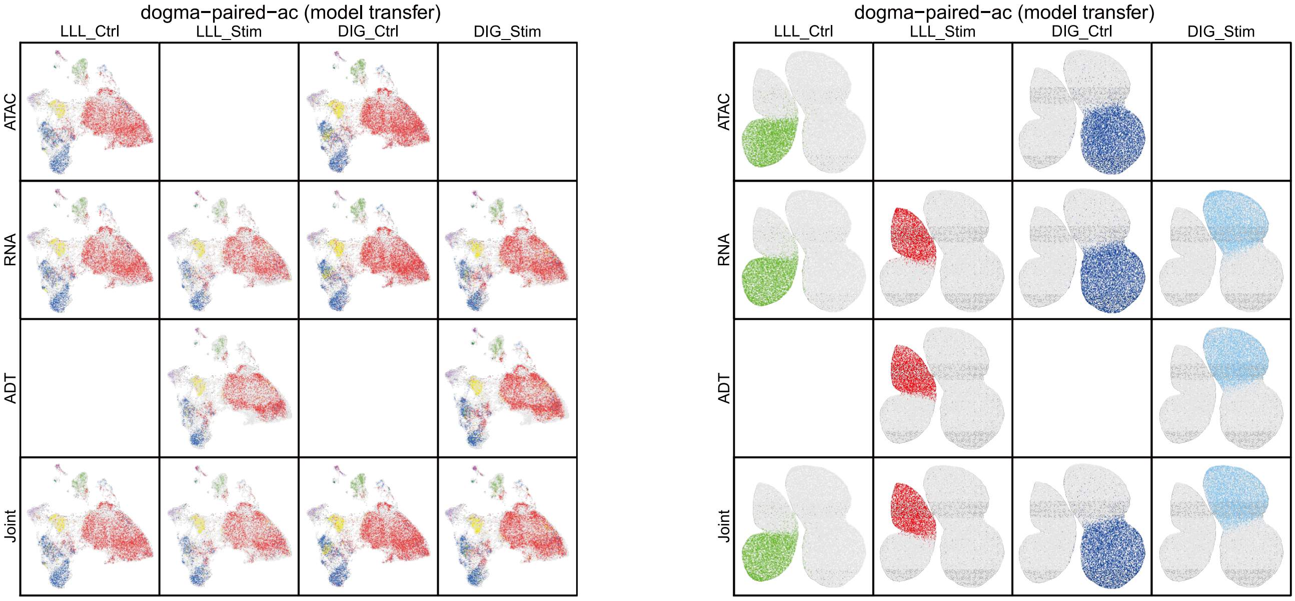
dogma−paired−ac (model transfer)
LLL_Ctrl
LLL_Stim
DIG_Ctrl
DIG_Stim
ATAC
RNA
ADT
Joint

f

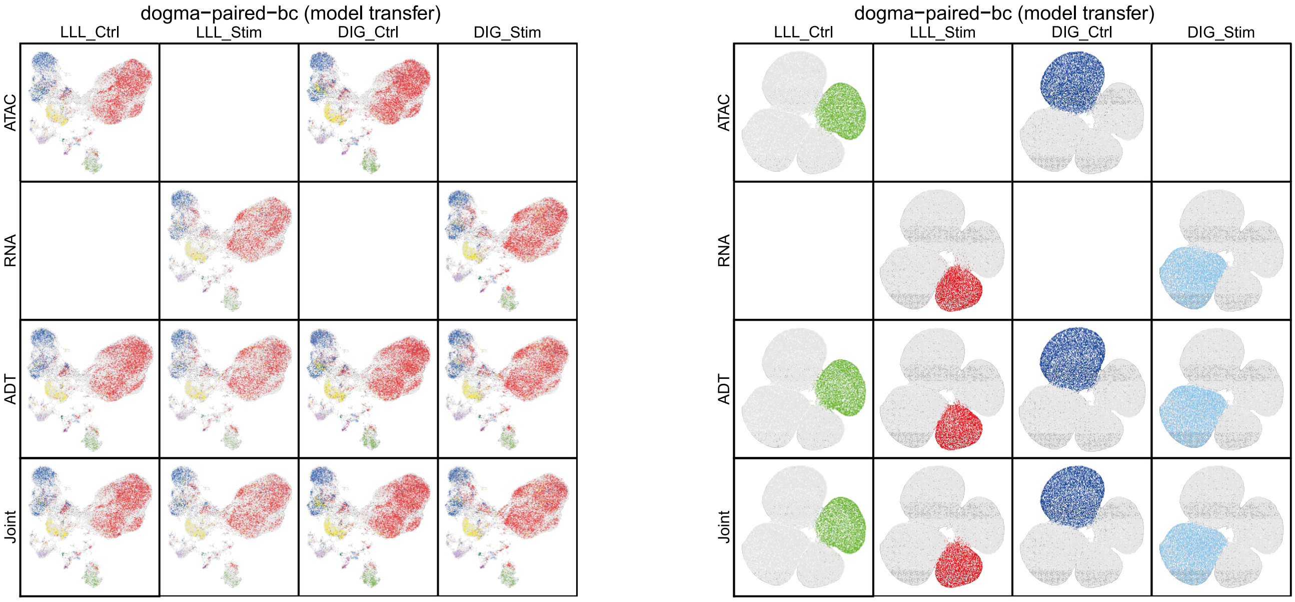
dogma−paired−bc (model transfer)
LLL_Ctrl
LLL_Stim
DIG_Ctrl
DIG_Stim
ATAC
RNA
ADT
Joint

### Biological state / Technical noise

**g**

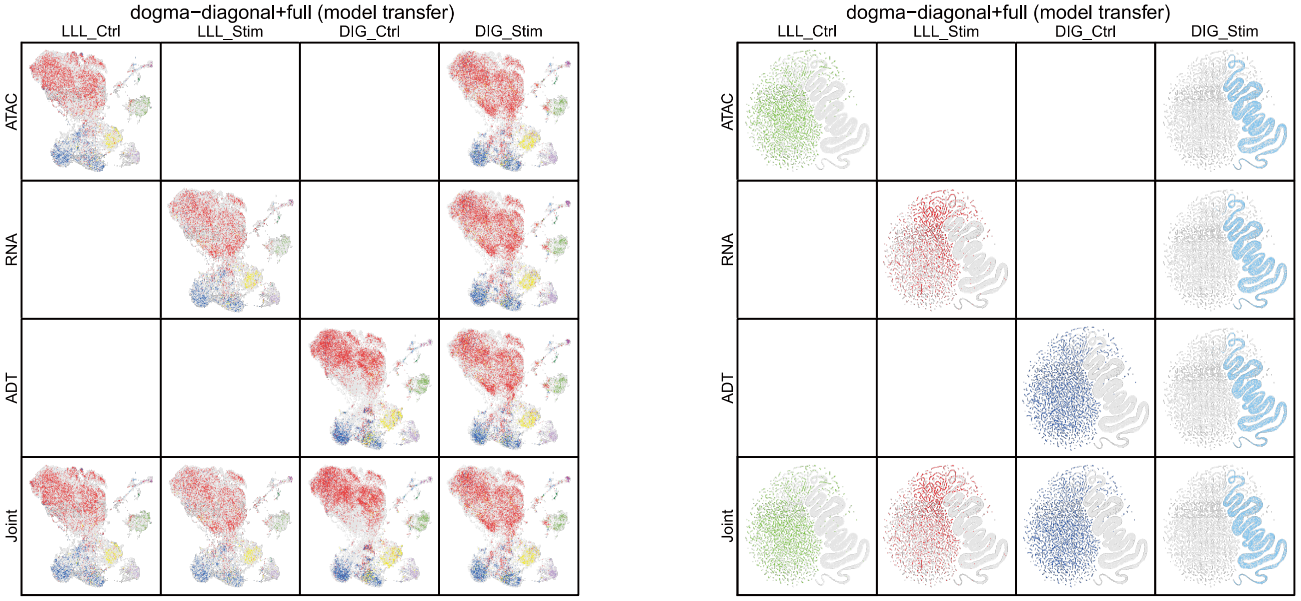

**h**

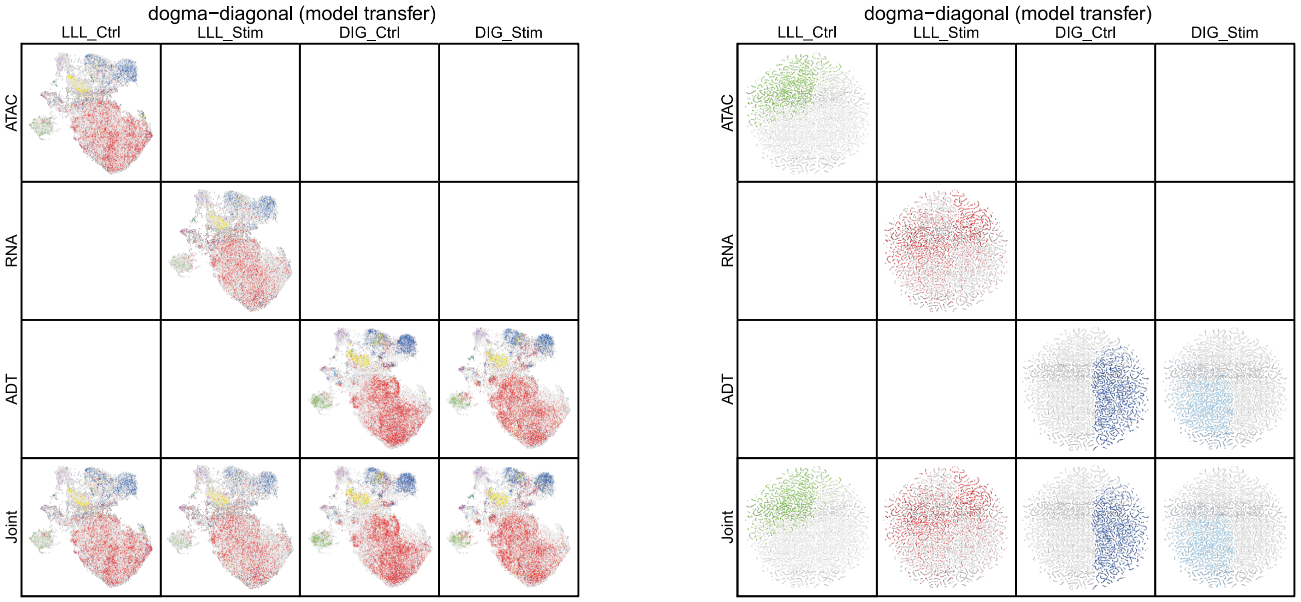

**i**

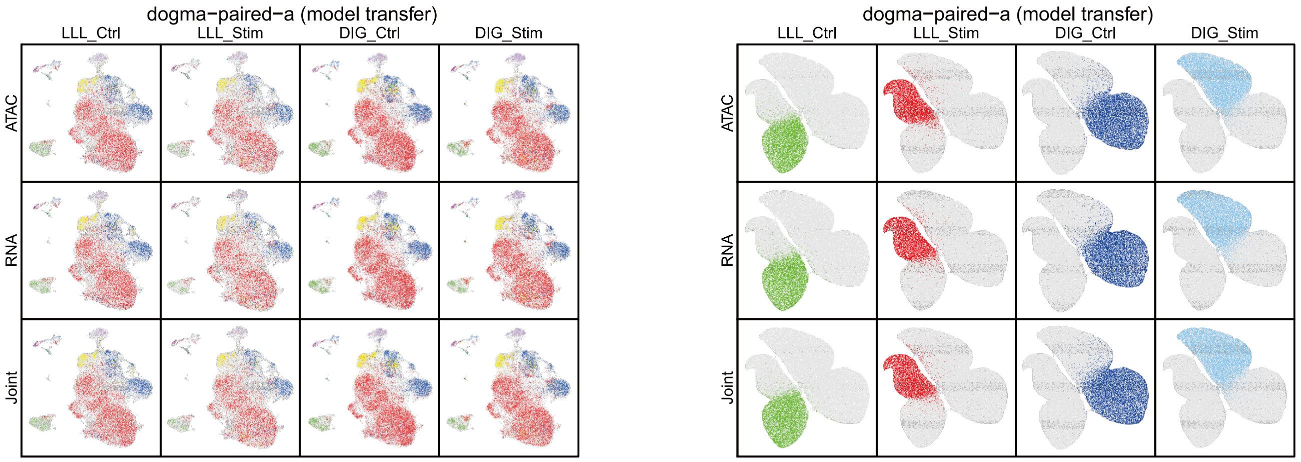

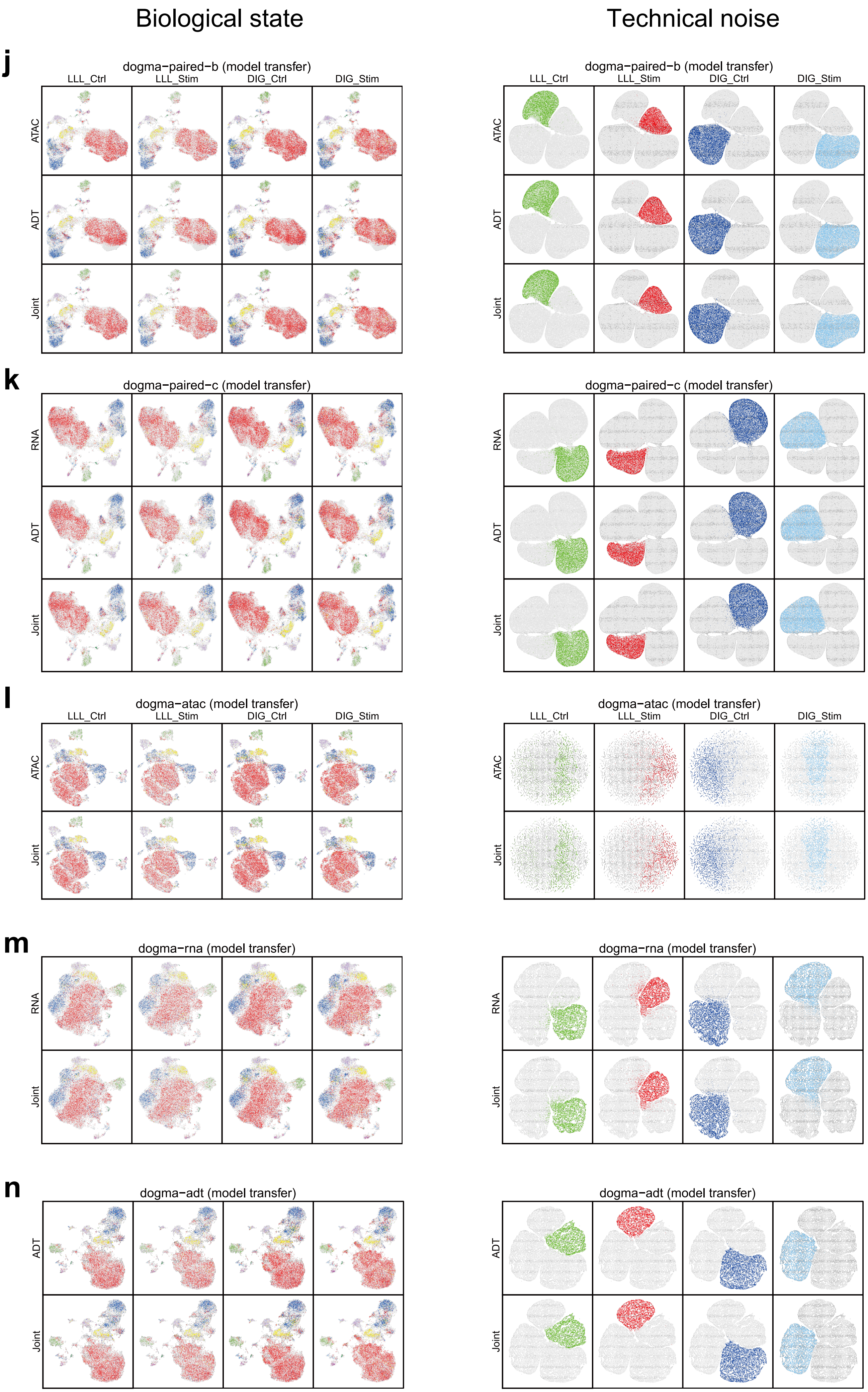
Biological state
Technical noise
j
dogma−paired−b (model transfer)
LLL_Ctrl
LLL_Stim
DIG_Ctrl
DIG_Stim
ATAC
ADT
Joint
k
dogma−paired−c (model transfer)
RNA
ADT
Joint
l
dogma−atac (model transfer)
LLL_Ctrl
LLL_Stim
DIG_Ctrl
DIG_Stim
ATAC
Joint
m
dogma−rna (model transfer)
RNA
Joint
n
dogma−adt (model transfer)
ADT
Joint

**Supplementary Figure 22. UMAP visualization of the biological states (left, colored by cell type) and technical noise (right, colored by batch) inferred by the transfer-learned MIDAS on 14 different dogma mosaic datasets (a–n).**

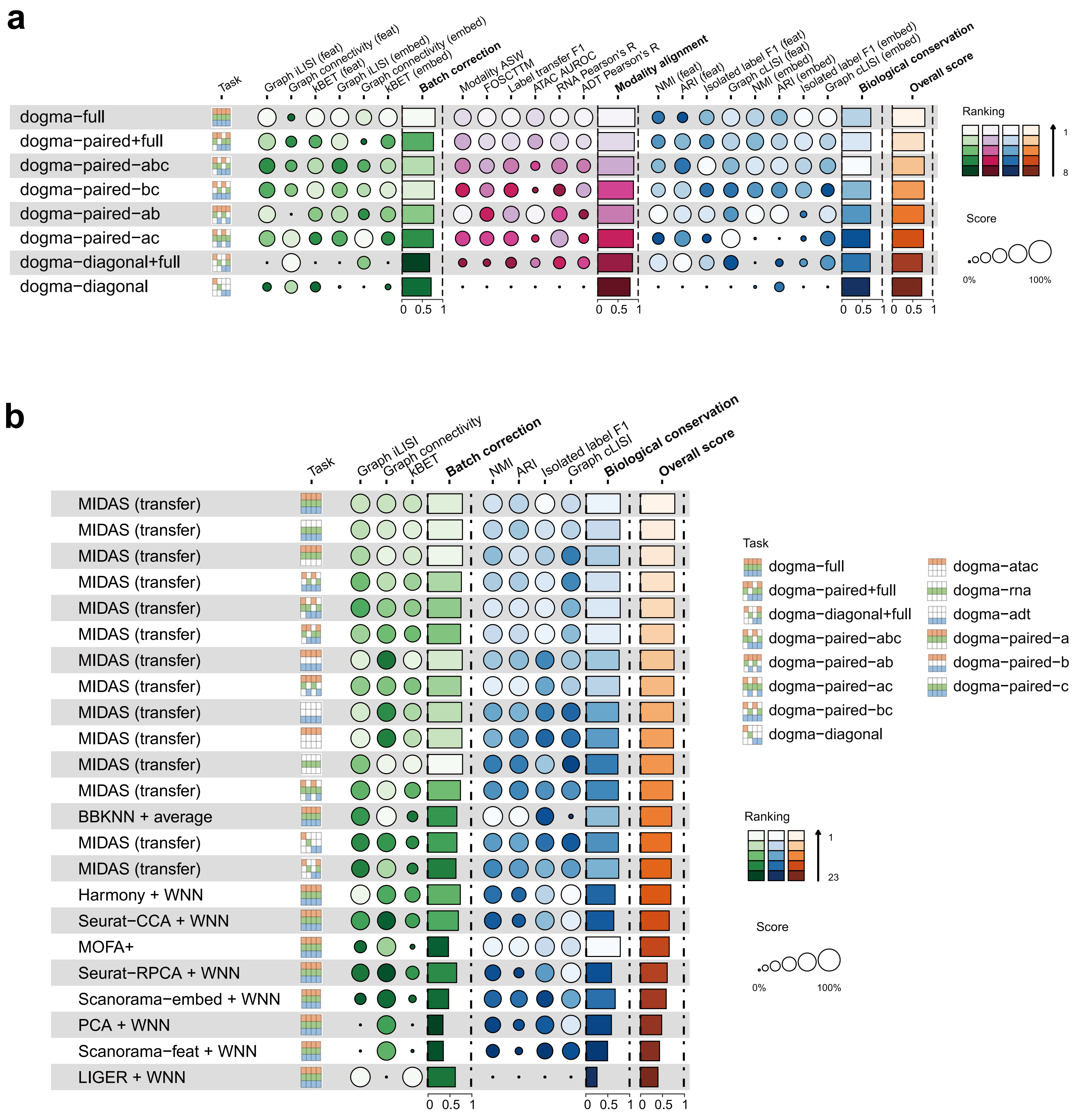

**Supplementary Figure 23. scIB and scMIB benchmarking of the performance of the transfer-learned MIDAS and other strategies. a**, Benchmarking the performance of transfer-learned MIDAS on dogma mosaic integration tasks using our proposed scMIB. **b**, scIB benchmarking the performance of transfer-learned MIDAS on dogma mosaic integration tasks against nine other strategies on the dogma-full rectangular integration task.

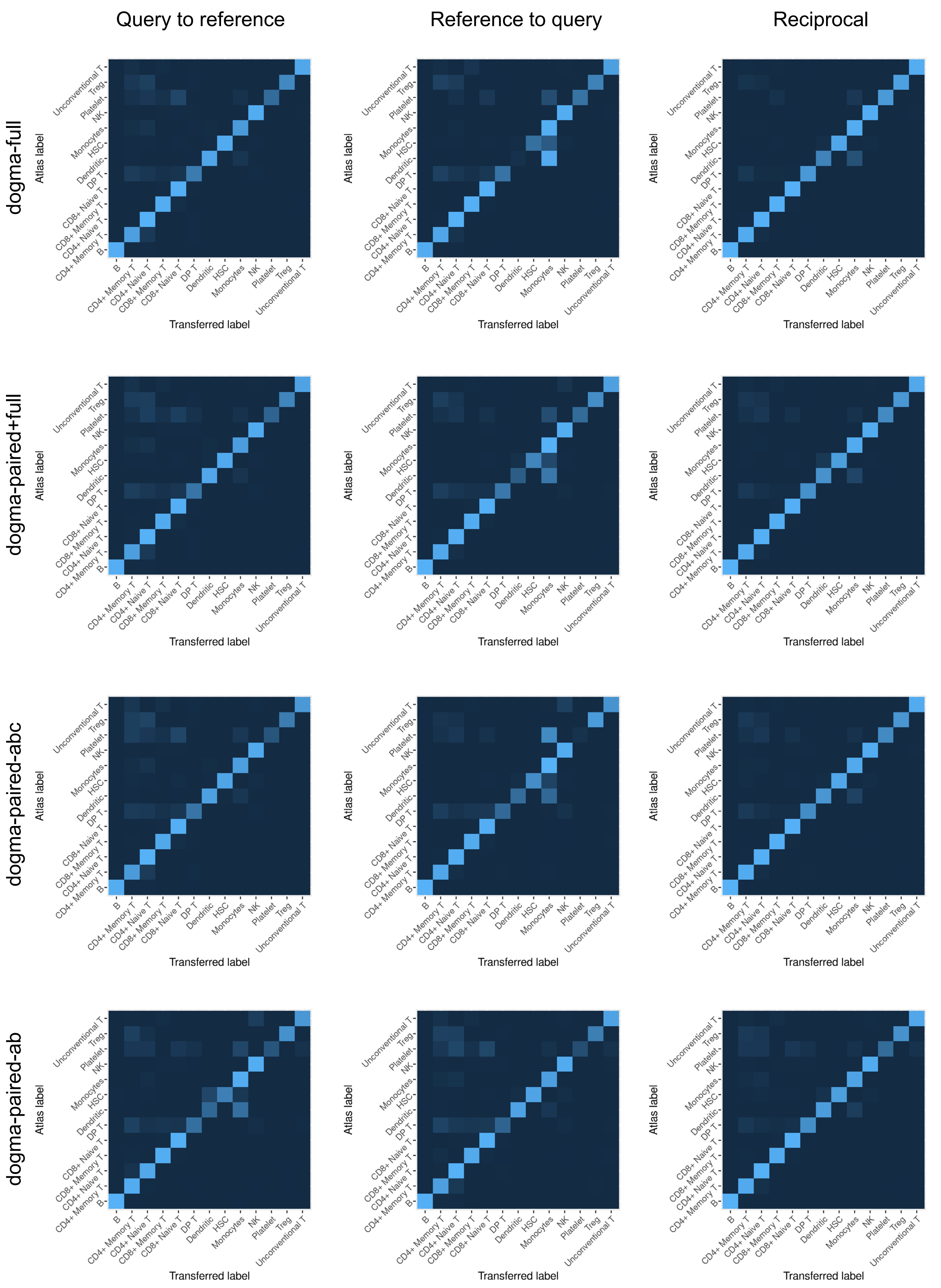

Query to reference
Reference to query
Reciprocal
dogma-full
dogma-paired+full
dogma-paired-abc
dogma-paired-ab
Atlas label
Transferred label
B
CD4+ Memory T
CD4+ Naive T
CD8+ Memory T
CD8+ Naive T
DP T
Dendritic
HSC
Monocytes
NK
Platelet
Treg
Unconventional T

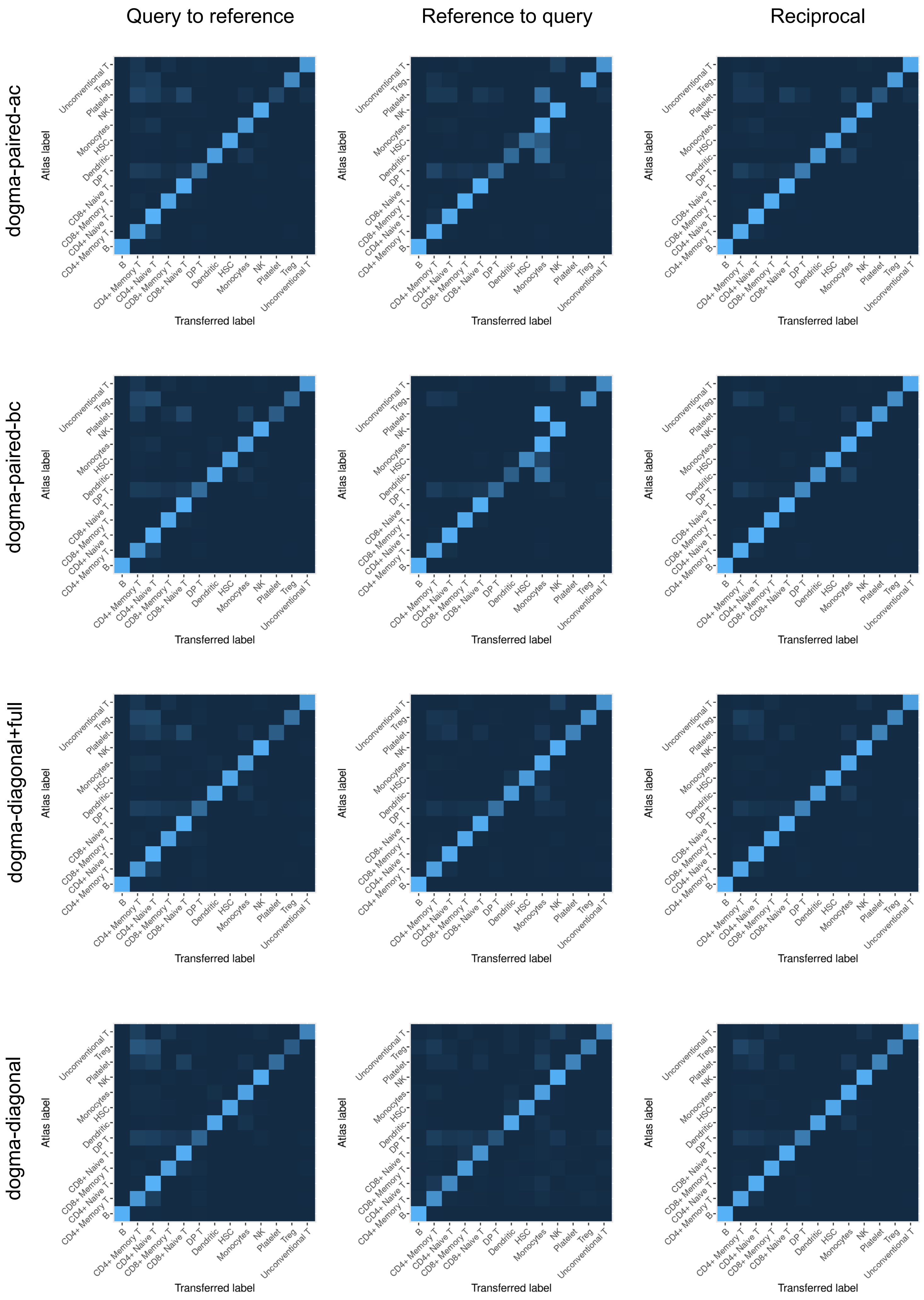
Query to reference
Reference to query
Reciprocal
dogma-paired-ac
dogma-paired-bc
dogma-diagonal+full
dogma-diagonal
Atlas label
Transferred label
B
CD4+ Memory T
CD4+ Naive T
CD8+ Memory T
CD8+ Naive T
DP T
Dendritic
HSC
Monocytes
NK
Platelet
Treg
Unconventional T

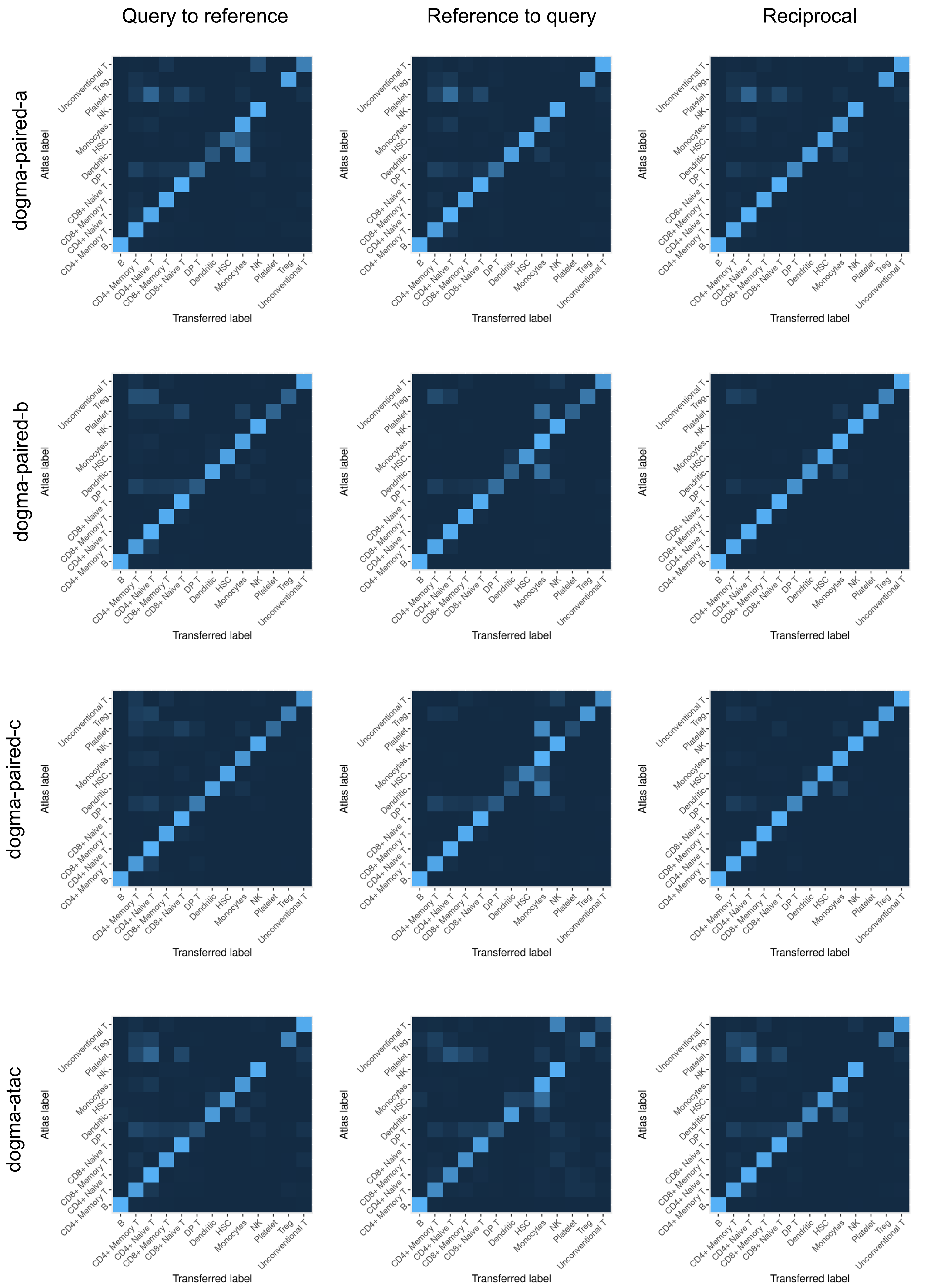
Query to reference
Reference to query
Reciprocal
dogma-paired-a
dogma-paired-b
dogma-paired-c
dogma-atac
Atlas label
Transferred label
B
CD4+ Memory T
CD4+ Naive T
CD8+ Memory T
CD8+ Naive T
DP T
Dendritic
HSC
Monocytes
NK
Platelet
Treg
Unconventional T

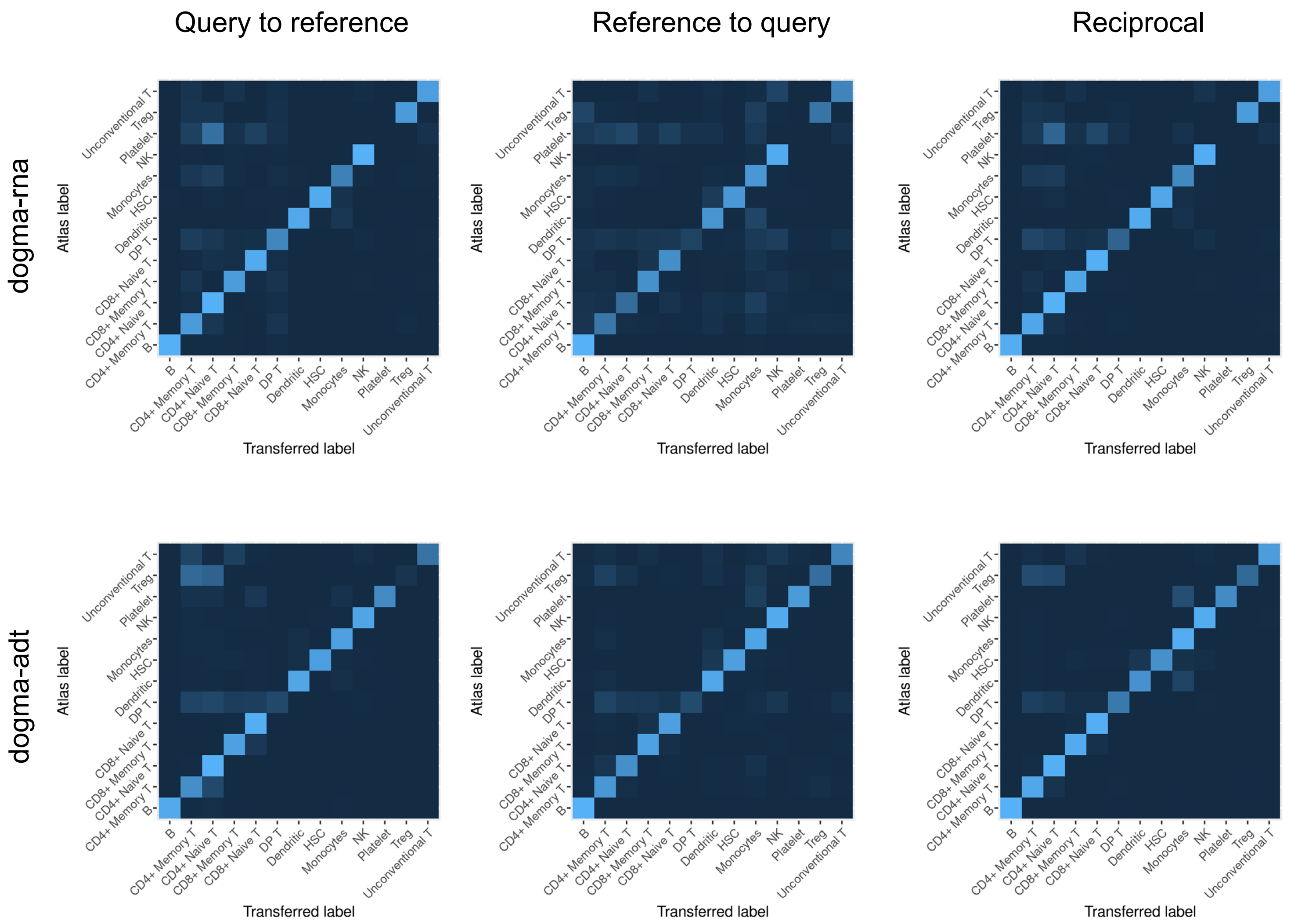

**Supplementary Figure 24. Comparison of confusion plots corresponding to the label transfer micro F1-scores of different mapping strategies on different dogma query dataset.** In each plot, rows correspond to PBMC atlas labels and columns correspond to transferred labels. The horizontal plots show 14 tasks while the vertical plots show query-to-reference mapping, reference-to-query mapping and reciprocal reference mapping methods.

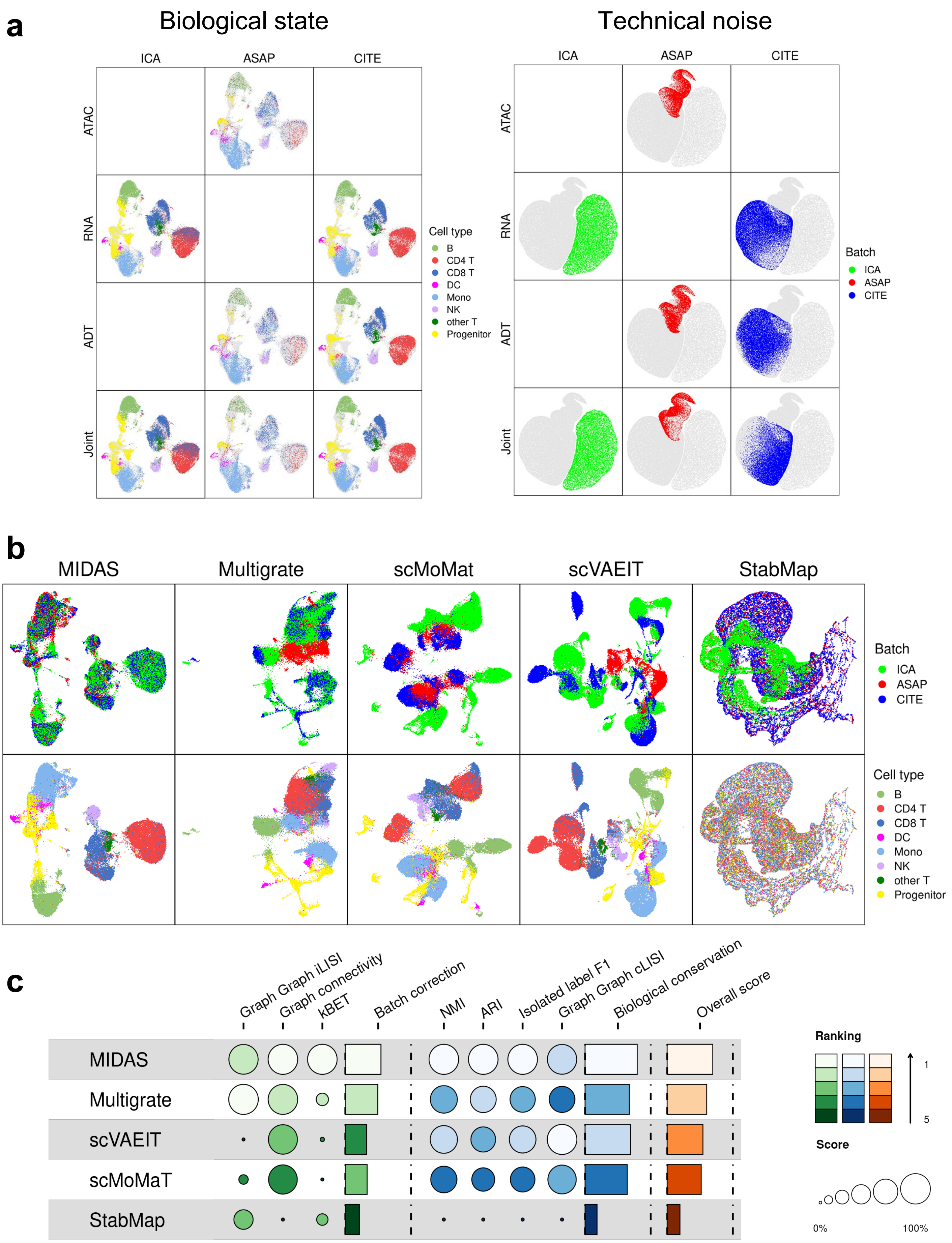

**Supplementary Figure 25. Qualitative and quantitative evaluation of MIDAS performance on BMMC mosaic dataset. a,** UMAP visualization of the biological states (left panel) and technical noises (right panel) inferred by MIDAS. **b,** UMAP comparison of embeddings on BMMC mosaic integration. Cells in the top row are colored by batches, and cells in the bottom row are colored by cell types. **c,** scIB benchmarking the performance of MIDAS against four methods.

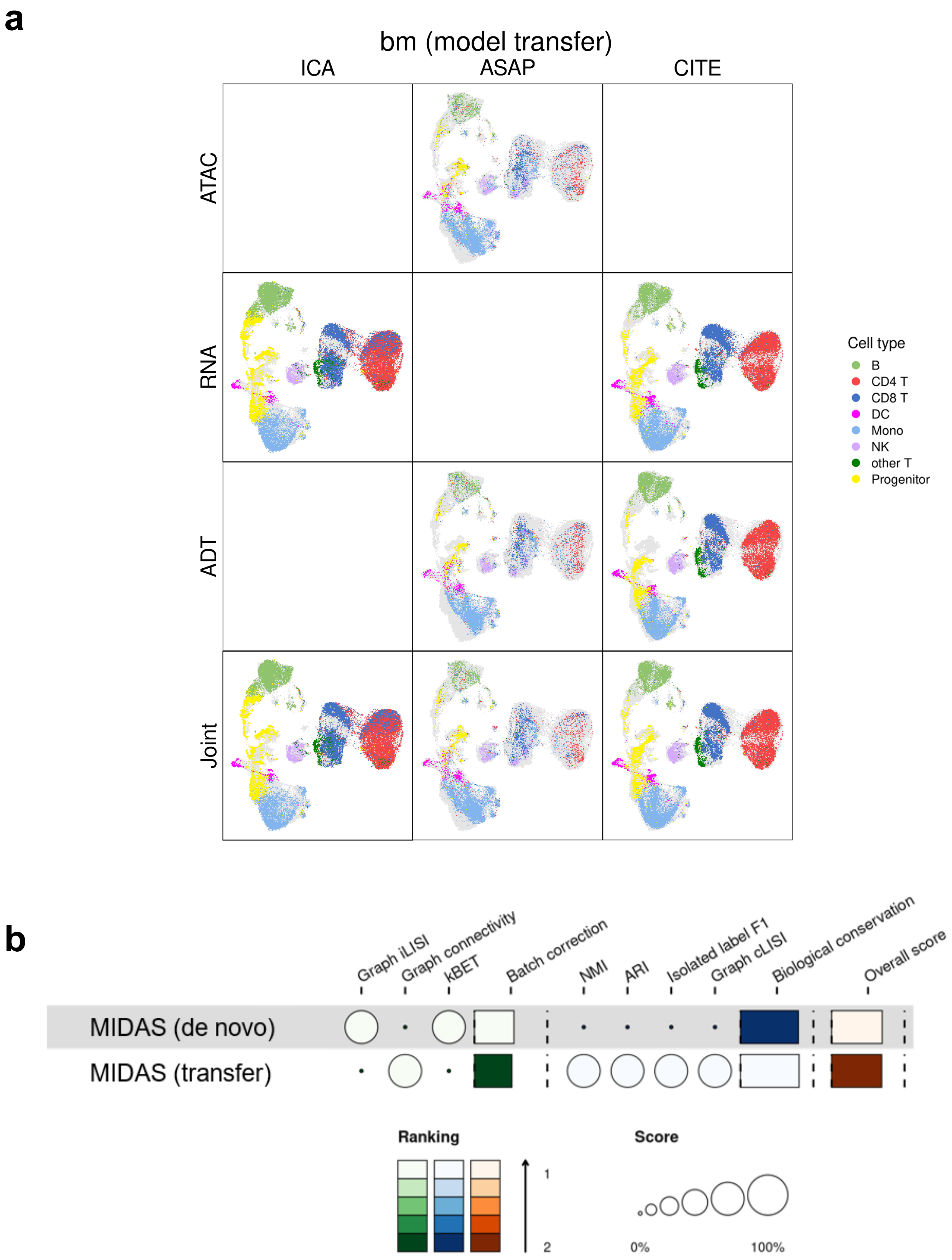

**Supplementary Figure 26. Performance of MIDAS on model transfer in BMMC mosaic dataset. a**, UMAP visualization of the biological states inferred by the transfer-learned MIDAS on bm dataset. **b**, scIB benchmarking the performance of *de novo* and model transfer integration.

Supplementary Table 1. Public datasets used in this study.

| Dataset | Batch | Protocal | Accession ID/URL | Filtered cell number | RNA | ADT | ATAC |
|---|---|---|---|---|---|---|---|
| DOGMA dataset | LLL Ctrl | DOGMA-seq | GSE166188 (GSM5065524, GSM5065525, GSM5065526) | 7,361 | √ | √ | √ |
| | LLL Stim | DOGMA-seq | GSE166188 (GSM5065527, GSM5065528, GSM5065529) | 5,897 | √ | √ | √ |
| | DIG Ctrl | DOGMA-seq | GSE166188 (GSM5065530, GSM5065531, GSM5065532) | 10,190 | √ | √ | √ |
| | DIG Stim | DOGMA-seq | GSE166188 (GSM5065533, GSM5065534, GSM5065535) | 9,527 | √ | √ | √ |
| TEA dataset | W1 | TEA-seq | GSE158013 (GSM4949911) | 7,325 | √ | √ | √ |
| | W3 | TEA-seq | GSE158013 (GSM5123951) | 6,587 | √ | √ | √ |
| | W4 | TEA-seq | GSE158013 (GSM5123952) | 6,897 | √ | √ | √ |
| | W5 | TEA-seq | GSE158013 (GSM5123953) | 6,910 | √ | √ | √ |
| | W6 | TEA-seq | GSE158013 (GSM5123954) | 7,137 | √ | √ | √ |
| TEA Multiome dataset | W1 | 10x Chromium Single Cell Multiome ATAC + Gene Expression | GSE158013 (GSM5123949) | 6,096 | √ | | √ |
| | W2 | 10x Chromium Single Cell Multiome ATAC + Gene Expression | GSE158013 (GSM5123950) | 7,284 | √ | | √ |
| 10X Multiome dataset | ChrX | 10x Chromium Single Cell Multiome ATAC + Gene Expression | https://www.10xgenomics.com/resources/datasets/10-k-human-pbm-cs-multiome-v-1-0-chromium-x-1-standard-2-0-0 | 9,868 | √ | | √ |
| | ChrC | 10x Chromium Single Cell Multiome ATAC + Gene Expression | https://www.10xgenomics.com/resources/datasets/10-k-human-pbm-cs-multiome-v-1-0-chromium-controller-1-standard-2-0-0 | 9,582 | √ | | √ |
| | ARC2 10K | 10x Chromium Single Cell Multiome ATAC + Gene Expression | https://www.10xgenomics.com/resources/datasets/pbmc-from-a-healthy-donor-no-cell-sorting-10-k-1-standard-2-0-0 | 11,116 | √ | | √ |
| | ARC2 3K | 10x Chromium Single Cell Multiome ATAC + Gene Expression | https://www.10xgenomics.com/resources/datasets/pbmc-from-a-healthy-donor-no-cell-sorting-3-k-1-standard-2-0-0 | 2,566 | √ | | √ |
| ASAP dataset | Ctrl | ASAP-seq | GSE156473 (GSM4732109, GSM4732110) | 4,255 | | √ | √ |
| | Stim | ASAP-seq | GSE156473 (GSM4732111, GSM4732112) | 5,241 | | √ | √ |
| ASAP CITE dataset | Ctrl | CITE-seq | GSE156473 (GSM4732113, GSM4732114) | 5,086 | √ | √ | |
| | Stim | CITE-seq | GSE156473 (GSM4732115, GSM4732116) | 3,629 | √ | √ | |

| | | | | | | | |
|---|---|---|---|---|---|---|---|
| WNN CITE dataset | P1_0 | CITE-seq | https://atlas.fredhutch.org/nygc/multimodal-pbmc/ | 6,378 | √ | √ | |
| | P2_0 | CITE-seq | https://atlas.fredhutch.org/nygc/multimodal-pbmc/ | 5,899 | √ | √ | |
| | P3_0 | CITE-seq | https://atlas.fredhutch.org/nygc/multimodal-pbmc/ | 4,628 | √ | √ | |
| | P4_0 | CITE-seq | https://atlas.fredhutch.org/nygc/multimodal-pbmc/ | 5,285 | √ | √ | |
| | P5_0 | CITE-seq | https://atlas.fredhutch.org/nygc/multimodal-pbmc/ | 6,952 | √ | √ | |
| | P6_0 | CITE-seq | https://atlas.fredhutch.org/nygc/multimodal-pbmc/ | 6,060 | √ | √ | |
| | P7_0 | CITE-seq | https://atlas.fredhutch.org/nygc/multimodal-pbmc/ | 8,854 | √ | √ | |
| | P8_0 | CITE-seq | https://atlas.fredhutch.org/nygc/multimodal-pbmc/ | 8,908 | √ | √ | |
| BMMC mosaic dataset | ICA | 10X v2 Sequencing | https://www.dropbox.com/s/xe5tithw1xjxrfs/ica_bone_marrow.h5?dl=0 | 34,592 | √ | | |
| | ASAP | ASAP-seq | GSE156477 | 10,671 | | √ | √ |
| | CITE | CITE-seq | GSE128639 | 26,286 | √ | √ | |

Supplementary Table 2. Construction of rectangular and mosaic datasets with trimodal data.

| **Data** | **Batch** | **Rectangular dataset** | | | **Mosaic dataset** | | | | | | | | | | | | | | | | | | | | |
|---|---|---|---|---|---|---|---|---|---|---|---|---|---|---|---|---|---|---|---|---|---|---|---|---|---|
| | | full | | | paired+full | | | paired-abc | | | paired-ab | | | paired-ac | | | paired-bc | | | diagonal+full | | | diagonal | | |
| | | ATAC | RNA | ADT | ATAC | RNA | ADT | ATAC | RNA | ADT | ATAC | RNA | ADT | ATAC | RNA | ADT | ATAC | RNA | ADT | ATAC | RNA | ADT | ATAC | RNA | ADT |
| DOGMA | LLL Ctrl | √ | √ | √ | √ | √ | | √ | √ | | √ | √ | | √ | √ | | √ | | √ | √ | | | √ | | |
| | LLL Stim | √ | √ | √ | √ | | √ | | √ | √ | √ | | √ | | √ | √ | | √ | √ | | √ | | | √ | |
| | DIG Ctrl | √ | √ | √ | | √ | √ | √ | | √ | √ | √ | | √ | √ | | √ | | √ | | | √ | | | √ |
| | DIG Stim | √ | √ | √ | √ | √ | √ | | √ | √ | √ | | √ | | √ | √ | | √ | √ | √ | √ | √ | | | √ |
| TEADOG | W1 | √ | √ | √ | √ | √ | | √ | √ | | √ | √ | | √ | √ | | √ | | √ | √ | | | √ | | |
| | W6 | √ | √ | √ | √ | | √ | | √ | √ | √ | | √ | | √ | √ | | √ | √ | | √ | | | √ | |
| | LLL ctrl | √ | √ | √ | | √ | √ | √ | | √ | √ | √ | | √ | √ | | √ | | √ | | | √ | | | √ |
| | DIG stim | √ | √ | √ | √ | √ | √ | | √ | √ | √ | | √ | | √ | √ | | √ | √ | √ | √ | √ | | | √ |

Supplementary Table 3. An overview of compared methods for single-cell multimodal integration.

| **Method** | | | **Trimodal task (ATAC+RNA+ADT)** | | | | | | | | **Bimodal task (ATAC+RNA)** | | | | | **Bimodal task (RNA+ADT)** | | | | |
|---|---|---|---|---|---|---|---|---|---|---|---|---|---|---|---|---|---|---|---|---|
| **Integration type** | | **Name** | **full** | **paired +full** | **paired -abc** | **paired -ab** | **paired -ac** | **paired -bc** | **diag. +full** | **diag.** | **paired** | **atac+ paired** | **rna+ paired** | **diag.+ paired** | **diag.** | **paired** | **rna+ paired** | **adt+ paired** | **diag.+ paired** | **diag.** |
| Trimodal (ATAC+RNA+ ADT) | Mosaic | MIDAS | √ | √ | √ | √ | √ | √ | √ | √ | √ | √ | √ | √ | √ | √ | √ | √ | √ | √ |
| | | Multigrate | √ | √ | √ | √ | √ | √ | √ | √ | | | | | | | | | | |
| | | scMoMaT | √ | √ | √ | √ | √ | √ | √ | √ | | | | | | | | | | |
| | | scVAEIT | √ | √ | √ | √ | √ | √ | √ | √ | | | | | | | | | | |
| | | StabMap | √ | √ | √ | √ | √ | √ | √ | √ | | | | | | | | | | |
| | Rectangular | BBKNN+average | √ | | | | | | | | | | | | | | | | | |
| | | Harmony+WNN | √ | | | | | | | | | | | | | | | | | |
| | | LIGER+WNN | √ | | | | | | | | | | | | | | | | | |
| | | MOFA+ | √ | | | | | | | | | | | | | | | | | |
| | | PCA+WNN | √ | | | | | | | | | | | | | | | | | |
| | | Scanorama-embed+WNN | √ | | | | | | | | | | | | | | | | | |
| | | Scanorama-feat+WNN | √ | | | | | | | | | | | | | | | | | |
| | | Seurat-CCA+WNN | √ | | | | | | | | | | | | | | | | | |
| | | Seurat-RPCA+WNN | √ | | | | | | | | | | | | | | | | | |
| Bimodal (ATAC+RNA) | Mosaic | Cobolt | | | | | | | | | √ | √ | √ | √ | | | | | | |
| | | MultiVI | | | | | | | | | √ | √ | √ | √ | | | | | | |
| | Diagonal | uniPort | | | | | | | | | √ | | | | √ | | | | | |
| | | GLUE | | | | | | | | | | | | | √ | | | | | |
| Bimodal (RNA+ADT) | Mosaic | totalVI | | | | | | | | | | | | | | √ | √ | | | |
| | Rectangular | sciPENN | | | | | | | | | | | | | | √ | | | | |

Supplementary Table 4. Metrics of scMIB.

| | **Batch correction** | **Madality alignment** | **Biological conservation** |
|---|---|---|---|
| **Feature space** | Graph iLISI (feat)<br>Graph connectivity (feat)<br>kBET (feat) | ATAC AUROC<br>RNA Pearson's *r*<br>ADT Pearson's *r* | NMI (feat)<br>ARI (feat)<br>Graph cLISI (feat)<br>Isolated label F1 (feat) |
| **Embedding space** | Graph iLISI (embed)<br>Graph connectivity (embed)<br>kBET (embed) | Modality ASW<br>FOSCTTM<br>Label transfer F1 | NMI (embed)<br>ARI (embed)<br>Isolated label F1 (embed)<br>Graph cLISI (embed) |

Supplementary Table 5. Construction of cell type missing datasets.

| **Original data** | | | | | | | | |
|---|---|---|---|---|---|---|---|---|
| | **B** | **CD4 T** | **CD8 T** | **DC** | **Mono** | **NK** | **other** | **other T** |
| **LLL Ctrl** | 296 | 4983 | 1435 | 43 | 53 | 166 | 21 | 364 |
| **LLL Stim** | 159 | 3974 | 1080 | 45 | 11 | 213 | 27 | 388 |
| **DIG Ctrl** | 463 | 6913 | 1713 | 57 | 112 | 298 | 23 | 611 |
| **DIG Stim** | 332 | 6377 | 1608 | 62 | 64 | 431 | 33 | 620 |
| **Major cell type missing** | | | | | | | | |
| | **B** | **CD4 T** | **CD8 T** | **DC** | **Mono** | **NK** | **other** | **other T** |
| **LLL Ctrl** | 296 | 4983 | 1435 | 43 | 53 | 166 | 21 | **0** |
| **LLL Stim** | **0** | 3974 | 1080 | 45 | 11 | 213 | 27 | 388 |
| **DIG Ctrl** | 463 | **0** | 1713 | 57 | 112 | 298 | 23 | 611 |
| **DIG Stim** | 332 | 6377 | **0** | 62 | 64 | 431 | 33 | 620 |
| **Minor cell type missing** | | | | | | | | |
| | **B** | **CD4 T** | **CD8 T** | **DC** | **Mono** | **NK** | **other** | **other T** |
| **LLL Ctrl** | 296 | 4983 | 1435 | 43 | 53 | 166 | 21 | **0** |
| **LLL Stim** | 159 | 3974 | **0** | 45 | 11 | 213 | 27 | 388 |
| **DIG Ctrl** | 463 | 6913 | 1713 | 57 | 112 | **0** | 23 | 611 |
| **DIG Stim** | **0** | 6377 | 1608 | 62 | 64 | 431 | 33 | 620 |

*Cell numbers of each batch belonging different cell types are presented.

Supplementary Table 6. Construction of datasets with different sequencing depth.

| | Original | D_0.5 | D_0.2 | D_0.1 | D_0.05 | D_0.02 |
|---|---|---|---|---|---|---|
| **LLL Ctrl** | 1 | 0.5 | 0.2 | 0.1 | 0.05 | 0.02 |
| **LLL Stim** | 1 | 0.5 | 0.2 | 0.1 | 0.05 | 0.02 |
| **DIG Ctrl** | 1 | 1 | 1 | 1 | 1 | 1 |
| **DIG Stim** | 1 | 1 | 1 | 1 | 1 | 1 |

*Sampling ratios of each batch for different constructed datasets (D_0.5, D_0.2, D_0.1, D_0.05, D_0.02) are presented.

*Each modality of the original mosaic datasets was downsampled using the downsampleMatrix function (bycol = F) from the scuttle package in R.

Supplementary Table 7. Construction of rectangular and mosaic datasets with ATAC+RNA bimodal data.

| **Data (ATAC+RNA)** | **Batch** | **Rectangular dataset** | | **Mosaic dataset** | | | | | | | |
|---|---|---|---|---|---|---|---|---|---|---|---|
| | | paired | | rna+paired | | atac+paired | | diagonal+paired | | diagonal | |
| | | RNA | ATAC | RNA | ATAC | RNA | ATAC | RNA | ATAC | RNA | ATAC |
| DOGMA | LLL Ctrl | √ | √ | √ | | | √ | √ | | √ | |
| | LLL Stim | √ | √ | √ | √ | √ | √ | | √ | | √ |
| | DIG Ctrl | √ | √ | √ | | | √ | √ | | √ | |
| | DIG Stim | √ | √ | √ | √ | √ | √ | √ | √ | | √ |
| TEADOG | W1 | √ | √ | √ | | | √ | √ | | √ | |
| | W6 | √ | √ | √ | √ | √ | √ | | √ | | √ |
| | LLL ctrl | √ | √ | √ | | | √ | √ | | √ | |
| | DIG stim | √ | √ | √ | √ | √ | √ | √ | √ | | √ |

Supplementary Table 8. Construction of rectangular and mosaic datasets with RNA+ADT bimodal data.

| **Data (RNA+ADT)** | **Batch** | **Rectangular dataset** | | **Mosaic dataset** | | | | | | | |
|---|---|---|---|---|---|---|---|---|---|---|---|
| | | paired | | rna+paired | | adt+paired | | diagonal+paired | | diagonal | |
| | | RNA | ADT | RNA | ADT | RNA | ADT | RNA | ADT | RNA | ADT |
| DOGMA | LLL Ctrl | √ | √ | √ | | | √ | √ | | √ | |
| | LLL Stim | √ | √ | √ | √ | √ | √ | | √ | | √ |
| | DIG Ctrl | √ | √ | √ | | | √ | √ | | √ | |
| | DIG Stim | √ | √ | √ | √ | √ | √ | √ | √ | | √ |
| TEADOG | W1 | √ | √ | √ | | | √ | √ | | √ | |
| | W6 | √ | √ | √ | √ | √ | √ | | √ | | √ |
| | LLL ctrl | √ | √ | √ | | | √ | √ | | √ | |
| | DIG stim | √ | √ | √ | √ | √ | √ | √ | √ | | √ |

Supplementary Table 9. Datasets used in atlas and knowledge transfer.

| **Dataset** | **Batch** | **ATAC** | **RNA** | **ADT** | **Cell number** | **Atlas** | **Knowledge transfer** | |
|---|---|---|---|---|---|---|---|---|
| | | | | | | | **Reference** | **Query** |
| DOGMA dataset | LLL Ctrl | √ | √ | √ | 7,361 | √ | | √ |
| | LLL Stim | √ | √ | √ | 5,897 | √ | | √ |
| | DIG Ctrl | √ | √ | √ | 10,190 | √ | | √ |
| | DIG Stim | √ | √ | √ | 9,527 | √ | | √ |
| TEA dataset | W1 | √ | √ | √ | 7,325 | √ | √ | |
| | W3 | √ | √ | √ | 6,587 | √ | √ | |
| | W4 | √ | √ | √ | 6,897 | √ | √ | |
| | W5 | √ | √ | √ | 6,910 | √ | √ | |
| | W6 | √ | √ | √ | 7,137 | √ | √ | |
| TEA Multiome dataset | W1 | √ | √ | | 6,096 | √ | √ | |
| | W2 | √ | √ | | 7,284 | √ | √ | |
| 10X Multiome dataset | ChrX | √ | √ | | 9,868 | √ | √ | |
| | ChrC | √ | √ | | 9,582 | √ | √ | |
| | ARC2 10K | √ | √ | | 11,116 | √ | √ | |
| | ARC2 3K | √ | √ | | 2,566 | √ | √ | |
| ASAP dataset | Ctrl | √ | | √ | 4,255 | √ | √ | |
| | Stim | √ | | √ | 5,241 | √ | √ | |
| ASAP CITE dataset | Ctrl | | √ | √ | 5,086 | √ | √ | |
| | Stim | | √ | √ | 3,629 | √ | √ | |
| WNN CITE dataset | P1_0 | | √ | √ | 6,378 | √ | √ | |
| | P2_0 | | √ | √ | 5,899 | √ | √ | |
| | P3_0 | | √ | √ | 4,628 | √ | √ | |
| | P4_0 | | √ | √ | 5,285 | √ | √ | |
| | P5_0 | | √ | √ | 6,952 | √ | √ | |
| | P6_0 | | √ | √ | 6,060 | √ | √ | |
| | P7_0 | | √ | √ | 8,854 | √ | √ | |
| | P8_0 | | √ | √ | 8,908 | √ | √ | |
| Total cells | | 123,839 | 176,022 | 139,006 | 185,518 | 185,518 | 152,543 | 32,975 |

Supplementary Table 10. Construction of 14 dogma mosaic query datasets for knowledge transfer.

| Batch | Dogma query datasets for knowledge transfer | | | | | | | | | | | | | | | | | | | | |
|---|---|---|---|---|---|---|---|---|---|---|---|---|---|---|---|---|---|---|---|---|---|
| | full | | | paired+full | | | paired-abc | | | paired-ab | | | paired-ac | | | paired-bc | | | diagonal+full | | |
| | ATAC | RNA | ADT | ATAC | RNA | ADT | ATAC | RNA | ADT | ATAC | RNA | ADT | ATAC | RNA | ADT | ATAC | RNA | ADT | ATAC | RNA | ADT |
| LLL Ctrl | √ | √ | √ | √ | √ | | √ | √ | | √ | √ | | √ | √ | | √ | | √ | √ | | |
| LLL Stim | √ | √ | √ | √ | | √ | | √ | √ | √ | | √ | | √ | √ | | √ | √ | | √ | |
| DIG Ctrl | √ | √ | √ | | √ | √ | √ | | √ | √ | √ | | √ | √ | | √ | | √ | | | √ |
| DIG Stim | √ | √ | √ | √ | √ | √ | | √ | √ | √ | | √ | | √ | √ | | √ | √ | √ | √ | √ |

| Batch | diagonal | | | paired-a | | | paired-b | | | paired-c | | | atac | | | rna | | | adt | | |
|---|---|---|---|---|---|---|---|---|---|---|---|---|---|---|---|---|---|---|---|---|---|
| | ATAC | RNA | ADT | ATAC | RNA | ADT | ATAC | RNA | ADT | ATAC | RNA | ADT | ATAC | RNA | ADT | ATAC | RNA | ADT | ATAC | RNA | ADT |
| LLL Ctrl | √ | | | √ | √ | | √ | | √ | | √ | √ | √ | | | | √ | | | | √ |
| LLL Stim | | √ | | √ | √ | | √ | | √ | | √ | √ | √ | | | | √ | | | | √ |
| DIG Ctrl | | | √ | √ | √ | | √ | | √ | | √ | √ | √ | | | | √ | | | | √ |
| DIG Stim | | | √ | √ | √ | | √ | | √ | | √ | √ | √ | | | | √ | | | | √ |

Supplementary Table 11. Functionality comparison of MIDAS and other mosaic integration methods.

| Method | Diverse mosaic combination | Embedding space | | Feature space | | Knowledge transfer | Flexible knowledge transfer | General mosaic integration |
|---|---|---|---|---|---|---|---|---|
| | | Batch correction | Modality alignment | Imputation | Batch correction | | | |
| Multigrate | √ | √ | | √ | | √ | | |
| scMoMaT | √ | | | √ | | | | |
| scVAEIT | √ | | | √ | | √ | | |
| StabMap | | | | √ | | | | |
| MIDAS | √ | √ | √ | √ | √ | √ | √ | √ |

Supplementary Table 12. The layer sizes of each module in MIDAS.

<table>
<tr><th colspan="2">Module</th><th colspan="3">Layer size</th></tr>
<tr><td rowspan="4">Encoder</td><td>$f^{ATAC}$</td><td>$D^{chr1}$→128→32<br>⋯<br>$D^{chr22}$→128→32</td><td>32×22→1024</td><td rowspan="3">1024→128→68<br>(shared layers)</td></tr>
<tr><td>$f^{RNA}$</td><td colspan="2">$D^{RNA}$→1024</td></tr>
<tr><td>$f^{ADT}$</td><td colspan="2">$D^{ADT}$→1024</td></tr>
<tr><td>$f^{s}$</td><td colspan="3">$D^{s}$→16→16→68</td></tr>
<tr><td rowspan="4">Decoder</td><td>$g^{ATAC}$</td><td rowspan="3">34→128→1024<br>(shared layers)</td><td>1024→32×22</td><td>32→128→$D^{chr1}$<br>⋯<br>32→128→$D^{chr22}$</td></tr>
<tr><td>$g^{RNA}$</td><td colspan="2">1024→$D^{RNA}$</td></tr>
<tr><td>$g^{ADT}$</td><td colspan="2">1024→$D^{ADT}$</td></tr>
<tr><td>$g^{s}$</td><td colspan="3">2→16→16→$D^{s}$</td></tr>
<tr><td rowspan="4">Classifier</td><td>$r$</td><td colspan="3">32→128→64→$D^{s}$</td></tr>
<tr><td>$r^{ATAC}$</td><td colspan="3">32→128→64→$D^{s}$</td></tr>
<tr><td>$r^{RNA}$</td><td colspan="3">32→128→64→$D^{s}$</td></tr>
<tr><td>$r^{ADT}$</td><td colspan="3">32→128→64→$D^{s}$</td></tr>
</table>

Supplementary Table 13. The input layer sizes of MIDAS on different tasks.

| **Input size** | **Dataset** | | | | |
|---|---|---|---|---|---|
| | **dogma-full** | **teadog-full** | **atlas-no_dogma** | **atlas** | **bm** |
| $D^{ATAC}$ | 30,589 | 31,243 | 52,393 | 52,867 | 37,840 |
| - $D^{chr1}$ | 3,165 | 3,210 | 5,318 | 5,361 | 3,692 |
| - $D^{chr2}$ | 2,338 | 2,315 | 4,124 | 4,160 | 3,050 |
| - $D^{chr3}$ | 1,764 | 1,748 | 3,234 | 3,270 | 2,473 |
| - $D^{chr4}$ | 993 | 985 | 1,861 | 1,875 | 1,494 |
| - $D^{chr5}$ | 1,407 | 1,370 | 2,503 | 2,535 | 1,917 |
| - $D^{chr6}$ | 1,853 | 1,861 | 3,293 | 3,312 | 2,421 |
| - $D^{chr7}$ | 1,466 | 1,473 | 2,552 | 2,591 | 1,894 |
| - $D^{chr8}$ | 1,169 | 1,175 | 2,032 | 2,066 | 1,589 |
| - $D^{chr9}$ | 1,211 | 1,258 | 2,106 | 2,131 | 1,544 |
| - $D^{chr10}$ | 1,418 | 1,437 | 2,473 | 2,500 | 1,861 |
| - $D^{chr11}$ | 1,600 | 1,670 | 2,761 | 2,774 | 2,019 |
| - $D^{chr12}$ | 1,656 | 1,685 | 2,877 | 2,896 | 2,087 |
| - $D^{chr13}$ | 620 | 606 | 1,120 | 1,127 | 800 |
| - $D^{chr14}$ | 1,147 | 1,141 | 1,892 | 1,905 | 1,262 |
| - $D^{chr15}$ | 1,026 | 1,042 | 1,692 | 1,708 | 1,248 |
| - $D^{chr16}$ | 1,317 | 1,384 | 2,063 | 2,088 | 1,447 |
| - $D^{chr17}$ | 1,929 | 2,073 | 3,142 | 3,162 | 2,024 |
| - $D^{chr18}$ | 498 | 497 | 917 | 928 | 705 |
| - $D^{chr19}$ | 1,986 | 2,176 | 2,980 | 3,001 | 1,988 |
| - $D^{chr20}$ | 866 | 933 | 1,519 | 1,525 | 1,079 |
| - $D^{chr21}$ | 391 | 379 | 644 | 654 | 437 |
| - $D^{chr22}$ | 769 | 825 | 1,290 | 1,298 | 809 |
| $D^{RNA}$ | 4,045 | 4,047 | 4,051 | 4,047 | 4,096 |
| $D^{ADT}$ | 208 | 213 | 228 | 228 | 239 |
| $D^{s}$ | 4 | 4 | 23 | 27 | 3 |

Supplementary Table 14. Hyperparameter settings for MIDAS training.

| Hyperparameter | Value |
| --- | --- |
| Modality alignment loss weight $\alpha$ | 50 |
| Technical IB loss weight $\beta^s$ | 30 |
| Biological IB loss weight $\beta^x$ | 4 |
| Technical noise likelihood loss weight $\gamma$ | 1000 |
| Classifier update number $K$ | 3 |
| Training-validation ratio | 95:5 |
| Mini-batch size | 256 |
| Maximum number of training epochs | 2000 |
| Learning rate | 0.0001 |
| Hidden layer dropout rate | 0.2 |
